# scINTILLA: Single-Cell Integrated Inference, Labelling, and Landscape Analysis for Assessing the Quality and Hidden Diversity of Cell Annotations

**DOI:** 10.64898/2026.07.27.740477

**Authors:** Sina Kanannejad, Noemi Bongiorni, Elisa Nordera, Sara Redaelli, Irene Rusconi, Rachele Zanin, Alice Giustacchini, Soumick Chatterjee

**Author notes:** These authors contributed equally: Noemi Bongiorni, Elisa Nordera, Sara Redaelli, Irene Rusconi, Rachele Zanin.

## Abstract

Single-cell RNA sequencing has made it possible to build atlases of millions of cells, but the cell-type labels in these atlases are rarely checked once they are assigned. When a label covers a transcriptionally heterogeneous population, every analysis that depends on it inherits the problem, and automated label transfer in particular becomes unreliable. We present scINTILLA (Single-Cell Integrated Inference, Labelling, and Landscape Analysis), a framework that scores every label in an annotated dataset from three complementary signals: clustering confusion (neighbours that share a cluster but carry another label), label fragmentation (one label spread over several clusters) and supervised consistency (agreement, confidence and entropy of a classifier ensemble). The three signals are combined into a per-label quality score, scINTILLA-F, which can optionally be rank-fused with the Silhouette width. To test whether the score finds real annotation problems, we injected three kinds of known errors (merged labels, contaminated labels and over-split labels) into five Human Cell Atlas datasets, giving 150 perturbed annotations, and compared scINTILLA with the Silhouette baseline, kNN purity, SCCAF and ROGUE in a pre-registered analysis. The four best of the eleven methods that score every label were all scINTILLA-based. The fusion of scINTILLA-F with Silhouette ranked first (balanced AUROC 0.908; Silhouette 0.896, kNN purity 0.815, SCCAF 0.770) and improved on the Silhouette baseline in every atlas, and adding fragmentation raised the detection of merged labels in every atlas. scINTILLA and the Silhouette baseline were complementary. The Silhouette baseline was better at finding merges, whereas scINTILLA was better at finding oversplit labels. Applied to the real atlases, scINTILLA flagged the “Unknown” and monocytederived macrophage labels of the lung atlas and the “stem cell” label of the endoderm organoid atlas. Re-analysis of these populations resolved disease-associated monocyte and macrophage states and a set of progenitor, stromal and differentiated identities with distinct tissue and protocol origins. Finally, we tested the score on merges that arise naturally, in a leukaemia atlas of 183 patients whose blast populations we had annotated. Label fragmentation revealed the diversity of the blasts hidden under a patient label: in acute myeloid leukaemia it was higher in patients whose blasts formed several populations. In each of these cases scINTILLA pointed to the labels that needed a closer look, and the interpretation remained with the expert. It is therefore advisory: it ranks labels for expert review and shows which signal flagged each one.

## 1 Introduction

Single-cell RNA sequencing (scRNA-seq) profiles the transcriptomes of individual cells, and can do so for millions of cells at once [42]. Bulk measurements average the signal from distinct cell populations and so obscure them; scRNA-seq characterises each cell on its own terms. It is now used widely in developmental biology, immunology and oncology, because it captures the range of cell states present in a tissue at the moment of sampling.

International consortia have set out to catalogue the cell types and cell states of the whole human body. The best known, the Human Cell Atlas (HCA), aims to build reference maps of every cell type in every tissue and organ, at single-cell resolution and across individuals, developmental stages and disease contexts [52, 55]. Datasets of tens of millions of cells from hundreds of donors are now routine. The atlases are meant to record the cellular composition of healthy tissues and the molecular identity of their cell types and states, so that physiological variation and disease-associated alterations can be measured against them [58, 14].

The scale that makes these atlases valuable also makes them hard to annotate. In practice, atlas construction pipelines typically involve integrating cells across datasets, embedding them into shared latent spaces in which technical batch effects are removed, clustering in transcriptomic space, inspecting marker gene expression within each cluster, and manually curating labels on the basis of expert knowledge and the published literature [41, 9]. When this process is applied to hundreds of thousands or millions of cells spanning multiple tissues, donors and experimental batches, the rigour that smaller studies can afford is no longer practical. Labels may then be inconsistent across datasets, too coarse, or simply wrong, particularly for cell types that lack wellestablished canonical markers, that exist in rare proportions, or that occupy transcriptional states intermediate between two recognised identities. Annotation is also increasingly automated. Labels are assigned by classifiers trained on annotated data, such as CellTypist [14] and scANVI [70], or by reference-based methods that compare a new dataset with an annotated reference or map it into one, such as SingleR [1] and Symphony [32]. These methods carry the labels of the reference, together with any errors in them, into every dataset they annotate.

Yet the atlas analysis toolkit does little to help assess cell types, or decide which to look at first, by how internally consistent their labels are. A cluster-level label assumes that the cells sharing it are transcriptionally coherent and correspond to a meaningful biological entity. That assumption often fails in heterogeneous tissues and in cell types whose biology is incompletely characterised. A label that covers a heterogeneous population is not always an error. It may be an umbrella over several distinct cell types or states [6], and these are often the populations of most interest, because discovering such diversity is a main reason for sequencing an atlas. In either case the label deserves a closer look. Such a label can also mislead tools that train classifiers to transfer cell type labels from reference to query datasets [32, 63, 71]. These tools assume that the cells bearing a given label form a coherent, learnable category; when the training labels cover heterogeneous populations, classifier performance degrades, label transfer becomes unreliable and the errors propagate into downstream analyses.

Machine learning has contributed substantially to cell type classification and label transfer in single-cell genomics, and these tools are now standard parts of atlas workflows. But they take the labels as given. A label is ground truth to be propagated, and its own quality is not something these tools assess. General-purpose measures of cluster coherence do exist, among them the Silhouette width [54], the self-projection accuracy of SCCAF [45] and the entropy-based purity score ROGUE [39], but each captures a single notion of coherence, none indicates which kind of error is likely behind a low score, and, as we show below, most are weak for at least one common type of annotation error. Other recent tools go after over- and under-clustering by testing or rebuilding the clusters themselves. scSHC asks whether two clusters are statistically distinct [22]; CHOIR runs random-forest permutation tests down a cluster hierarchy [57]; scTriangulate chooses among competing clusterings by reclassification stability and marker specificity [37]. Outside single-cell biology, confident learning estimates label errors from out-of-sample predicted probabilities [50]. All four decide how a dataset should be partitioned, or which cells are mislabelled. None takes an existing label set and ranks its labels, which is the task here. Users therefore have no systematic way to tell where an analysis that depends on the annotation can be trusted.

We developed scINTILLA – Single-Cell Integrated Inference, Labelling, and Landscape Analysis – a frame-work that combines supervised and unsupervised machine learning to score the quality of every cell-type label in a single-cell dataset. Each label is scored from three signals that fail in different ways: clustering confusion, label fragmentation and supervised consistency. Because no ground truth exists for the quality of a real annotation, we built a benchmark in which known errors are injected into real atlases, and used it to compare scINTILLA with established label- and cluster-quality measures. We then applied the benchmarked configuration to five Human Cell Atlas datasets and followed up two flagged populations in detail. Finally, we asked whether the score responds to merges that occur naturally and not by injection: in an atlas of patients with acute leukaemia, a patient label covers one or several blast populations, and we tested whether the scores follow an annotation of that heterogeneity. An overview of the framework and the benchmark is given in Fig. 1.

**Fig. 1:**
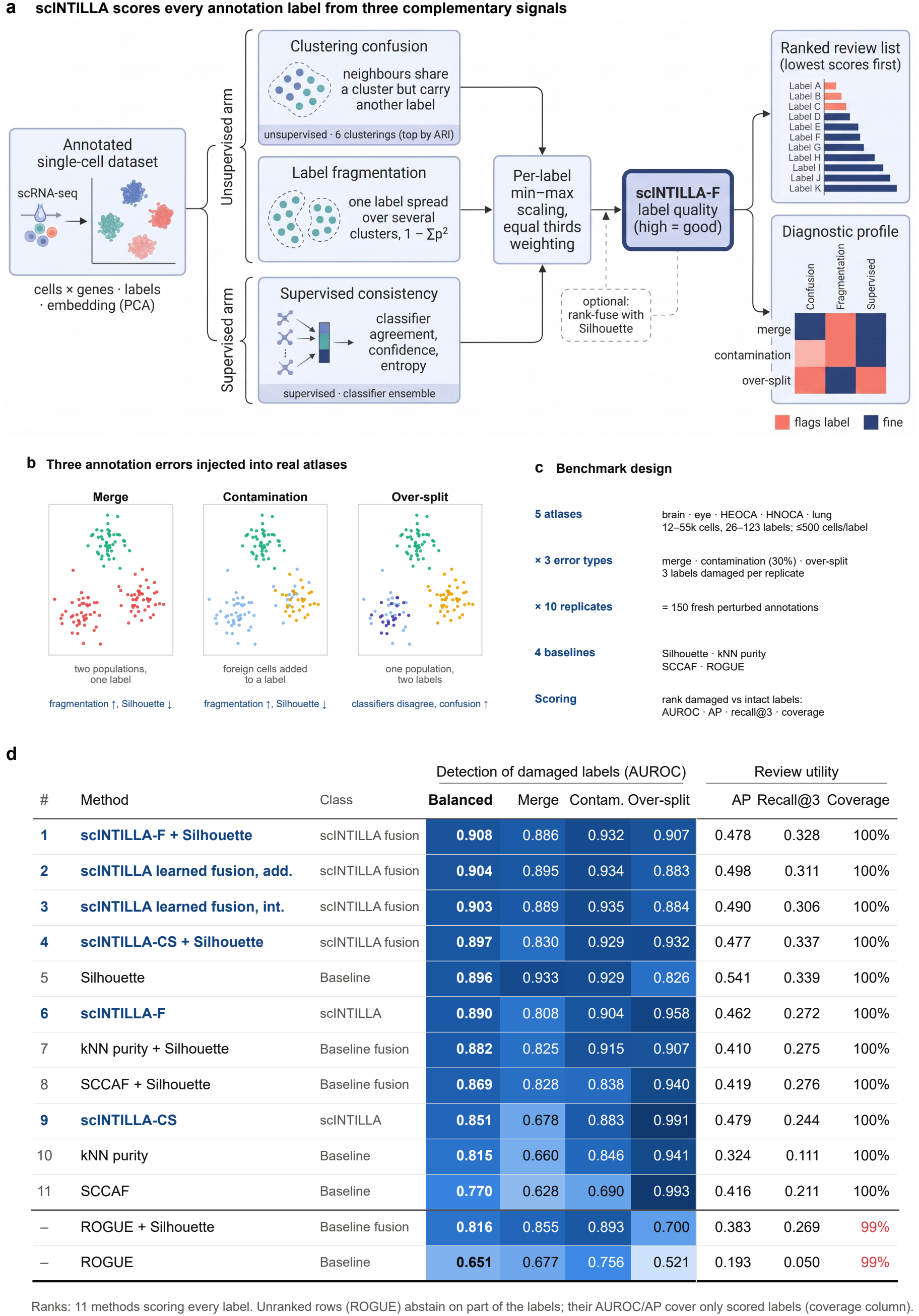
The scINTILLA framework and its benchmark. **(a)** scINTILLA scores every annotation label from three complementary signals computed on an annotated dataset (cells× genes, labels and an embedding): clustering confusion (a cell s neighbours share its cluster but carry another label), label fragmentation (one label spread over several clusters, 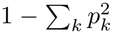) and supervised consistency (classifier agreement, confidence and entropy). Confusion and fragmentation are computed for the six clusterings that best match the annotation (the six highestARI partitions over six clustering algorithms and their parameter settings), and supervised consistency is computed from a classifier ensemble. Per-label means are min-max scaled and weighted so that each signal contributes one third, giving scINTILLA-F (high = good). Labels are returned as a ranked review list, lowest scores first, together with a diagnostic profile showing which signal failed. scINTILLA-F can optionally be rank-fused with the Silhouette width. Created in https://BioRender.com **(b)** The three annotation errors injected into real atlases. Merge: two populations given one label (fragmentation rises, Silhouette falls). Contamination: foreign cells added to a label (fragmentation rises, Silhouette falls). Over-split: one population given two labels (classifiers disagree, confusion rises). **(c)** Benchmark design (pre-registered): five atlases (12-55 thousand cells and 26-123 labels each, at most 500 cells per label) ×three error types× ten replicates, giving 150 freshly perturbed annotations, four baselines, and damaged and intact labels ranked by each method and scored by AUROC, average precision (AP), recall@3 and coverage. **(d)** Detection of damaged labels. AUROC is the median over the ten replicates of each atlas and error type; “Balanced” is the mean over error types and atlases. Recall@3 is the fraction of all damaged labels that appear among the three lowest-scoring labels (at most 0.5 for over-splits, which have six), and coverage is the fraction of labels a method scores. scINTILLA-CS denotes the score without fragmentation (clustering confusion and supervised consistency only), shown as an ablation; the learned fusions and the “scINTILLA-CS + Silhouette” row are built on that score. Ranks are given for the eleven methods that score every label; ROGUE abstains on 1% of labels, so its AUROC and AP cover only the labels it scores.

## 2 Methods and models

### 2.1 Overview

scINTILLA is distributed as a general pipeline for singlecell analysis, and the label-quality score is built on two of its modules: the clustering benchmark and the classification benchmark. This section summarises those two modules and the methods that were run in this study, and then describes the score, the benchmark and the leukaemia analysis. The two modules are described in full in the Supplementary Methods (Sections S5 and S6). The parts of the pipeline that were not used here are described there as well (Sections S1–S10): exploratory data analysis, preprocessing and normalisation, feature selection, non-linear dimensionality reduction, spectral, consensus and patient-level clustering, SHAP-based feature importance, differential expression, cell-type annotation, batch correction and the statistical utilities. The tables that list the functions of the package with their inputs and outputs are given there as well (Supplementary Tables 7–13).

### 2.2 Clustering

The unsupervised signals of the score are computed from a benchmark of clustering methods (`benchmark_clustering_methods`), all run on the same embedding. Six families were run in this study. The K-means family [23] comprises standard K-means with `k-means++` and with random initialisation, spherical K-means [27], in which each cell vector is L2-normalised so that distances reflect angle and not magnitude, and bisecting K-means [62], which reaches the partition through successive binary splits. Agglomerative hierarchical clustering [49] is run over a grid of four linkages (complete, average, single and Ward) and seven distance metrics, with Ward restricted to the Euclidean metric. DBSCAN [17] is swept over a grid of neighbourhood radius and minimum neigh-bourhood size, and HDBSCAN [44] over a grid of minimum cluster size and `min_samples`; both leave cells that fall in no cluster unassigned, as noise. For HDBSCAN, each parameter combination is scored as the number of clusters minus the noise fraction, and only the highest-scoring partition enters the benchmark. This rule favours the finest partition in the grid whatever its noise, which is why HDBSCAN returned no partition with at most 50% of cells as noise for the lung and HNOCA atlases. Leiden [65] and Louvain [2] detect communities by optimising modularity on a *k*-nearest-neighbour graph of the embedding, and both sweep a resolution grid from 0.1 to 3.0.

Every partition is compared with the annotation by the Adjusted Rand Index (ARI) [28]. Cells labelled as noise are excluded from the ARI, and a configuration that labels more than 50% of the cells as noise receives no ARI, so that degenerate solutions are not rewarded. Density-based methods are thus scored on their core cells only, which favours them slightly. The benchmark results for the five atlases are shown in Supplementary Fig. 1. The package can also choose the Leiden resolution without the labels, by a stability criterion that picks the plateau of the Normalised Variation of Information curve along the resolution grid; this criterion was not used here.

The objective functions, options and implementations of these methods, the three methods of the package that were not run (spectral, consensus and patient-level clustering) and the function interfaces are given in Supplementary Methods, Section S5 (Supplementary Tables 10 and 11).

### 2.3 Classification

The supervised signal of the score is computed from an ensemble of classifiers trained to reproduce the annotation. Twelve classifiers were run in this study: linear and quadratic discriminant analysis [18]; a linear support vector machine [10] (one-vs-rest), which returns decision scores and not class probabilities; logistic regression [12]; Gaussian naive Bayes [53]; *k*-nearest neighbours [11] with *k* = 5; a multi-layer perceptron [56] with two hidden layers of 100 and 50 units; a random forest [5]; histogram-based gradient boosting [20]; XGBoost [8]; LightGBM [33]; and a stacking ensemble [69, 4], in which a logisticregression meta-model is fitted on the 5-fold out-of-fold predictions of a logistic regression, a random forest and a linear support vector machine. XGBoost and LightGBM use their own libraries, and the other classifiers use scikitlearn.

The classification benchmark (`benchmark_models_-comprehensive`) evaluates the classifiers by accuracy and by the macro-averaged F1-score (Supplementary Fig. 1). How their predictions enter the score is described in Section 2.4. The model definitions, the hyperparameters, the validation strategies of the benchmark and the function interfaces are given in Supplementary Methods, Section S6 (Supplementary Tables 12 and 13).

### 2.4 Label-quality scoring

scINTILLA works on an annotated dataset together with an embedding. In the analyses reported here, the embedding was a label-free principal component analysis (30 components) computed on 2,000 highly variable genes with Scanpy [68], and the classifiers were trained on the expression of the same genes.

#### Clustering confusion

The clustering benchmark (Section 2.2) is run over a range of parameters (methods that take a cluster count are given the number of annotated labels; Leiden and Louvain sweep resolutions, and DB-SCAN and HDBSCAN sweep parameter grids), and the six clusterings whose partitions best agree with the annotation (by ARI) are kept. The default preset runs six families (the K-means family, agglomerative hierarchical clustering, DBSCAN, HDBSCAN, Louvain and Leiden); spectral and consensus clustering are in the library but were not run (Supplementary Methods, Section S5). The six are chosen over all methods and parameter settings together, so near-identical partitions from neighbouring resolutions can take several of the six places. Cells that DBSCAN or HDBSCAN leave unassigned (label− 1) are treated as one cluster in both formulas below. Such partitions reached the top six in two atlases only: HEOCA (44% noise) and the brain (21%). For each of these clusterings and each cell *i*, the confusion score is the fraction of the cell s *K* = 50 nearest neighbours in the embedding that fall in the same cluster as *i* but carry a different label. With *y*_*i*_ the label of cell *i*, 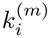 its cluster in the *m*-th selected clustering (*m* = 1, …, 6) and *N*_*K*_(*i*) its neighbours,

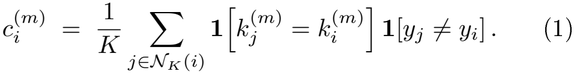

High confusion means that cells of the label are routinely grouped with cells of another label.

#### Label fragmentation

For a label *L* and a clustering *c*, let *p*_*L,k*_ be the fraction of the cells of *L* that fall in cluster *k*. Fragmentation is the Gini-Simpson index [60]

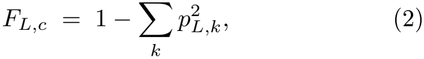

which is zero when a label sits entirely in one cluster and grows as the label spreads over several clusters. Confusion and fragmentation look at the same clusterings from opposite sides: confusion asks whether a label s cluster also holds other labels, and fragmentation asks whether a label is split across clusters.

#### Supervised consistency

A classifier ensemble is trained to reproduce the annotation. The classifiers are fitted on a stratified 80% of the cells, and predictions are then taken for every cell, training cells included. With *M* fitted classifiers, *ŷ*_*i,m*_ the class predicted for cell *i* by classifier *m*, and 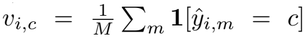 the share of votes for class *c*, three per-cell quantities are computed: prediction agreement *a*_*i*_ = max_*c*_ *v*_*i,c*_, prediction entropy *H*_*i*_ = − *∑*_*c*_ *v*_*i,c*_ log_2_ *v*_*i,c*_, and confidence 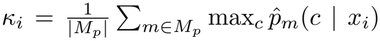, where *M*_*p*_ is the set of classifiers that return probabilities (all but the linear SVM). The default preset fits twelve classifiers, and all of them enter with equal weight.

#### Composite score (sclNTlLLA-F)

Each per-cell quantity is averaged within labels and min-max scaled across the labels of the dataset; confusion, fragmentation and entropy are inverted so that high always means good. The six confusion columns and the six fragmentation columns each receive weight 1, and the three supervised columns weight 2, so that each of the three signals contributes one third of the total. Writing 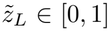, 1] for the scaled label mean of a per-cell quantity *z* (a quantity that is constant across labels is set to 0 before inversion, so it contributes 1 if it is negatively oriented and 0 otherwise), and *a, H* and *κ* for prediction agreement, entropy and confidence, the score of label *L* is

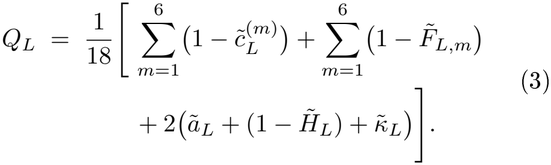

The same composite computed without the fragmentation columns (dropping the second sum and dividing by 12), from clustering confusion and supervised consistency alone, is denoted scINTILLA-CS and is used as an ablation. Because the score is scaled within a dataset, it is a ranking of labels inside one atlas and is not comparable between atlases.

#### Rank fusion with Silhouette

For a quality score *s* computed on *n* labels, the within-annotation percentile rank of label *L* is *r*_*s*_(*L*) = (rank_*s*_(*L*) −0.5)*/n*, with ties given their average rank. The fused score is the mean of the two percentile ranks,

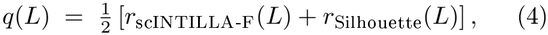

where the Silhouette of a label is the mean per-cell Silhouette width [54] of its cells on the shared embedding, with labels used as clusters. The same rule was applied to every baseline, so that any gain from fusion could be compared fairly.

### 2.5 Benchmark with injected annotation errors

#### Datasets and perturbations

From each of the five atlases (Section 3) we drew a pool of up to 1,000 random cells per label; highly variable genes and the PCA were computed on this pool without using the labels. Three error types were injected, each damaging three labels per replicate: *merge*, in which three disjoint pairs of labels were combined into single labels; *contamination*, in which cells from other labels, equal to 30% of the target label s size, were added to three labels (about 23% of the final label); and *over-split*, in which three labels were divided at random into two halves, giving six damaged child labels. Each atlas and error type was perturbed ten times, giving 150 annotations. A label was eligible for perturbation if it had at least 30 cells in the pool (60 for an over-split, so that each half kept 30). Labels were drawn uniformly at random without replacement; merge partners were paired at random, with no regard to similarity, and contaminating cells were drawn without replacement from the cells of untouched labels. Labels named “unknown” were never perturbed and were never counted as intact or damaged; their cells stayed in every scored dataset and were left out of the evaluation only. After perturbation, every label was capped at 500 random cells, as in the real-atlas analyses, so that damaged labels carried no size cue (12,147–54,672 cells and 26–123 labels per atlas). The two halves of an over-split label are smaller than their parent, so a small label can still carry such a cue after the cap; intact labels range from 30 to 500 cells as well.

#### Baselines

scINTILLA was compared with four labelquality measures computed on the same shared embedding: the mean Silhouette width of each label [54]; kNN purity, the mean fraction of a cell s 50 nearest neighbours that carry its label; the self-projection accuracy of SCCAF [45]; and the entropy-based purity score ROGUE [39], which abstains on labels with too few cells. Each baseline was also rank-fused with Silhouette using the rule above. Two learned fusions, logistic regressions of damage on the ranks of scINTILLA-CS and of Silhouette (with and without an interaction term), were fitted on the perturbations of an earlier, unregistered run of the benchmark (other random draws from the same five atlases) and frozen in the pre-registration, before the 150 benchmark annotations were evaluated. They use scINTILLA-CS because scINTILLA-F did not exist when they were fitted. With *s* and *g* the within-annotation percentile ranks of scINTILLA-CS and Silhouette, the frozen linear predictors of damage are

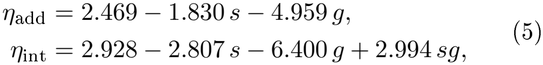

and the quality score is −*η*. Both models were fitted without penalty, with equal total weight for each error type and class, and are not calibrated probabilities.

#### Evaluation

Each method ranked the labels of a perturbed annotation from lowest to highest quality, and damaged labels were treated as positives. We report AU-ROC, average precision (AP), recall@3 (the fraction of all damaged labels that appear among the three lowest-scoring labels; precision@3 is the fraction of those three that are damaged) and coverage (the fraction of labels that receive a score). Values were summarised as the median over the ten replicates of each atlas and error type; “balanced” values are the mean over the three error types and five atlases. An over-split annotation contains six damaged labels, so its recall@3 cannot exceed 0.5. Paired comparisons between two methods were computed on labels scored by both. A paired effect is the median over replicates of the per-replicate difference between the two methods, aggregated in the same way; it is not the difference between their summary values, and the two can differ in the second decimal place. For paired effects, 95% intervals were obtained from 5,000 bootstrap resamples of whole atlases [15]. For the pre-specified headline contrasts, one-sided exact sign-flip tests over atlases were computed with Holm correction [26]. With only five atlases the smallest attainable *p* is 1*/*32 and no contrast survives the correction, so we report the intervals as exploratory effect sizes. Per-atlas values of every endpoint, additional endpoints (precision at 1, 3 and 10% of labels, standardised partial AUROC [43]) and leave-one-atlas-out estimates are given in Supplementary Figs. 11-13.

#### Pre-registration and sensitivity analyses

The analysis plan, predictions and decision rules were frozen in a version-controlled pre-registration [51] before any score was computed; 12 of 13 predictions were confirmed. One deviation was recorded: an import error in the ROGUE wrapper was fixed and ROGUE was rerun. Two sensitivity analyses used the same perturbation design. In the first, 20,000 cells were sampled uniformly from each atlas before perturbation instead of capping each label (20-61 labels per atlas). In the second, exploratory analysis, the PCA was batch-corrected with Harmony [34] (donor as batch, fitted without labels), and scINTILLA, Silhouette and kNN purity were rescored; SCCAF and ROGUE do not use the embedding. The real-atlas scoring was likewise repeated with Harmony (all atlases) and with the atlas-provided scVI space [40] (eye).

#### Scoring the real atlases

For the real-atlas analyses, at most 500 cells were sampled per label, and labels with fewer than 30 cells in the full atlas were excluded. Because scINTILLA-F is scaled within each atlas, atlases were compared on the unscaled per-label quantities (classifier disagreement with the label, confusion and fragmentation for the top clustering, and Silhouette), summarised by their median and interquartile range over labels. Classifier disagreement with the label is 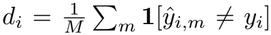, and consensus mismatch is **1**[arg max_*c*_ *v*_*i,c*_ *y*_*i*_]; unlike agreement, both refer to the assigned label *y*_*i*_. The scores were reproducible: the Spearman correlation with an independent run was 1.00 for the brain, lung, HEOCA and HNOCA atlases and 0.990 for the eye atlas (scINTILLA-F; 0.976–0.995 over the three scores) (Supplementary Fig. 15).

#### Re-annotation of flagged populations

Two flagged populations were re-analysed. In the lung atlas, the monocyte-derived macrophages, the labels their predictions were most often confused with in the initial scoring run, and the cells labelled “Unknown” (823,751 cells in all) were integrated with MrVI [3] and clustered with the Leiden algorithm on the MrVI neighbour graph (`scanpy.tl.leiden`, igraph implementation, two iterations, resolution 0.9). In this atlas “Unknown” marks cells that the original study mapped to its reference with low confidence [58]. The nine clusters containing monocytes and macrophages (493,585 cells) were kept, and each received a refined label from its marker genes, giving nine refined labels. In HEOCA, the “stem cell” label shared clusters with five differentiated epithelial labels in the top-ranked clustering, so all cells of the six labels were taken together, in order that re-clustering could separate genuine progenitors from differentiated cells. This stem-cell group (373,373 cells: stem cell 211,159, epithelial cell 63,324, goblet cell 44,884, enterocyte 31,239, colon epithelial cell 14,094 and M cell of gut 8,673) was integrated with MrVI and clustered in the same way at resolution 0.7; the 19 resulting clusters were merged into 13 refined labels. Marker genes were ranked with `scanpy.tl.rank_genes_groups` [68] on library-sizenormalised, log-transformed expression. Enrichment of each refined label in clinical conditions (lung) or tissues of origin (HEOCA) was tested with Fisher s exact test for every condition-label or tissue-label pair, with Benjamini–Hochberg correction (FDR < 0.005 for the lung and < 0.05 for HEOCA). These tests treat cells as independent and are descriptive. To check that the enrichments did not reflect pseudoreplication, we repeated them with donors as units. A unit was one donor within one condition (lung) or tissue (HEOCA), and units with fewer than 10 cells in the compartment were excluded. For each unit we computed the share of its cells in every refined label, and for each label–group pair that was significant at cell level we compared the shares inside and outside the group with a two-sided Mann–Whitney U test and with a quasi-binomial generalised linear model, both with Benjamini–Hochberg correction. An enrichment was taken as supported when its FDR was below 0.05, its direction matched the cell-level result and at least three donors in the group carried the label. In the lung the three COVID-19 categories were merged and the generic “ILD” category (2,218 cells) was removed (Supplementary Table 5; Supplementary Fig. 10). To re-score the refined labels, cells were sampled stratified by refined label (at most 1,000 per label, seed 0; 9,000 lung and 13,000 HEOCA cells), and the original and refined labellings were scored on the identical cells with the benchmarked configuration. Because composite scores are scaled within a labelling, only the unscaled, cell-weighted metrics were compared (Supplementary Table 4). As a sensitivity analysis, the rescoring was repeated after correcting the PCA with Harmony (donor as batch, labels not used); cells, genes and classifier inputs were unchanged, and only clustering, confusion, fragmentation and Silhouette used the corrected embedding.

### 2.6 Benchmark against annotated blast heterogeneity in leukaemia

The injected errors of Section 2.5 are synthetic. As a complementary test we used a setting in which labels of unequal heterogeneity occur naturally. In single-cell data from patients with acute leukaemia, the leukaemic cells (blasts) of one patient may form one population or several, so a label that names only the patient is, for some patients, a merge of several populations. No ground truth exists for this heterogeneity, so the analysis has two parts: a reference annotation of blast heterogeneity, and a test of whether scINTILLA scores follow it.

#### Atlas

We used diagnosis samples from 184 patients and ten healthy controls (1,296,005 cells): 84 patients with B-cell acute lymphoblastic leukaemia (B-ALL) [30] and 100 patients with acute myeloid leukaemia (AML), of whom 86 came from three published cohorts (42, 26 and 18 patients) [36, 35, 48] and 14 were profiled in-house. The ten control marrows came from one of the AML studies [36]. Each patient was assigned to a major cytogenetic group from the metadata of its study.

#### Per-patient clustering

Each donor was clustered on its own. We selected 2,000 highly variable genes, computed a PCA and ran Leiden clustering at resolution 1.0, which deliberately over-clusters a donor into 10–20 raw clusters. Clusters were then merged unless they differed by genes that were on in one and off in the other. A gene counted as on in a cluster when it was detected in at least 35% of its cells and as off when detected in at most 10%, and two clusters stayed separate if either had at least five genes that were on in it and off in the other. Pairs that failed this rule in both directions were merged, the most similar pair first (correlation of PCA centroids), and the test was repeated until every remaining pair was distinct. An off call was ignored when the cluster showing it detected fewer than 60% as many genes per cell as the other cluster. Cell-cycle, histone, stress and immediate-early, interferon-response, mitochondrial and ribosomal genes, long nuclear transcripts and immunoglobulin V, D and J segments could not separate clusters, so cycling, stressed and low-quality versions of a population collapsed into it. This gave 1,370 clusters. Sequencing depth was low (a median of about 400 genes detected per cell), which makes small clusters the least reliable.

#### Annotation of blast and healthy clusters

Every cluster received a lineage from the detection rates of about 150 marker genes and was then labelled healthy or blast. In B-ALL, T, NK, mature B, plasma, erythroid, monocytic and dendritic cells, normal progenitors and normal pre-B cells were healthy, and every early-B cluster with its own on/off genes was a blast cluster. In AML, lymphoid, erythroid and stromal clusters were healthy, as were monocyte-like and dendritic-like populations that made up less than 5% of the patient; any myeloid population at 5% or more, progenitor-like, promyelocyte-like and mast- or basophil-like populations of any size, and clusters expressing HOX genes were blast clusters. The ten control marrows each contained four to eight normal myeloid and progenitor clusters, and the abundance thresholds were set above this range. Fifty-five clusters were merged into their parent blast cluster as false splits (residual cycling clusters, shallow copies lacking only abundant cytoplasmic genes, and blasts carrying haemoglobin or myeloid transcripts from doublets or ambient RNA). A small cluster with a shared signature (IRF4, DTX1, RUBCNL, ACSM3 and light chains) recurred in six B-ALL patients of different subtypes and was labelled as normal pre-B cells. The on/off gene list of every secondary B-ALL blast cluster was read individually, whereas the AML labels were rule-based. In total 882 clusters were healthy (231,887 cells) and 488 were blast (1,064,118 cells). The 5% rule and the treatment of small myeloid populations are decisions of the annotator; expression alone cannot show that a myeloid cluster is leukaemic.

#### Heterogeneity index

Blast heterogeneity was summarised per patient as the effective number of blast clusters, the exponential of the Shannon entropy [25, 31]. With *p*_*i*_ the fraction of the patient s blast cells in blast cluster *i* of *k*,

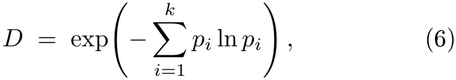

which is 1 for a single population and *k* for *k* clusters of equal size; two clusters holding 87% and 13% of the blasts give 1.47. Healthy cells do not enter the index.

#### Scoring with merged, split and shuffled labels

We sampled 300 blast cells at random from every patient with at least 300 blasts (183 patients, 99 with AML and 84 with B-ALL), which keeps each blast cluster at its true proportion, and removed blast clusters with fewer than 20 sampled cells (53,927 cells remained). scINTILLA was run three times on these cells. In the *merged* run the label was the patient (183 labels). In the *split* run the label was the patient together with the blast cluster (329 labels), so that each label covered one annotated population. In the *shuffled* run the blast-cluster labels of the split run were permuted among the cells of each patient, which keeps the number and the sizes of the labels but removes their biology. All runs used the sample-aware latent space (*z*) of an MrVI model [3] fitted to the atlas as the embedding, the fast preset of scINTILLA (the K-means family and Leiden for clustering; logistic regression, random forest and *k*-nearest neighbours as classifiers) and one random seed; the shuffled run used one permutation. The heterogeneity index was recomputed from the sampled cells, so that the reference describes the cells that were scored.

For each patient, the merged value of a quantity is that of the patient label, and the split value is the mean over the patient s blast-cluster labels, weighted by their cell numbers. Every quantity was oriented so that higher is better, and its *change on splitting* is the split value minus the merged value. For label fragmentation (the mean over the selected clusterings, Section 2.4) this change is the reduction in fragmentation. Because merged and split values are computed on the same cells of the same patient, properties of the patient that affect both, such as its similarity to other patients or its blast fraction, cancel in the difference. If the scores detect hidden populations, then within each disease the patient label should score worse as the index rises, the change on splitting should rise with the index, and patients with one blast cluster, whose two labels are identical, should show little change. Composite scores are min-max scaled over the labels of a run, and the merged and split runs hold different label sets, so, as in the re-scoring of the atlases, we rely on the unscaled quantities and report the composites for completeness only.

#### Statistics

The two diseases were analysed separately throughout, because the AML index depends on the 5% rule. Correlations are Spearman rank correlations. Patients with several blast clusters were compared with patients with one by a one-sided Mann-Whitney U test, reported with its AUC, and annotated with shuffled splits by a one-sided paired Wilcoxon signed-rank test. Patients with one blast cluster carry the same label in the annotated and shuffled runs, so comparisons with the shuffled run are restricted to patients with several clusters. The change on splitting was not tested against zero, because splitting the labels of other patients changes the label set and shifts the scores of single-cluster patients slightly (between −0.12 and 0.07 for fragmentation). The analysis was repeated inside every cytogenetic group with at least six patients, using the scores of the run on all patients, and across groups after subtracting each group s median from both the index and the change.

## 3 Results

### 3.1 sclNTlLLA scores each label from three complementary signals

scINTILLA asks how well a cell-type label is supported by the structure of the data, and it does so from three directions (Fig. 1a). The first signal, clustering confusion, comes from the unsupervised arm. scINTILLA runs a panel of clustering algorithms - the K-means family, hierarchical agglomerative clustering with several distances and linkages, DBSCAN, HDBSCAN [44], Louvain [2] and Leiden [65] - over a range of parameters, keeps the six partitions that best match the annotation, and measures for every cell how often its nearest neighbours share its cluster but carry a different label. The second signal, label fragmentation, uses the same six clusterings but looks from the label s side: a label whose cells are spread over several clusters is fragmented. The third signal, supervised consistency, comes from a classifier ensemble trained to reproduce the annotation, and summarises how much the classifiers agree, how confident they are and how uncertain their predicted distribution is.

The three signals are scaled within the dataset and given equal weight, which yields the per-label score scINTILLA-F (Section 2.4). The output is a ranked review list together with a diagnostic profile that shows which of the signals failed for each flagged label. The profile matters because the signals respond to different problems. Two populations sharing one label tend to raise fragmentation without confusing the classifiers, whereas one population divided into two labels confuses the classifiers and raises clustering confusion without fragmenting either label (Fig. 1b).

### 3.2 A benchmark with known annotation errors

In a single-cell atlas the labels are not a fixed ground truth. They group cells that were judged to be similar, and the resolution of those groups is partly a choice. A low score in a real atlas therefore has no reference against which it can be checked. For this reason we introduced errors of known type and location into real annotations and asked whether the damaged labels were ranked first (Fig. 1b,c). In five atlases – the adult human brain [59], the human lung [58], the human retina [38], the Human Endoderm Organoid Cell Atlas (HEOCA) [72] and the Human Neural Organoid Cell Atlas (HNOCA) [24] - we merged pairs of labels, contaminated labels with 30% foreign cells, or split labels at random into two halves. Each error type damaged three labels per annotation and was repeated ten times per atlas, giving 150 perturbed annotations. As in the real-atlas analyses, every label was then capped at 500 cells, so that label size gave no cue. The analysis, predictions and decision rules were preregistered before any score was computed. scINTILLA was compared with four existing measures of label or cluster quality: the Silhouette width [54] (hereafter the Silhouette baseline), kNN purity, SCCAF [45] and ROGUE [39] (Section 2.5).

### 3.3 Label fragmentation is needed to detect merged labels

To see what each signal contributes, we compared scINTILLA-F with the same score computed without fragmentation, that is, from clustering confusion and supervised consistency alone (scINTILLA-CS; Fig. 1d, Fig. 2a). scINTILLA-CS detected over-split labels almost perfectly (AUROC 0.991, a near-tie with SCCAF at 0.993) but merged labels poorly (0.678), giving a balanced AUROC of 0.851. To find out why, we scored the damaged labels with each signal on its own (Supplementary Fig. 2a). Classifier agreement detected over-splits well (AUROC 0.99) but merges poorly (0.68). When two populations share one label, the classifiers learn that combined label and predict it consistently, so nothing looks wrong to them. Fragmentation behaves in the opposite way. On its own it detected merges (0.87) and contamination (0.87) well, but it was close to chance for oversplits (0.63), because two random halves of one population spread over the clusters in the same way as the undivided label did.

**Fig. 2:**
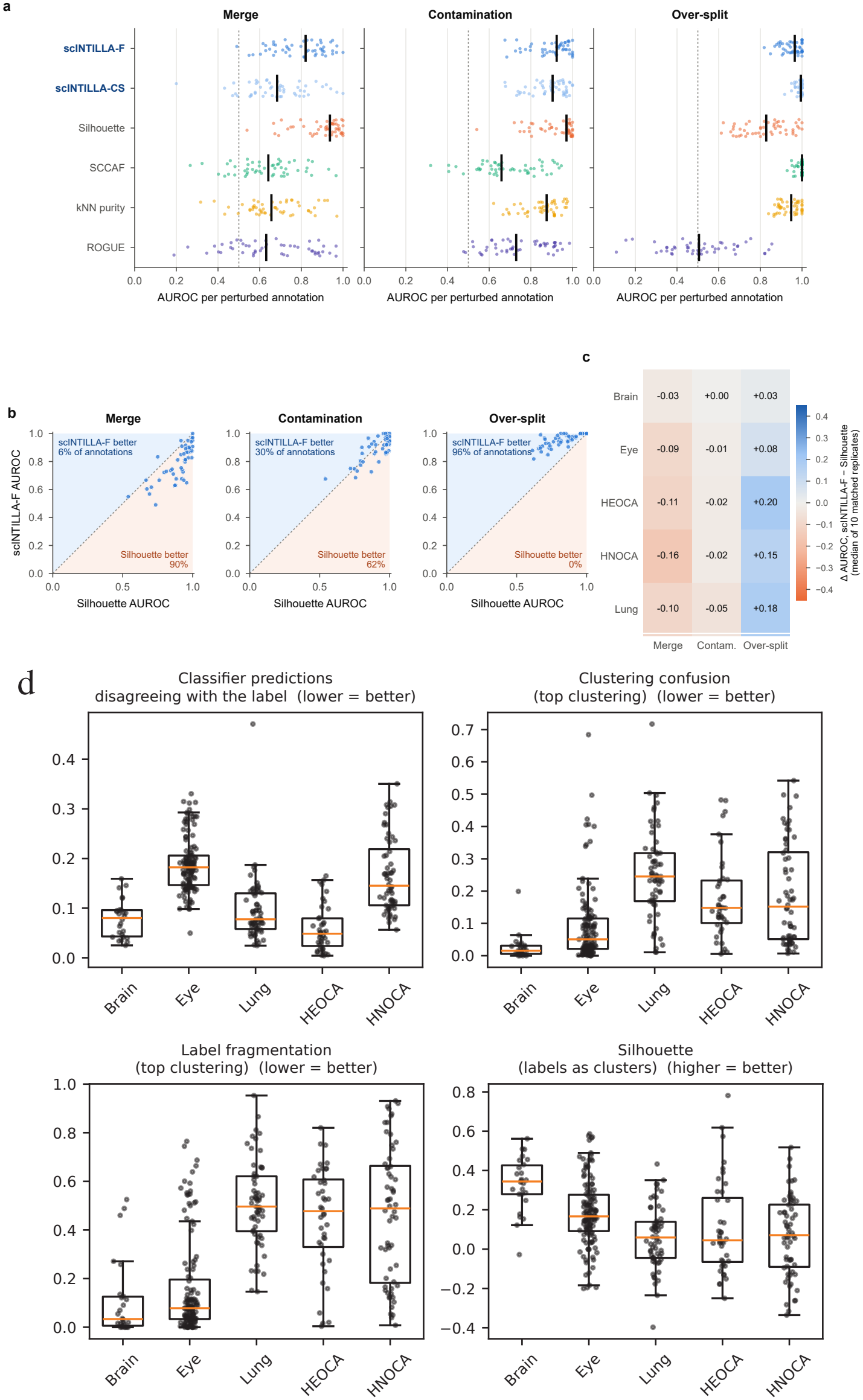
Benchmark results and the coherence of five real atlases. **(a)** AUROC for each of the 150 perturbed annotations, by error type, for scINTILLA-F, scINTILLA-CS (without fragmentation) and four baselines; vertical black bars mark medians and the dotted line marks chance. **(b)** scINTILLA-F against Silhouette baseline AUROC, one point per perturbed annotation; the percentages give the share of annotations in which each method was better. **(c)** Difference in AUROC between scINTILLA-F and the Silhouette baseline for each atlas and error type (median of ten matched replicates); blue favours scINTILLA-F and red favours the Silhouette baseline. **(d)** Unscaled per-label quality measures in the five real atlases: the fraction of classifier predictions that disagree with the label, clustering confusion and label fragmentation for the top clustering (lower is better), and the Silhouette width of each label (higher is better). Each point is one label; boxes show the median and interquartile range.

Including fragmentation therefore made the score more balanced. On matched labels, scINTILLA-F outperformed scINTILLA-CS by +0.110 AUROC for merges (95% interval +0.055 to +0.175), with the gain positive in all five atlases, and by +0.011 for contamination (+0.004 to +0.019). The cost was a small loss for over-splits (−0.033, −0.055 to −0.013). Overall, the balanced AUROC rose from 0.851 to 0.890 (paired difference +0.029, +0.005 to +0.059; Supplementary Fig. 2b,c). On its own, scINTILLA-F matched the Silhouette baseline (0.896; paired difference +0.003, −0.007 to +0.014) and outperformed SCCAF (0.770; +0.111), kNN purity (0.815; +0.067) and ROGUE in every atlas. ROGUE reached a balanced AUROC of only 0.651 (+0.238 in favour of scINTILLA-F), even though it was evaluated only on the 99% of labels it could score (Fig. 1d; Supplementary Fig. 14).

### 3.4 sclNTlLLA and the Silhouette baseline are complementary

The Silhouette baseline proved strong. With a balanced AUROC of 0.896, it was level with scINTILLA-F on its own, and it was clearly the best method for merges (0.933). The comparison is uneven across error types, however. Annotation by annotation, scINTILLA-F outperformed the Silhouette baseline in only 6% of the merge annotations, but in 30% of the contamination annotations and in 96% of the over-split annotations, where the Silhouette baseline was better in none (Fig. 2b). The same pattern held in every atlas. The Silhouette baseline led on merges in all five atlases (by 0.03–0.16 AUROC), whereas scINTILLA-F led on over-splits in all five, by up to 0.20 (HEOCA 0.20, lung 0.18, HNOCA 0.15; Fig. 2c). This is what one would expect from the error types. A merge places two distinct populations under one label and directly lowers its Silhouette width. Two random halves of one population, by contrast, can each have a reasonable Silhouette width while being impossible to tell apart, which is what classifier disagreement detects.

Because the two scores capture different information, we combined them with a parameter-free rank fusion, the mean of the two percentile ranks. The fusion of scINTILLA-F and Silhouette had the highest balanced AUROC of the eleven methods that score every label (0.908; merge 0.886, contamination 0.932, over-split 0.907), and the four best of these eleven methods were all scINTILLA-based (Fig. 1d). Two learned fusions, trained on separate development perturbations, did no better (0.904 and 0.903). This was not simply a benefit of fusing anything with Silhouette. Among all the methods fused with Silhouette in the same way, only the scINTILLA scores raised the balanced AUROC of the Silhou-

ette baseline, and scINTILLA-F was the only one to do so in every atlas (+0.014, 95% interval +0.006 to +0.024; scINTILLA-CS +0.011, −0.005 to +0.025, four of five atlases). Fusing Silhouette with SCCAF (−0.019), kNN purity (−0.007) or ROGUE (− 0.063) diluted it instead (Supplementary Fig. 3a), and the scINTILLA-F fusion outperformed each of these fusions in every atlas (+0.035, +0.022 and +0.077; Supplementary Fig. 7).

The gain over the Silhouette baseline is small, however, and should be read with care. With five atlases no contrast survives correction for multiple testing, so the interval is an exploratory effect size. Which score to use depends on how many labels a reader will inspect. For a single label, the Silhouette baseline is the better guide: precision@1 was 0.60 against 0.42 for the fusion, average precision 0.541 against 0.478 and recall@3 0.339 against 0.328 (Supplementary Fig. 12). A severe merge gives a label an extreme negative Silhouette width, and that label goes straight to rank one. For a review list the order reverses. Among the lowest tenth of labels the fusion found more of the damage (precision 0.40 against 0.35; recall 0.55 against 0.52), because it brings over-split labels into the list, which Silhouette cannot see. An annotator rarely stops at one label, and the fusion also says which signal failed. Its gain in AUROC comes from over-splits (+0.080, +0.043 to +0.114; better in 48 of the 50 oversplit annotations and worse in none), and the fusion is worse on merges (−0.037; Supplementary Fig. 3c,d; peratlas values in Supplementary Fig. 5). Across 66 mixtures of the three error types, the fusion outperformed the Silhouette baseline in 64%, losing only when merges dominated (Supplementary Fig. 6). Our reading is that scINTILLA complements the Silhouette baseline and does not replace it: it adds information that the Silhouette baseline lacks, chiefly about labels that divide a population without a transcriptional basis, and it reports which signal is responsible.

The ranking held when the benchmark design was changed. When 20,000 cells were sampled uniformly from each atlas instead of capping every label, scINTILLAF + Silhouette again ranked first among the methods that score every label (balanced AUROC 0.874; Silhouette baseline 0.863; Supplementary Fig. 8). With a batch-corrected (Harmony) embedding, scINTILLA scores occupied the top five places, and scINTILLA-F + Silhouette again improved on the Silhouette baseline (+0.024, +0.004 to +0.055, four of five atlases; Supplementary Tables 1 and 2). Harmony lowered the balanced AUROC of every method that uses the embedding, the baselines more than scINTILLA (Silhouette 0.896 to 0.857; kNN purity 0.815 to 0.747; scINTILLA-F 0.890 to 0.872). The order at the top changed: scINTILLA-CS + Silhouette came first (0.886), just ahead of scINTILLA-F + Silhouette (0.882). We suspect that removing donor structure leaves merged and contaminated labels less fragmented, so that fragmentation adds less. No single atlas drove the results: when each atlas was left out in turn, the gain of scINTILLA-F + Silhouette over the Silhouette baseline stayed between +0.010 and +0.016, and every balanced contrast of scINTILLA-F against a baseline kept its sign (Supplementary Fig. 13). On the real atlases, the labels flagged in the case studies also stayed in the review list when the scores were recomputed on the Harmony-corrected or scVI embeddings (Supplementary Table 3).

### 3.5 Applying sclNTlLLA to five Human Cell Atlas datasets

We next scored the full label sets of the five atlases, using the configuration validated in the benchmark (Section 2.5). Because scINTILLA-F is scaled within each atlas, we did not compare atlases on it. Comparing the means of a within-atlas score would mostly reflect how labels are spread inside each atlas, not how good the atlas is. Instead we compared the unscaled per-label quantities (Fig. 2d). The adult brain atlas was the most coherent on clustering confusion (median over labels 0.015), fragmentation (0.034) and Silhouette (0.344), and the eye atlas was second on all three (0.051, 0.078 and 0.167). The lung, HEOCA and HNOCA atlases were less coherent, with median fragmentation of 0.48–0.50 and median Silhouette below 0.08, and their order depended on the measure. Classifier disagreement gave a different picture. It was highest in the eye atlas (median 0.18), partly because the eye atlas has 123 labels and the ensemble includes weak classifiers. Coherence did not simply follow the number of labels: the eye atlas has 123 labels yet was the second most coherent on the unsupervised measures, whereas the lung and HNOCA atlases each have about 60. Plausibly, the differences reflect how distinct the populations of each tissue are and how much prior knowledge guided their annotation.

Within each atlas, the review list (scINTILLA-F fused with Silhouette) placed broad or catch-all labels at the top, as expected. These included “Unknown” in the lung atlas (rank 1 of 61) and “neuron” in the brain (rank 1 of 26) and HNOCA (rank 3 of 60) atlases. Because every label was scored on an equally sized sample, most baselines also placed these catch-all labels near the top, so on these labels the methods agree. Per-label scINTILLA-F scores, with their components and confusion networks, are given for each atlas in Supplementary Figs. 16–22. The labels that were not catch-alls were more informative, and the diagnostic profile showed that different labels were flagged for different reasons (Supplementary Fig. 4c). Monocytederived macrophages in the lung (rank 3 of 61) were low on every signal, most strongly on classifier consistency, whereas the HEOCA “stem cell” label was flagged most strongly by fragmentation, and mesodermal cells in HEOCA mainly by classifier consistency, with confusion and Silhouette also low. On the real labels, scINTILLA-F correlated only partly with the Silhouette baseline (mean Spearman *ρ* = 0.73) and weakly with ROGUE (0.13), so the scores rank real labels differently (Supplementary Fig. 4b). We followed up two flagged populations that are biologically well defined rather than vaguely named: the monocyte-macrophage compartment of the lung atlas and the “stem cell” label of HEOCA, which ranked first for review among the 40 HEOCA labels. scINTILLA-F + Silhouette was the only score that ranked this label first (Silhouette and kNN purity ranked it third, scINTILLA-F and SCCAF fourth).

As a by-product, scoring the atlases also benchmarks the classifiers and clustering methods themselves (Supplementary Fig. 1). Averaged over the five atlases, the stacking ensemble, logistic regression and multi-layer perceptron gave the highest F1 scores (0.910, 0.910 and 0.905). Na’ive Bayes, gradient boosting and LightGBM gave the lowest (0.52-0.61), and gradient boosting and LightGBM failed badly on the eye atlas (F1 0.12 and 0.17), as did LightGBM on HNOCA (0.14). Among the clustering methods, Leiden and Louvain achieved the highest ARI in most atlases. HDBSCAN was the best method on HEOCA (ARI 0.59) and the brain (adjusted mutual information [66], AMI 0.96), but it returned no usable partition for the lung and HNOCA atlases. Spherical K-means and hierarchical clustering stayed close to the graph-based methods. Scoring an atlas can therefore also help in choosing a classifier for label transfer, and the results suggest that classical clustering methods remain useful as independent checks.

### 3.6 Lung atlas: the monocyte-macrophage compartment

In the lung atlas, prediction agreement and clustering confusion were strongly anti-correlated across labels (Pearson *r* = −0.787; Fig. 3a). The “Unknown” label was an extreme outlier, and monocyte-derived macrophages and alveolar macrophages were among the labels with the highest confusion. The confusions of the monocyte-derived macrophages pointed in several directions. Of their misassigned predictions, 28% went to MTpositive alveolar macrophages, 22% to interstitial perivascular macrophages, 15% to alveolar macrophages, 12% to “Unknown” and 7–10% each to non-classical monocytes, classical monocytes and CCL3-positive alveolar macrophages. In the confusion network “Unknown” was linked to five of the monocyte and macrophage labels, and the heaviest of all its links was to the monocytederived macrophages (Fig. 3b). To test whether these confusions reflected genuine similarity, we re-clustered the monocyte-macrophage compartment together with the unannotated cells. A total of 297,952 cells previously labelled “Unknown” co-segregated with the monocytemacrophage cells (Fig. 3c). “Unknown” cells were the majority of every refined macrophage state: 86% of the cycling macrophages, 77% of the interstitial perivascular macrophages, 76% of the alveolar macrophages and 62% of the TREM2-high monocyte-derived macrophages described below. This was also the compartment in which the original study reported marked, disease-linked heterogeneity [58].

**Fig. 3:**
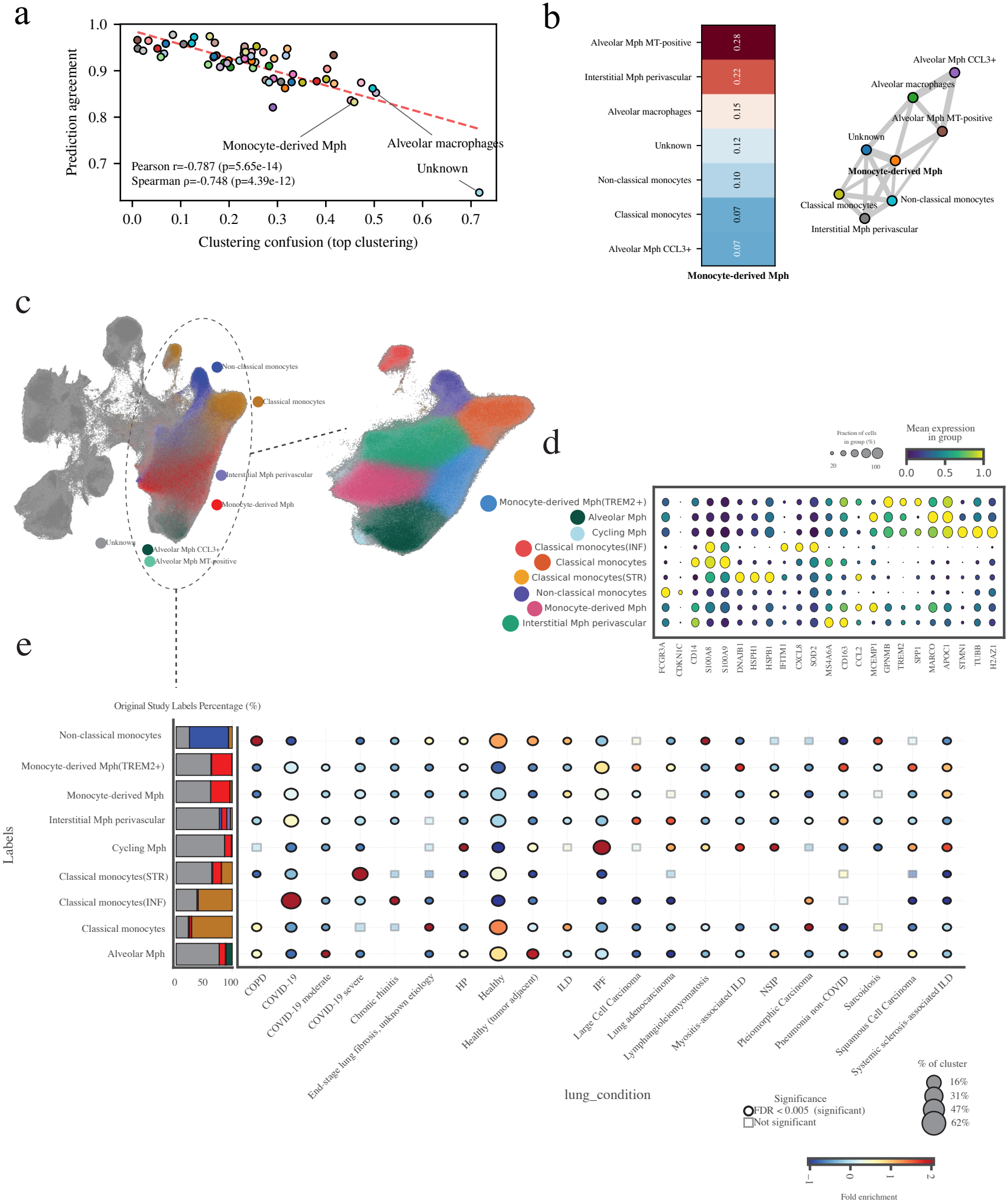
Re-analysis of the monocyte-macrophage compartment of the lung atlas. **(a)** Prediction agreement against clustering confusion (top clustering) for each label in the lung atlas, with the Pearson and Spearman correlations; “Unknown”, monocyte-derived macrophages and alveolar macrophages are labelled. **(b)** Left: the labels that monocytederived macrophages are confused with, coloured and annotated by the fraction of confused predictions. Right: confusion network of the monocyte, macrophage and “Unknown” labels; edge width is proportional to the amount of confusion. **(c)** UMAP of the lung atlas highlighting the monocyte-macrophage compartment (dashed outline; grey, other cells), with the re-clustered compartment enlarged and coloured by the new labels. **(d)** Marker gene expression across the re-annotated populations; dot size shows the fraction of expressing cells and colour the scaled mean expression. **(e)** Left: composition of each new population by original study label. Right: enrichment of each population across lung conditions; dot size shows the percentage of the cluster, colour the fold enrichment, and outlined circles mark significant enrichment (Fisher s exact test on cells, Benjamini–Hochberg FDR < 0.005; descriptive, with donor-level tests in Supplementary Table 5). The refined “Alveolar Mph” state is composed of cells originally labelled “Unknown”, “Alveolar Mph CCL3+”, “Monocyte-derived Mph” and “Alveolar Mph MT-positive”; the atlas label “Alveolar macrophages” is not among the retained cells.

The original study had also reported elevated labeltransfer uncertainty for macrophages in disease samples, with high-uncertainty macrophages in idiopathic pulmonary fibrosis (IPF) carrying a profibrotic programme [58]. Together with the low scINTILLA score and the large number of unannotated cells, this suggested that the existing labels did not resolve the structure of the compartment. We therefore re-annotated it from marker genes (Fig. 3d) and identified four new transcriptionally distinct states. The first was TREM2-high monocyte-derived macrophages, which expressed GPNMB, TREM2 and SPP1. The second was cycling macrophages (STMN1, TUBB, H2AZ1). The remaining two were subsets of classical monocytes distinguished by their dominant programme: an inflammatory subset (IFITM1, CXCL8, SOD2; classical monocytes, INF) and a stressed subset dominated by heat-shock and proteostasis genes (HSPH1, HSPB1, DNAJB1; classical monocytes, STR).

Mapping these states onto the clinical metadata showed disease-specific enrichment (Fig. 3e). The tests in that panel treat cells as units and are descriptive, so we repeated them with donors as units and give most weight to the enrichments that held (Supplementary Table 5; Supplementary Fig. 10). TREM2-high monocyte-derived macrophages were enriched in non-COVID pneumonia, at cell level and at donor level (12 donors; median share of a donor s cells 0.38 against 0.14 elsewhere; FDR 0.007). Their cell-level enrichments in myositis-associated interstitial lung disease (ILD) and in squamous cell carcinoma rest on one and three donors, which is too few to support a claim. That this state appears across lung conditions fits evidence that TREM2-expressing monocyte-derived macrophages are profibrotic. TREM2 is expressed mainly on monocyte-derived alveolar macrophages in fibrotic mouse lungs, is raised in lung macrophages from patients with IPF, and regulates macrophage survival and profibrotic activity [13]. The TREM2-high state also corresponds to the population that the original study singled out: in IPF, alveolar macrophages with high label-transfer uncertainty upregulated SPP1, PLA2G7 and CCL2 and downregulated PPARG and FABP4 [58]. TREM2-high macrophages had the highest SPP1 (Fig. 3d) and PLA2G7 expression of the nine states and lower PPARG and FABP4 than the alveolar macrophages. Among IPF cells that the atlas had transferred as alveolar macrophages, they made up 26% of the half with the higher uncertainty scores and 8.5% of the other half. They were also enriched in IPF, at cell level and with donors as units (52 donors; median share 0.33 against 0.12 elsewhere; Supplementary Table 5). The label that scINTILLA flagged for review thus led to the disease-affected subset that the original study had found with its uncertainty scores, recovered here without disease labels or uncertainty scores.

Cycling macrophages were enriched in IPF, and this was the best-supported enrichment in the compartment (52 donors; median share 0.035 against 0.003 elsewhere; FDR 3 ×10^−13^). At cell level they were also enriched in hypersensitivity pneumonitis, myositis-associated ILD and non-specific interstitial pneumonia, but these rest on three, one and three donors; the first was supported by one of the two donor-level tests and the other two could not be tested. The IPF result agrees with singlecell studies of fibrotic lung disease, in which proliferating macrophages are expanded in the fibrotic lower lobes in IPF and co-localise with myofibroblasts in a way that suggests a direct role in fibrotic niches [47]. Here these cells were largely recovered from previously unannotated cells, which adds independent evidence for macrophage proliferation in IPF. The stressed classical monocytes were enriched in COVID-19 (58 donors; FDR 2 ×10^−10^). The cell-level enrichment was strongest among severe cases, but one of those six donors supplied 89% of the cells, so we make the claim for COVID-19 as a whole. A heat-shock response has also been reported in severe SARS-CoV-2 disease: in a single-cell study of SARS-CoV-2-associated paediatric encephalopathy, HSPA1A and HSPB1 were upregulated across several immune cell types in the severe case, but not in mild encephalopathy or in febrile seizures caused by other pathogens [64]. That study was small and did not examine lung tissue, so the parallel is suggestive rather than direct.

We also asked whether the new labels were better supported by the data, by re-scoring the same cells with the original and the refined labels (Supplementary Table 4). The answer was mixed. With the refined labels, classifier disagreement fell from 0.157 to 0.107 and the mean Silhouette rose from− 0.042 to 0.000, but clustering confusion (0.295 to 0.347) and fragmentation (0.437 to 0.751) both rose. Going from seven to nine labels raises confusion and fragmentation mechanically. We had also suspected the embedding, because the refined labels were defined in the MrVI latent space and scored in an uncorrected PCA, but repeating the re-scoring on a Harmony-corrected PCA gave the same pattern: the Silhouette was again higher with the refined labels (−0.056 to −0.029), and clustering confusion still rose (0.396 to 0.572). The rise therefore follows from the finer labels and not from the uncorrected embedding. The support for the re-annotation therefore rests mainly on the biology: distinct marker programmes and coherent disease associations.

### 3.7 HEOCA: the “stem cell” label

The Human Endoderm-derived Organoid Cell Atlas (HEOCA) integrates nearly one million single-cell transcriptomes from 218 samples covering organoids of many endodermal origins, derived from pluripotent (PSC), fetal (FSC) and adult (ASC) stem cells under a range of protocols [72]. In this atlas too, prediction agreement and confusion were strongly anti-correlated (Pearson *r* = −0.909). The “stem cell” label, which ranked first for review among the 40 HEOCA labels, sat in the low-agreement, high-confusion tail, next to a group of mesenchymal and stromal labels (Fig. 4a). Those labels - mesodermal cells, mesothelial cells, pericytes, stromal cells, interstitial cells of Cajal and smooth muscle cells - formed a densely connected confusion module (Fig. 4b). We read this module as transcriptional proximity along the mesenchymal spectrum. The “stem cell” label was not part of it. Its misassigned predictions went to enterocytes (31%), M cells of the gut (27%), colon epithelial cells (27%) and ciliated cells (15%), all of them epithelial, and none to the mesenchymal labels (Fig. 4b; Supplementary Fig. 19). The classifiers were thus confusing the label with differentiated epithelium, which is what the re-analysis below found inside it.

**Fig. 4:**
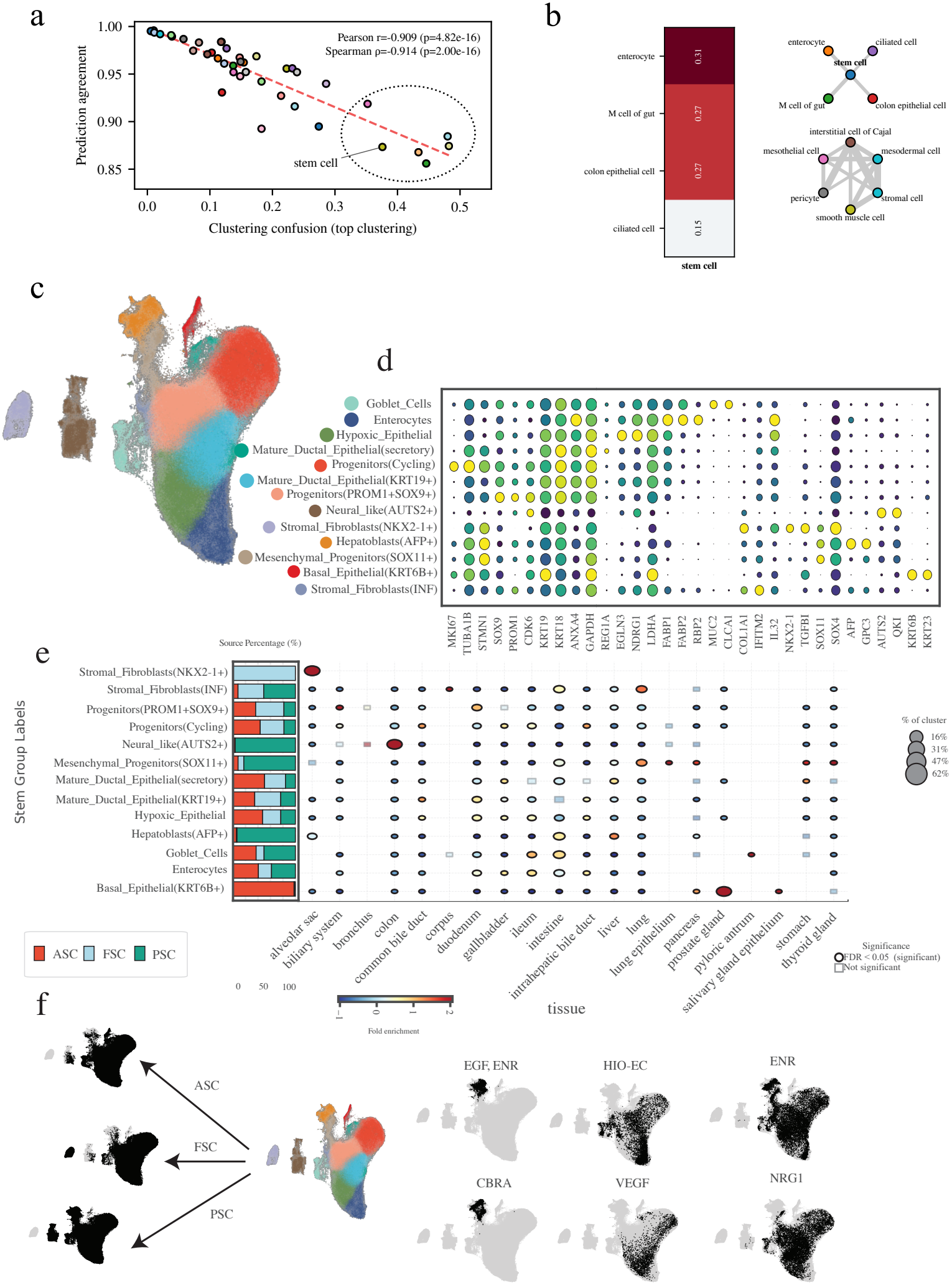
Resolution of the “stem cell” label in HEOCA. **(a)** Prediction agreement against clustering confusion (top clustering) for each HEOCA label, with the Pearson and Spearman correlations; the stem cell label and the circled mesenchymal and stromal labels lie in the low-agreement, high-confusion tail. **(b)** Left: the labels that the stem cell label is confused with, annotated by the fraction of misassigned predictions. Right: confusion network of the stem cell label, its partners and the mesenchymal and stromal labels circled in (a); edge width is proportional to the amount of confusion, and the two groups are not connected. **(c)** UMAP of the re-clustered stem cell group, coloured by the newly identified states. **(d)** Marker gene expression across the new states. **(e)** Left: proportion of cells of each state derived from adult (ASC), fetal (FSC) and pluripotent (PSC) stem cells. Right: enrichment of each state across tissues of origin; dot size shows the percentage of the cluster, colour the fold enrichment, and outlined circles mark significant enrichment (Fisher s exact test on cells, Benjamini–Hochberg FDR < 0.05; descriptive, with donor-level tests in Supplementary Table 5). **(f)** Left: cells of the stem cell group from each stem cell source (black) on the UMAP shown in the centre. Right: cells grown under each culture protocol (black), using the protocol categories of HEOCA [72]: “EGF, ENR”, HIO-EC, ENR, CBRA, VEGF and NRG1. Protocol information is available for 15% of the cells; the DS/SB protocol (six cells) is not shown.

To resolve the “stem cell” label, we took the stem-cell group (the label and the five epithelial labels it shared clusters with; Section 2.5), re-clustered it and identified a series of transcriptionally distinct states (Fig. 4c,d). They included cycling progenitors (MKI67, TUBA1B, STMN1) and SOX9/PROM1-positive progenitors. They also included KRT19-positive mature ductal epithelial cells, consistent with the ductal or cholangiocyte progenitor identity that marks biliary and hepatic organoidinitiating cells [29], and a secretory ductal subset. A neural-like population expressed AUTS2 and QKI. A hypoxic epithelial population was marked by EGLN3 and NDRG1, which encode well-established sensors and effectors of the HIF-mediated hypoxic response [16, 7]. Other states were AFP-positive hepatoblast-like cells and SOX11-positive mesenchymal-like progenitors. The SOX11-positive state is notable because SOX11 has a broad role in tissue remodelling during organogenesis, and its deletion causes defects that include hypoplasia of the lung, stomach and pancreas [61], the tissues in which we find SOX11-positive cells here. The remaining states were an inflammatory stromal fibroblast population, KRT6B-positive basal epithelial cells, a cluster formed entirely by one fetal alveolar organoid sample (NKX2-1-positive), and more differentiated enterocytes and goblet cells (Fig. 4d,e). Most of the enterocytes and goblet cells already carried those labels (64% and 76%); 13% and 8% had been labelled as stem cells. The other states came from the “stem cell” label itself: 97% of the neural-like cells, 82% of the hepatoblast-like cells and 80% of the SOX11-positive progenitors had carried it.

Mapping these states back to their tissue of origin gave a coherent picture (Fig. 4e), and each enrichment was then checked with donors as units (Supplementary Table 5). One state coincided exactly with a single fetal alveolar organoid sample at day 3 of culture (9,736 cells); its cells expressed NKX2-1 together with epithelial keratins and the stress-associated genes THBS1 and TGFBI (Fig. 4d). SOX9/PROM1-positive progenitors were enriched in biliary system organoids (five donors; median share of a donor s cells 0.41 against 0.13 elsewhere; donorlevel Mann-Whitney FDR 0.027), in agreement with the identification of PROM1-expressing cells in the peribiliary glands of the mouse extrahepatic bile duct as progenitors that give rise to cholangiocytes [73]. The neural-like population (AUTS2, QKI) came predominantly from PSC-derived colon organoids (found in five of the twelve colon donors), which may reflect off-target neural-like lineages arising in pluripotent differentiation protocols. Mesenchymal states were particularly prominent in PSC-derived lung epithelium and pancreas organoids. AFP-positive hepatoblast-like cells were concentrated in liver and intestinal organoids (19 donors each; donor-level Mann–Whitney FDR 0.026 and 0.003), consistent with AFP as a marker of hepatoblasts, the fetal liver progenitors that give rise to both hepatocytes and cholangiocytes [67]. KRT6B-positive basal epithelial cells came almost entirely (95%) from the single ASC-derived prostate gland organoid sample in the atlas, so their tissue association could not be tested across donors. SOX11-positive mesenchymal progenitors were found across lung, lung epithelium, pancreas, stomach and thyroid organoids, a distribution that mirrors the broad embryonic expression of SOX11 in these tissues. HEOCA donors include cell lines and study-level placeholders, so the donor-level tests are conservative, and a state found in one or two donors of a tissue could not be tested.

Stem cell source and culture protocol added further detail (Fig. 4f). Protocol information was available for 15% of the cells (56,033), using the protocol categories of the atlas [72]. Mesenchymal and hepatoblast-like states were concentrated in the CBRA (CHIR99021, BMP4 and retinoic acid) protocol and in the protocol annotated “EGF, ENR”, where together they made up more than 99% of the cells. The hypoxic progenitors were selectively enriched in VEGF-supplemented conditions, consistent with HIF-driven VEGF expression [19] and with the role of low oxygen in maintaining progenitor states [46]. They were almost absent from HIO-EC protocols (2.3% of cells), which suggests that endothelial co-culture relieves hypoxic stress rather than causing it. The canonical SOX9/PROM1 progenitor population was in turn almost absent from the hypoxic VEGF conditions (3.8% of cells).

Re-scoring the same cells with the six original labels and the thirteen refined labels again gave a mixed result (Supplementary Table 4). The mean Silhouette rose from 0.021 to 0.157 and fragmentation fell from 0.645 to 0.527, whereas clustering confusion rose (0.095 to 0.204) and classifier disagreement was unchanged (0.076 to 0.073). On the Harmony-corrected PCA the pattern was the same: Silhouette rose (−0.044 to 0.042), fragmentation fell (0.604 to 0.556) and confusion rose (0.270 to 0.382). As in the lung, the evidence for the new states lies mainly in their markers and in their coherent tissue and protocol origins. This is the situation scINTILLA was designed to detect: a single label covering cells of distinct identities, tissue origins and culture dependencies. scINTILLA flagged it without any prior knowledge of what the label contained, and the flag led to a data-driven re-annotation.

### 3.8 Leukaemia: patient labels that conceal several blast populations

In the benchmark we introduced the label errors ourselves, and in the two case studies we started from labels that scINTILLA had already flagged. We therefore looked for a setting in which merged labels arise naturally and can be annotated independently of the score. Acute leukaemia provides one. The blasts of a patient can form a single population or several, and in AML they are known to span progenitor-like and more differentiated myeloid states [21]. Labelling every blast cell only by its patient therefore reproduces the merge error naturally: the label covers one population in some patients and several in others. We annotated the blast clusters of 184 patients at diagnosis, 84 with B-ALL and 100 with AML, and summarised each patient by a heterogeneity index, the effective number of blast clusters (Section 2.6). Four patients illustrate its range (Fig. 5a). In a KMT2A-rearranged AML and a hyperdiploid B-ALL all blasts formed one cluster (index 1.00). In a BCR::ABL1 B-ALL the blasts formed four early-B clusters (2.73), and in a second KMT2A-rearranged AML they formed eight progenitor-like and monocyte-like clusters (5.33). Each of these clusters had its own set of genes that were on in it and off in the other clusters of the patient (Supplementary Fig. 9).

**Fig. 5:**
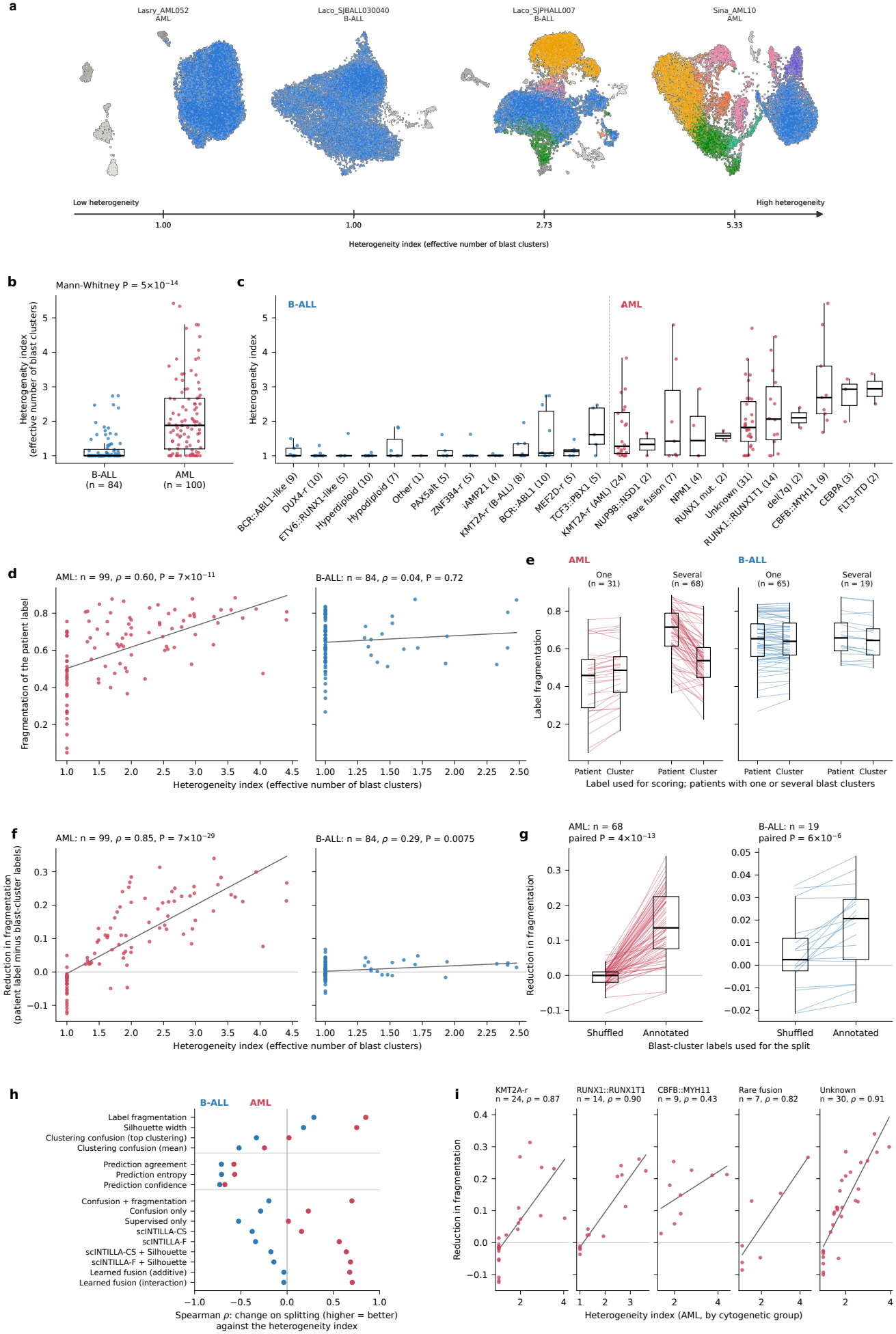
Blast heterogeneity in acute leukaemia and its detection by scINTILLA. **(a)** UMAPs of all cells of four example patients, ordered by heterogeneity index (two with a single blast cluster and two with several); cells are coloured by annotated blast cluster and healthy cells are grey. Marker genes of these clusters are shown in Supplementary Fig. 9. In the remaining panels AML is shown in red and B-ALL in blue. **(b)** Heterogeneity index (effective number of annotated blast clusters) of each patient at diagnosis; one-sided Mann-Whitney test. **(c)** Heterogeneity index by cytogenetic group, with the number of patients in brackets. **(d)** Fragmentation of the patient label (mean over the selected clusterings; lower is better) against the heterogeneity index recomputed on the 300 sampled blast cells of each patient, with the Spearman correlation. **(e)** Fragmentation of each patient scored with the patient label and with its blast-cluster labels (cell-weighted mean), for patients with one and with several blast clusters; one line per patient. **(f)** Reduction in fragmentation on splitting (patient label minus blast-cluster labels) against the heterogeneity index. **(g)** Reduction in fragmentation with shuffled and with annotated blast-cluster labels, in patients with several blast clusters; one-sided paired Wilcoxon test. **(h)** Spearman correlation between the change on splitting and the heterogeneity index for every scINTILLA quantity, the Silhouette width and the composite scores, each oriented so that higher is better. Composite scores are rescaled within each run and are shown for completeness. **(i)** As (f), inside the AML cytogenetic groups with at least six patients. Boxes show the median and interquartile range.

The two diseases differed (Fig. 5b,c). In B-ALL the median index was 1.00 and 52 of the 84 patients had a single blast cluster, whereas in AML the median was 1.88 and only 16 of 100 patients had a single cluster (Mann-Whitney AUC 0.81, *p* = 5 ×10^−14^). The index was low in every cytogenetic group of B-ALL and varied between the groups of AML; among those with at least six patients the median ranged from 1.28 in KMT2A-rearranged AML to 2.69 in CBFB::MYH11. The AML index depends on how we decided which myeloid clusters were leukaemic, so its values are not directly comparable with those of B-ALL, and we analysed the two diseases separately from here on. We then scored the same blast cells with two labellings, one that hides the annotated blast clusters (the patient) and one that exposes them (the patient together with the blast cluster), for 99 AML and 84 B-ALL patients. In AML the patient label was more fragmented the more blast clusters it concealed (Spearman *ρ* = 0.60, *p* = 7× 10^−11^; Fig. 5d). The median fragmentation was 0.72 for the 68 patients with several blast clusters and 0.46 for the 31 patients with one. The other signals pointed the same way: every score of the patient label fell as the index rose (*ρ* between −0.44 and −0.65; Supplementary Table 6). This is how the tool would be used in practice, because a user has only the patient label and no annotation of the blast clusters.

When each blast cluster received its own label, most of this extra fragmentation disappeared. In patients with several clusters the median fragmentation fell from 0.72 to 0.54, whereas in patients with one cluster it did not fall (Fig. 5e). The reduction in fragmentation grew with the index (*ρ* = 0.85, *p* = 7× 10^−29^; Fig. 5f). Its median was 0.136 in the patients with several clusters and −0.026 in those with one (AUC 0.98, *p* = 2 ×10^−14^). The Silhouette width showed the same pattern: it was lower for patient labels with a higher index (*ρ* = −0.65), and its improvement on splitting grew with the index (*ρ* = 0.75). The poor score of a heterogeneous patient therefore came from the blast clusters hidden under its label. Had it come from a general property of the patient, it would have remained after the clusters were labelled.

Splitting any label into smaller labels might reduce fragmentation by itself, even when the split is meaning-less. We therefore repeated the split with the blast-cluster labels shuffled among the cells of each patient, which keeps the number and sizes of the labels (Fig. 5g). In AML the shuffled split left fragmentation unchanged (median reduction 0.000), and the annotated split reduced it more than the shuffled one in all 68 patients with several clusters (paired Wilcoxon *p* = 4× 10^−13^). The shuffled reduction was unrelated to the index (*ρ* = 0.05), and the extra reduction that the annotated labels gave over the shuffled ones grew with the index (*ρ* = 0.67, *p* = 3 × 10^−10^). The relation also held inside cytogenetic groups (Fig. 5i): *ρ* = 0.87 in KMT2A-rearranged AML (24 patients), 0.90 in RUNX1::RUNX1T1 (14), 0.82 in the rare-fusion group (7) and 0.91 in patients of unknown cytogenetics (30). In CBFB::MYH11 all nine patients had several clusters, so there were no single-cluster patients to compare with, and the correlation was not significant (0.43, *p* = 0.24). To exclude an effect of cytogenetic group, we subtracted each groups median from the index and from the reduction; the reduction still rose with the index among the patients with several clusters (*ρ* = 0.76, 57 patients).

In B-ALL the effect was in the same direction but small. The fragmentation of the patient label did not follow the index (*ρ* = 0.04; Fig. 5d), so here the tool would not have flagged the heterogeneous patients from the patient label alone. Splitting reduced fragmentation only slightly (median 0.021 in the 19 patients with several clusters against −0.003 in the 65 with one; AUC 0.70, *p* = 0.005; *ρ* = 0.29 with the index). The reduction was nonetheless larger than with shuffled labels in 18 of the 19 patients (*p* = 6× 10^−6^), and the difference rose with the index (*ρ* = 0.56, *p* = 0.012), a result we regard as tentative at this sample size. The Silhouette width did not separate annotated from shuffled splits in B-ALL (*p* = 0.70), so fragmentation was the only signal that responded. No cytogenetic group of B-ALL had more than four patients with several clusters, and none could be tested on its own. The effect is probably weaker in B-ALL because its blast clusters are much more alike than those of AML. The AML blast clusters were distinct cell states, such as progenitor-like, monocyte-like and mast-cell-like populations, whereas the B-ALL blast clusters differed by 5-20 on/off genes and were usually small.

Each signal responded here in the same way as it did in the benchmark, where we had injected the errors our-selves (Fig. 5h; Supplementary Table 6). Fragmentation and Silhouette, the two measures that detected merges there, are the ones that responded to hidden blast populations here. The supervised signals moved in the opposite direction. Prediction agreement, entropy and confidence all worsened when the blast clusters were labelled, and worsened more in more heterogeneous patients (*ρ* between −0.57 and −0.73), because two blast clusters of one patient are harder for a classifier to tell apart than two patients. The shuffled split corresponds to the over-split error of the benchmark, and the supervised signals penalised it far more strongly: in AML the median prediction agreement fell by 0.148 with shuffled labels against 0.018 with the annotated clusters (*p* = 5 ×10^−13^), and in B-ALL by 0.054 against 0.037 (*p* = 0.004). The classifiers thus distinguished arbitrary from annotated subdivisions, and the unsupervised signals distinguished merged from unmerged labels. The composite scores cannot be compared between the merged and the split run, for two reasons: they combine signals that move in opposite directions, and they are rescaled over a different set of labels in each run; scINTILLA-F nevertheless followed the index in AML (*ρ* = 0.56), but not in B-ALL (− 0.34). For the question asked here, whether one label hides several populations, the unscaled fragmentation and the Silhouette width are the readouts to use.

## 4 Discussion

The need for scINTILLA has grown with the size of singlecell atlases. scINTILLA asks two questions of every label: how diverse are the cells it covers, and how well can the label be learned from the data? A label whose cells form one coherent population, and which classifiers recover consistently, is well supported. A label that covers several populations, or that classifiers cannot learn, is not. Together, diversity and learnability measure how well a label is supported by the data. The label can also be a sample: in our leukaemia atlas the patient label covered a single blast population in some patients and several in others, most often in AML. Much effort has gone into transferring, harmonising and propagating cell-type labels, and much less into checking whether the cells under each label form one consistent population before the label is reused. Classification and label-transfer tools treat the annotation as ground truth, so any heterogeneity or ambiguity hidden in a label passes silently into every analysis built on it. scINTILLA reverses this assumption: the annotation itself is what gets examined.

We tested the score against known errors, and we regard this as the core of the work. Without such a test, a label-quality score can only be judged by whether its lowest-ranked labels look plausible, an argument that is easy to make after the fact. The benchmark gave a less tidy, and more useful, answer than we expected. Without fragmentation (scINTILLA-CS), the score was excellent at detecting arbitrary over-splitting but weak at detecting merged labels. A score built mostly on classifier consistency can be fooled by exactly the errors that broad, catch-all labels represent, because a classifier readily learns a merged label. Adding label fragmentation addressed this weakness and improved merge detection in all five atlases. It also showed why the signals have to be combined. Each of them alone is blind to at least one type of error.

The Silhouette width, a simple and long-established measure, turned out to be a strong baseline, level with scINTILLA-F alone. We report this plainly because it follows from how the errors behave. A merge places two distinct populations under one label, and that is what the Silhouette width measures; an over-split, by contrast, can leave both halves well separated from other labels yet indistinguishable from each other, and only a measure such as classifier disagreement detects that. The two scores are therefore complementary, and their fusion gave the best detection of damaged labels among the methods that score every label. Unlike ROGUE, every scINTILLA variant scores all labels. The four best of those methods were all scINTILLA-based, and scINTILLA was the only signal tested whose addition improved the Silhouette baseline; fusing the baseline with SCCAF, kNN purity or ROGUE diluted it. The scINTILLA-F fusion improved on the Silhouette baseline in every atlas and ranked first under both sampling designs, and with a batch-corrected embedding scINTILLA scores took the top five places, with the fusion that omits fragmentation narrowly first. But the gain was small. The Silhouette baseline remained better on merges and at the very top of the list, and with five atlases the statistics are exploratory, so we present scINTILLA as a complement to the Silhouette baseline, not a replacement for it.

In practice we recommend the fused score for ordering the review list, scINTILLA-F as the per-label quality value, and the diagnostic profile for seeing why a label is flagged. When the question is whether a single label hides several populations, as with the patient labels of the leukaemia atlas, we recommend reading the unscaled fragmentation directly, together with the Silhouette width.

The biological re-analyses suggest that the flags point to real structure rather than to statistical artefacts. In the lung atlas, the flagged monocyte-macrophage compartment, together with nearly 300,000 cells labelled “Unknown”, resolved into transcriptionally distinct states with clear disease associations: TREM2-high monocytederived macrophages in non-COVID pneumonia, cycling macrophages in IPF and a heat-shock monocyte state in COVID-19, each supported with donors as units. In HEOCA, the “stem cell” label dissolved into progenitor, stromal and differentiated states whose tissue origins and protocol dependencies were biologically informative. Rescoring with the refined labels gave mixed results. Classifier separability and Silhouette improved, whereas clustering confusion rose. This is expected when the number of labels grows, and it was not an artefact of the uncorrected embedding, because the same pattern appeared after Harmony correction. We therefore rest the case for these re-annotations on markers and on disease or tissue enrichment, not on a single summary number.

Several of the recovered states are known. TREM2- and SPP1-expressing macrophages and proliferating macrophages have been described in the original lung atlas and in the fibrosis literature [58, 47, 13], and we take that as validation: scINTILLA was given no markers and no prior knowledge, yet it pointed to the compartment in which independent studies had found heterogeneity. What those studies do not change is the state of the annotation. Between 61% and 86% of the cells in each refined macrophage state carried the label “Unknown”, so the states were present in the atlas but could not be retrieved through its labels, and anyone transferring labels from it would have inherited the gap. Finding such places is what the tool is for. None of the baseline scores says why a label is flagged, and the diagnostic profile is what turned a low score into a specific re-analysis: confusion with the unannotated cells in the lung, and fragmentation of the “stem cell” label in HEOCA.

The leukaemia analysis tests the same idea from the opposite direction. In the case studies the score came first and the biology followed; here the heterogeneity was annotated first and the score was then compared with it, patient by patient. In AML, the fragmentation and the Silhouette width of a patient label followed the number of blast populations it concealed, the scores recovered when those populations were labelled, the recovery scaled with the amount of heterogeneity, held inside cytogenetic groups and was absent when the labels were shuffled. This is the behaviour that the injected merges predicted, now in a setting where the merge was not of our making, and it adds a graded result to the benchmark: the score did not only separate damaged from intact labels but tracked how much was hidden. The signals also kept their roles. The unsupervised signals responded to hidden populations, whereas the supervised signals penalised subdivision, mildly for annotated clusters and strongly for shuffled ones. The B-ALL result sets the limit. Blast subclusters that differed by a handful of genes did not lower the score of the patient label, and were visible only as a small recovery against the shuffled control. scINTILLA therefore detects a merged label when the populations merged under it are distinct cell states, as in AML. We do not claim that it detects populations that differ by only a few genes, as in B-ALL.

scINTILLA is advisory rather than prescriptive. It identifies labels that deserve a second look and shows the evidence, but it does not impose an alternative nomenclature or correct the annotation automatically. Whether a low score reflects an error, a deliberately coarse umbrella term or a genuine biological continuum remains a decision for domain experts. Because the framework is not tied to a fixed set of algorithms, new clustering and classification methods can be added to it.

Most of the limitations concern the benchmark. Its errors are synthetic, injected into five atlases that were also used during development. Fresh perturbations guard against overfitting to particular replicates, but they are not independent biological replication, and with five atlases the bootstrap intervals are wide. Errors in real annotations are also more varied than our three types, and the original labels used as intact controls are themselves imperfect. The leukaemia analysis complements the benchmark with merges that were not injected, but its reference is our own annotation. That annotation records on/off differences between clusters and was built from the same expression data that scINTILLA scores, so agreement shows that the tool recovers annotated cluster structure, not independently confirmed subclones. Merged labels remain better detected by the Silhouette baseline, and the supervised arm has two weaknesses of its own. Classifier consistency is computed on all cells, including the 80% used for fitting, so flexible learners can fit a noisy boundary and understate the inconsistency. The arm still separated over-split labels almost perfectly (AUROC 0.99), so the signal survives, but out-of-fold predictions would be the cleaner design; nor are weak classifiers filtered, and three of the twelve had a mean F1 between 0.52 and 0.61. The second weakness is that agreement, entropy and confidence are computed against the ensemble s modal prediction and do not refer to the assigned label. A foreign cell that every classifier assigns, confidently, to its true class therefore scores well under its wrong label. In the benchmark, where the foreign cells came from many labels, the arm still reached an AUROC of 0.86 for contamination; we expect the blind spot to matter most when the contaminating cells form one coherent population, and there fragmentation and confusion carry the detection. For the real atlases we therefore report the label-targeted disagreement alongside the composite.

The scores also depend on upstream processing. If integration is poor, they may reflect batch structure rather than biological heterogeneity. The embedding used here is a label-free PCA without batch correction; with a Harmony-corrected embedding, scINTILLA scores again occupied the top five benchmark places, and the casestudy labels remained in the review list (rank correlation with the PCA-based review scores 0.67–0.96; 0.83 for the scVI space of the eye atlas; Supplementary Table 3). Because scINTILLA-F is scaled within each atlas, it is best read as a ranking of labels inside one dataset, and comparisons between atlases need the unscaled quantities. Benchmarking many classifiers and clusterings also has a computational cost, which we mitigate by sampling cells per label (scoring took 8-95 minutes per atlas on 16 CPUs).

One limitation is built into the design. The clusterings used for confusion and fragmentation are chosen by their agreement with the annotation, and the classifiers are evaluated on cells drawn from the same annotation. The scores therefore measure how well the data support the labels, not whether a label is biologically correct. Choosing clusterings by their agreement with the annotation ties the unsupervised arm to the granularity the annotators chose. That is deliberate. The question is how well the data support the labels at that resolution, and the error this choice could hide, a merge, is the one that fragmentation detects (AUROC 0.87). For an annotation suspected of being too fine or too coarse throughout, the stability criterion of Section 2.2 is the better selector.

## 5 Conclusions

scINTILLA ranks the cell-type labels of a single-cell dataset by how well the data support them, combining clustering confusion, label fragmentation and supervised consistency into one score with a per-signal diagnostic profile. In a pre-registered benchmark of 150 annotations with injected errors, adding fragmentation made the score more balanced across error types, and fusing scINTILLA-F with the Silhouette width gave the best detection of damaged labels, improving on the Silhouette baseline in every atlas. The two capture different kinds of error and are best used together.

In the five Human Cell Atlas datasets, scINTILLA flagged populations whose labels concealed real hetero-geneity. Re-annotating the lung monocyte-macrophage compartment recovered disease-associated macrophage and monocyte states, largely from cells that had been left unannotated; in HEOCA, the broad “stem cell” label resolved into progenitor, stromal and differentiated identities with coherent tissue and protocol origins.

In acute leukaemia, where a patient label naturally covers one or several blast populations, the fragmentation of the label followed the annotated heterogeneity of AML patients and recovered when the populations were labelled separately.

The framework can in principle be applied to any annotated single-cell dataset, although it has so far been tested only on these five atlases and the leukaemia atlas. It finds where an annotation is weak and leaves interpretation and re-annotation to the biologist.

## Author Contributions

S.C. conceived the study design, designed the architecture of the pipeline, contributed to the software development, co-wrote the initial draft, and supervised the overall project. S.K. collected and curated the data, contributed to the pipeline development, performed the formal analyses, generated the visualisations, and wrote the initial draft. A.G. provided supervision and reviewed the manuscript. N.B., E.N., S.R., I.R. and R.Z. contributed equally to this work, jointly designed the methodology, developed the software framework, performed the formal analyses, and wrote the original draft.

## Acknowledgements

This work was in part carried out using core funding from Human Technopole, funded by the Italian Ministry of Economy and Finance (MEF). S.K. is a member of the European School of Molecular Medicine (SEMM). We thank Prof. Piercesare Secchi and Prof. Francesca Ieva, together with everyone involved in the Applied Statistics course at the Politecnico di Milano, for organising the collaboration that made this work possible. We are also grateful to Dr Oliver Harschnitz of the Human Technopole for his comments on a draft of the manuscript. S.C. thanks Dr Craig Glastonbury of the Human Technopole for his continuous support and encouragement.

## Code availability

The pipeline *scINTILLA*, created as part of this research, is publicly available on GitHub: https://github.com/soumickmj/scINTILLA

## Data availability

The five atlases analysed in this study are publicly available and were downloaded from existing sources. Data were accessed via the Human Cell Atlas data portal (https://data.humancellatlas.org/) and the underlying datasets are also deposited on the CellxGene data portal (https://cellxgene.cziscience.com/), maintained by the Chan Zuckerberg Initiative. All atlases are publicly available as part of the Human Cell Atlas initiative. The leukaemia atlas combines four published single-cell datasets [30, 36, 35, 48], available from the repositories named in the original publications, with 14 AML samples profiled in-house. The integrated leukaemia atlas and the in-house samples are part of ongoing studies and are not publicly available; they can be requested from the corresponding author.

## Ethics statement

With the exception of 14 in-house AML samples, all analyses were performed on previously published, deidentified single-cell datasets that are publicly available through the Human Cell Atlas and CellxGene data portals or the repositories of the original leukaemia studies. The in-house AML samples were obtained from the VIVO Biobank or Great Ormond Street Hospital; all patients, or their legal guardians, provided written informed consent in accordance with the Declaration of Helsinki for sample collection and research use. Ethical approval and, where applicable, informed consent were obtained by the original studies that generated these data, as described in their respective publications. No additional ethical approval was required for the analysis of the published datasets.

