## Supplementary Methods for "scINTILLA: Single-Cell Integrated Inference, Labelling, and Landscape Analysis for Assessing the Quality and Hidden Diversity of Cell Annotations"

Kanannejad *et al.*

The scINTILLA package is a general pipeline for single-cell analysis. This section describes the package in full. The clustering and classification modules, on which the label-quality score is built, are summarised in the main Methods and described in detail in Sections S5 and S6. The other sections describe modules that are part of the package but were not used for the analyses reported in this paper. The tables list the functions of the package with their inputs and outputs.

#### S1 Exploratory Data Analysis

This module of the scINTILLA package provides utilities for exploratory data analysis (EDA) of single-cell RNA sequencing (scRNA-seq) datasets. It extracts summary statistics in a standard form from AnnData objects, in which rows represent individual cells and columns represent genes.

There are four functions:

- **dataset_summary**: reports the total numbers of cells and genes, the sparsity of the expression matrix (the ratio of zero-value entries) and the average transcript count per cell.
- **expressed_genes**: identifies transcriptionally active genes, those detected in at least a user-defined minimum number of cells. This is a standard step in quality control (QC).
- **sample_counts**: counts the cells in each experimental condition or biological cluster stored in the metadata.
- **mean_expression_by_group**: computes the average expression of selected genes in each cell population, so that groups can be compared directly.

These summaries let researchers check the quality and composition of a dataset quickly, before any more complex downstream modelling.

#### S2 Preprocessing and Normalisation

Single-cell omics data are often influenced by technical variability, including differences in sequencing depth, library size and measurement noise. These effects can hide the underlying biological signal, so preprocessing and normalisation are needed before any further analysis. scINTILLA therefore provides a preprocessing framework that brings together transformation and normalisation strategies commonly used in single-cell RNA-seq and CITE-seq analyses [1, 22]. Instead of a single predefined pipeline, it implements 14 transformation methods, each designed for particular characteristics of the data such as sparsity, overdispersion or compositional effects. Formally, let $X \in \mathbb{R}^{n\times p}$ denote the raw count matrix, where $n$ is the number of cells and $p$ the number of features (genes or proteins). A preprocessing transformation can be defined as a function $T : \mathbb{R}^{n\times p} \to \mathbb{R}^{n\times p}$, which maps the raw data matrix $X$ to a transformed matrix $X' = T(X)$ in which technical variability is reduced and the data are better suited to statistical analysis. The transformations include standard normalisation approaches, such as library size scaling and log-transformation, and more advanced techniques based on variance stabilisation, distributional assumptions and probabilistic modelling. Gene selection and scaling can also be applied, to reduce dimensionality and to make features more comparable with one another. The framework also contains an automated benchmarking procedure, which evaluates several preprocessing strategies on a given dataset and picks the most suitable transformation according to predefined criteria. Because the choice is made on the data, it can differ between data modalities and experimental settings, and no preprocessing pipeline has to be selected by hand. All transformations operate directly on standard single-cell data structures and return a normalised expression matrix with the associated metadata preserved. Their main characteristics are described below, and their input parameters are summarised in Supplementary Table 7.

**Log shift with size factor normalisation (`log_shift_size_factor`)** [1] Counts are normalised by library size and then log-scaled. For each cell, the size factor is the total count across all genes, and each expression value is divided by it. The transformed value is:

$$x'_{ij} = \log\left(\frac{x_{ij}}{s_i} + 1\right) \tag{S1}$$

**Supplementary table 7:** Summary of transformation methods. All take as input a gene expression dataset (e.g. AnnData or DataFrame) and return a transformed dataset of the same structure.

| Method | Additional parameters |
|---|---|
| `log_shift_size_factor` | – |
| `arcsinh_transform` | $\alpha$ (default: 0.05) |
| `log_alpha_transform` | $\alpha$ (default: 0.05) |
| `log_cpm_transform` | – |
| `log_shift_scale_by_std` | – |
| `log_shift_size_factor_hvg` | `top_frac` (default: 0.35) |
| `log_shift_size_factor_z` | – |
| `log_shift_hvg_z` | `top_frac` (default: 0.35) |
| `normalise_scran` | – |
| `normalise_tmm` | – |
| `box_cox_transform` | – |
| `pearson_residuals_transform` | $\theta$ (default: 100) |
| `glm_pca_transform` | `n_components` (default: 50) |
| `sanity_transform` | – |

where $x_{ij}$ is the raw count of gene $j$ in cell $i$, and $s_i = \sum_j x_{ij}$ is the total count for cell $i$. Dividing by the size factor reduces variability due to differences in sequencing depth across cells; the logarithm then stabilises the variance and damps the effect of highly expressed genes. It is one of the most commonly used normalisation strategies in scRNA-seq analysis, and a reasonable default where no strong assumptions about the data distribution are warranted.

**Arcsinh transformation (`arcsinh_transform`)** [1] The arcsinh transformation applies a nonlinear scaling that behaves like a logarithm for large values but stays approximately linear near zero:

$$x'_{ij} = \operatorname{arcsinh}(\alpha \cdot x_{ij}) \tag{S2}$$

where $\alpha$ is a scaling parameter that controls the strength of the transformation, set to 0.05 by default. It is widely used for cytometry data (e.g. CyTOF and CITE-seq) because it compresses large values without distorting small ones too much. Its main use is protein expression data, or any dataset with a wide dynamic range in which differences among low-expression values must be preserved.

**Log-alpha transformation (`log_alpha_transform`)** [1] Here the standard log transformation is generalised by a scaling factor:

$$x'_{ij} = \log(\alpha \cdot x_{ij} + 1) \tag{S3}$$

where $\alpha$ controls the degree of compression applied to the data (default: 0.05). Tuning $\alpha$ adjusts how sensitive the transformation is to low versus high expression values. Smaller values of $\alpha$ give a milder transformation and larger values more compression. The method is therefore useful when the scale of the data varies across datasets, or when a standard log transformation is either too aggressive or not aggressive enough.

**Log CPM normalisation (`log_cpm_transform`)** [1] In this approach, counts are first converted to Counts Per Million (CPM) to account for differences in library size, and then log-transformed:

$$x'_{ij} = \log\left(\frac{x_{ij}}{s_i} \cdot 10^6 + 1\right) \tag{S4}$$

where $s_i$ is the total count for cell $i$. Scaling every cell to a common reference makes expression values comparable across cells, and the log transformation that follows stabilises variance and reduces skewness. The method suits analyses concerned with relative rather than absolute expression, and is common in both bulk and single-cell RNA-seq.

**Log shift followed by standard deviation scaling (`log_shift_scale_by_std`)** [1] A log transformation is combined with feature-wise scaling:

$$x'_{ij} = \frac{\log(x_{ij} + 1)}{\sigma_j} \tag{S5}$$

where $\sigma_j$ is the standard deviation of the log-transformed expression of gene $j$ across all cells. The logarithm reduces skewness, and each gene is then scaled to account for differences in variability, so that highly variable genes do not dominate downstream analyses. This matters most for methods that are sensitive to feature scaling, such as clustering or dimensionality reduction.

**Log shift with highly variable gene selection (`log_shift_size_factor_hvg`)** [1] Size-factor normalisation is extended here with a feature selection step based on gene variability. After log normalisation, only the most variable genes are kept:

$$x'_{ij} = \log\left(\frac{x_{ij}}{s_i} + 1\right), \quad j \in \mathcal{H} \tag{S6}$$

where $\mathcal{H}$ denotes the set of highly variable genes selected on their variance across cells. The number of selected genes is controlled by a parameter top_frac $\in (0, 1]$, which defines the fraction of genes with the highest variance to retain. For instance, top_frac $= 0.35$ corresponds to selecting the top 35% most variable genes. Dimensionality is reduced by keeping the genes that contribute most to variability in the dataset, which are often biologically informative. The transformation is ordinarily applied before clustering or dimensionality reduction, where irrelevant or low-variance genes may introduce noise.

**Log shift with z-score normalisation (`log_shift_size_factor_z`)** [1] Log-normalised data are further standardised across genes by z-score normalisation:

$$x'_{ij} = \frac{\log\left(\frac{x_{ij}}{s_i} + 1\right) - \mu_j}{\sigma_j} \tag{S7}$$

where $\mu_j$ and $\sigma_j$ are the mean and standard deviation of the log-normalised expression of gene $j$, respectively. Each gene then has zero mean and unit variance, so features are directly comparable. Downstream methods that assume standardised input, such as principal component analysis or most machine learning algorithms, require this step or an equivalent.

**Log shift, highly variable gene selection, and z-score normalisation** (`log_shift_hvg_z`) [1] Normalisation, feature selection and scaling are combined in a single pipeline:

$$x'_{ij} = \frac{\log\left(\frac{x_{ij}}{s_i} + 1\right) - \mu_j}{\sigma_j}, \quad j \in \mathcal{H} \tag{S8}$$

where $\mathcal{H}$ is the set of highly variable genes. The result is a compact, standardised representation of the data restricted to the most informative genes. It was designed for clustering and classification workflows that need dimensionality reduction and comparable features at once. As in the log shift with highly variable gene selection, the number of selected genes is controlled by a parameter top_frac $\in (0, 1]$, the fraction of genes with the highest variance to retain, which is set by default to 0.35.

**Scran normalisation** (`normalise_scran`) [1] This method approximates the size-factor philosophy of Lun *et al.* [26] using a global geometric-mean reference rather than the full pool-and-deconvolve estimator. Each cell's size factor is computed from the library sizes alone, and the normalised counts are then log-shifted:

$$x'_{ij} = \log\left(\frac{x_{ij}}{s_i} + 1\right) \tag{S9}$$

$$s_i = \frac{\ell_i}{\exp\left(\frac{1}{n}\sum_{k=1}^{n} \log \ell_k\right)} \tag{S10}$$

where $\ell_i$ is the total count of cell $i$. This is library-size scaling with size factors centred on a geometric mean of one. It does not carry out the pool-and-deconvolve estimation of scran and does not correct compositional bias; cells with zero counts are given a size factor of one.

**TMM normalisation** (`normalise_tmm`) [34] Trimmed Mean of M-values (TMM) normalisation aims to correct compositional biases between samples by comparing each cell with a reference:

$$M_{ij} = \log_2 \left( \frac{x_{ij}/s_i}{x_{rj}/s_r} \right) \tag{S11}$$

where $x_{ij}$ is the count of gene $j$ in cell $i$, $s_i$ is the library size (sum of counts) for cell $i$, and $r$ denotes the reference cell. The overall normalisation factor for each cell is a weighted mean of these M-values, taken after trimming the extreme 30% of values to reduce the influence of highly expressed genes or outliers. In the implementation a pseudocount of 0.5 is added to every count, so that no ratio is undefined, and the reference $r$ is the cell at the 75th percentile of library size. The M-values and the corresponding mean log-abundances are both trimmed to their central 40%, the remaining genes are weighted by their inverse asymptotic variance, and the factor of cell $i$ is $f_i = 2^{\overline{M}_i}$, rescaled so that the factors have a geometric mean of one. The transformed value is $x'_{ij} = \log(x_{ij}/f_i + 1)$. TMM stabilises normalisation in datasets with strong compositional differences and is the natural choice where a reference-based scheme is wanted. It is long established in RNA-seq and single-cell transcriptomics; Robinson and Oshlack give the full derivation [34].

**Box-Cox transformation** (`box_cox_transform`) [4] The Box-Cox transformation is a parametric method used to stabilise variance and approximate normality:

$$x'_{ij} = \begin{cases} \frac{(x_{ij}+c)^\lambda - 1}{\lambda}, & \lambda \neq 0 \\ \log(x_{ij} + c), & \lambda = 0 \end{cases} \tag{S12}$$

where $c$ is a shift ensuring positivity and $\lambda$ is estimated by maximum likelihood, separately for each gene. The shift is $c = 1 - \min_i x_{ij}$ when the gene has a non-positive value, which gives $c = 1$ for counts that include zeros, and $c = 0$ otherwise; a gene that is constant across cells is left unchanged. Each gene is transformed independently, so the transformation adapts to that gene's distribution. Its place in the pipeline is where downstream methods assume approximately Gaussian data, or where variance stabilisation is the overriding concern.

**Pearson residuals transformation** (`pearson_residuals_transform`) [1] Analytic Pearson residuals are computed under a negative binomial model:

$$x'_{ij} = \frac{x_{ij} - \mu_{ij}}{\sqrt{\mu_{ij} + \frac{\mu_{ij}^2}{\theta}}} \tag{S13}$$

where $\theta$ is the dispersion parameter and the expected count is $\mu_{ij} = \sum_k x_{ik} \sum_l x_{lj} / \sum_{k,l} x_{kl}$, the product of the cell and gene totals divided by the grand total. Residuals are clipped to $\pm\sqrt{n}$. Because the mean–variance relationship is modelled explicitly, the transformation removes technical noise while preserving biological variability. It performs well on highly sparse single-cell data and is now widely used in scRNA-seq workflows.

**GLM-PCA transformation** (`glm_pca_transform`) [1, 38] This method approximates a generalised linear model-based dimensionality reduction. The count matrix is first stabilised by size-factor normalisation and a log-transform, and a truncated singular value decomposition is then computed,

$$\begin{aligned} \tilde{X}_{ij} &= \log\left(\frac{x_{ij}}{s_i} + 1\right), \\ \tilde{X} &\overset{\text{SVD}}{=} U \Sigma V^\top, \qquad Z = U_k \Sigma_k, \end{aligned} \tag{S14}$$

where $s_i$ is the library size for cell $i$, $k$ is the requested number of components (`n_components`, default 50), and $Z$ is the resulting embedding. The dimensionality of single-cell RNA-seq data is reduced while the dominant directions of count variability are kept. The implementation is an explicit approximation to a full Poisson GLM-PCA [38]: it forgoes the iterative Fisher-scoring fit and keeps only the variance-stabilised projection. Where dimensionality reduction is needed early in the pipeline, it gives a compact embedding that clustering or visualisation routines can use directly, even on large datasets.

**SANITY-like transformation (`sanity_transform`)** [1] A Bayesian-inspired normalisation is obtained by introducing a data-driven pseudocount:

$$x'_{ij} = \log\left(\frac{x_{ij}}{s_i} + \epsilon\right) \tag{S15}$$

where $\epsilon = n/\sum_{i,j} x_{ij}$ is a heuristic pseudocount; the Bayesian Sanity model is not fitted.

A single global pseudocount stabilises low-expression values and reduces noise in sparse regions of the data, which suits the method to datasets dominated by zero or near-zero counts.

**Evaluation of Preprocessing Methods (`benchmark_transformations`)** The benchmark measures, quantitatively, how each normalisation or transformation method affects the statistical and biological properties of the dataset. The input data are first brought into a consistent numerical format and, when available, ground-truth cell-type labels are extracted for the biological evaluations. Each transformation is applied independently, and the transformed data are scored on several complementary metrics covering local and global structure and distributional properties. The metrics used for benchmarking are:

- **k-nearest neighbour (kNN) overlap** (default weight 0.35): the fraction of neighbours shared between the PCA representation of the original data and the PCA of the transformed data. For each cell, the k nearest neighbours are identified in both spaces, and the overlap fraction is calculated as

$$\text{Overlap} = \frac{\#\ \text{shared neighbours}}{k}. \tag{S16}$$

  High overlap means that the local structure of the data, that is, the relationships among similar cells, is preserved after transformation.
- **PCA variance preservation** (default weight 0.30): how similar the principal component loadings are before and after transformation. For each top PC, the absolute loadings in the original and transformed data are compared by Spearman correlation:

$$\mathrm{r}_i = \text{Spearman}\big(|\mathrm{PC}_i^{\text{orig}}|, |\mathrm{PC}_i^{\text{trans}}|\big) \tag{S17}$$

  and the final score is the average across the selected components. High values indicate that the global variance structure of the dataset is maintained.
- **Silhouette score** (default weight 0.25): how well the transformed cells cluster by known cell-type labels (when provided) or by unsupervised clusters derived from KMeans. For each cell, the score is computed as

$$s = \frac{b - a}{\max(a, b)}, \tag{S18}$$

  where $a$ is the average distance to other cells in the same cluster, and $b$ is the minimum average distance to cells in other clusters. Scores range from $-1$ to 1, and higher values indicate better separation between groups relative to cohesion within them.
- **Shapiro-Wilk normality fraction** (default weight 0.05): the fraction of genes whose expression distribution passes the Shapiro-Wilk normality test ($p > 0.05$). It shows whether the transformed data are close to Gaussian, as many downstream statistical analyses assume.
- **Anderson-Darling normality fraction** (default weight 0.05): as for Shapiro-Wilk, but with the Anderson-Darling test for normality, which gives a second check on distributional shape.

Three details of the implementation help in reading these metrics. The reference is the PCA of the matrix as supplied (the raw counts, when raw counts are given), so the first two metrics measure how much of the structure of the input a transformation keeps. For the transformations that change the columns (the two that keep only highly variable genes, and the GLM-PCA approximation, which returns components), the loading vectors are compared over their first $\min(p, p')$ entries; the PCA-preservation value is best read for the transformations that keep every gene. The two normality fractions are estimated from 50 features of the transformed matrix.

Each metric is normalised to the range $[0, 1]$ and the metrics are combined into a composite score with the default weights shown above. The weighted sum balances preservation of biological structure (kNN overlap, silhouette, PCA preservation) against statistically well-behaved data (normality tests), with most of the weight on the former. Optionally, a Borda-count rank aggregation can replace the weighted sum. Each metric is ranked independently across all transformations, and the ranks are then aggregated into a final ranking. No weights are needed. This is what one wants when the relative importance of the metrics is unclear and any choice of weights would be arbitrary. Benchmarking can also use bootstrap resampling to estimate the uncertainty of the composite score. Cells are sampled

**Supplementary table 8:** Data transformation benchmarking and preservation metrics implemented in scINTILLA.

| Method | Inputs | Outputs |
|---|---|---|
| Transformation Benchmark `benchmark_-transformations` | `data`: AnnData or DataFrame (`cell_type_col`: ground-truth for silhouette) (`scoring_method`: `"weighted"` or `"borda"`) (`weights`: custom metric importance) (`bootstrap_ci`: compute 95% CIs) (`n_pca_components`: default 20) | `results_df`: per-transform scores (kNN, Silhouette, Normality) `best_name`: name of top-ranked transformation `best_adata`: AnnData transformed with the winning method |

repeatedly with replacement, each transformation is applied, and the relevant metrics are recomputed. A composite score is calculated for each replicate with the default weights, and the 2.5th and 97.5th percentiles of these scores define a 95% confidence interval. The interval shows how stable a transformation's performance is under resampling, which its average score alone cannot show.

The outputs of the benchmarking procedure include:

- `results_df`: a pandas DataFrame containing per-transformation metrics (kNN overlap, PCA variance preservation, silhouette score, Shapiro-Wilk fraction, Anderson-Darling fraction) as well as the final composite score and optional bootstrap confidence intervals.
- `best_name`: the name of the transformation with the highest overall composite score.
- `best_adata`: the original dataset transformed using the top-ranked transformation, ready for downstream analysis or visualisation.

Inputs and outputs are listed in Supplementary Table 8.

**Normality Assessment** Normality in single-cell expression data is assessed with two complementary statistical tests, Shapiro–Wilk and Anderson–Darling. The Shapiro-Wilk test is applied to each gene on a subsample of cells (500 by default) to test whether its expression values follow a normal distribution. The subsample is kept small on purpose. At larger cell counts the test becomes extremely sensitive and may reject normality even for well-normalised data, leading to false negatives. Several subsampling replicates can be run to reduce variability, and the median fraction of features passing the test is recorded. The Anderson-Darling test, by contrast, is applied to the full feature vectors and compares each gene's distribution with a theoretical Gaussian, which gives an independent assessment of normality.

Normality is declared overall when a sufficient fraction of genes passes both tests, the fraction being set by a user-defined threshold. This is a practical compromise between statistical rigour and the inherent noise of single-cell measurements. For a more formal evaluation that accounts for multiple testing, the Shapiro-Wilk p-values can optionally be corrected with the Benjamini-Hochberg procedure; the proportion of genes that are significantly non-normal is then estimated with the false discovery rate controlled, in place of the heuristic threshold.

**Principal Component Analysis (PCA)** Principal Component Analysis is a dimensionality reduction technique that identifies orthogonal directions, known as principal components, which capture the maximum variance in the data.

In single-cell analysis, PCA projects the high-dimensional gene expression matrix onto a lower-dimensional space while preserving as much of the total variability as possible. The number of components can be specified a priori or chosen to explain a target proportion of the total variance.

Data-adaptive rules such as the Gavish–Donoho optimal hard threshold and the Marchenko–Pastur cutoff can also be used to choose the number of components that capture meaningful structure while limiting the influence of noise. They rely on the statistical properties of the singular values of the data matrix, and may use an estimate of the noise variance to refine the choice.

The resulting low-dimensional representation is stored within the dataset for downstream analyses such as clustering, visualisation and classification. The cumulative explained variance shows how much of the original signal is retained.

### S3 Feature Selection

Feature selection identifies the subset of genes that carry the most relevant biological signal for a given classification or clustering task, and is an important preprocessing step in single-cell transcriptomic analysis. In high-dimensional scRNA-seq data the full gene space is extremely sparse and noisy, and a large proportion of genes contribute little to the biological variation of interest. Reducing the feature space to a compact, informative subset improves the performance and interpretability of downstream classifiers; it also cuts computational cost and mitigates the curse of dimensionality.

scINTILLA implements five feature selection strategies, covering unsupervised variance-based, information-theoretic and ensemble-learning approaches. A benchmarking wrapper (`benchmark_feature_selection`) compares all of them on downstream classification performance, so that the strategy for a given dataset can be chosen on the data.

**PCA Loadings (`extract_top_genes_per_pc`)** Principal Component Analysis (PCA) is a linear dimensionality reduction technique that projects the original high-dimensional gene expression matrix into a lower-dimensional space defined by orthogonal directions of maximum variance. Each principal component (PC) is a linear combination of the original features, and the contribution of each gene to a given PC is encoded in its *loading coefficient*. Genes with large absolute loadings drive the variance captured by that component, and are therefore candidates for biologically relevant features.

`extract_top_genes_per_pc` uses this property for unsupervised feature selection. If PCA has already been computed and stored in the `AnnData` object (in `varm["PCs"]`), the precomputed loading matrix $\mathbf{L} \in \mathbb{R}^{p \times k}$ is used directly, where $p$ is the number of genes and $k$ the number of PCs. Otherwise, PCA is computed on the expression matrix $\mathbf{X} \in \mathbb{R}^{n \times p}$ via singular value decomposition, retaining up to $\min(50, n-1, p-1)$ components. For each PC, the top $n\_per\_pc$ genes are selected by ranking absolute loading values. The final gene set is the deduplicated union across all considered PCs. Because the approach is entirely unsupervised, it suits a first-pass filter, or settings where labels are unavailable. Two utility functions complement it: `build_reduced_dataset`, which subsets the `AnnData` object to the selected gene list, and `validate_reduced_set`, which measures the information retained by comparing classification accuracy on the full and the reduced feature space with a user-specified classifier (QDA, LDA or Random Forest).

**Highly Variable Genes (`select_hvg`)** Highly Variable Gene (HVG) selection is one of the most widely used unsupervised feature selection strategies in single-cell analysis. Its rationale is that genes with high cell-to-cell variability across the dataset are more likely to reflect biological differences, such as cell type identity or state, than technical noise, which tends to be spread more uniformly across genes.

`select_hvg` wraps three established HVG detection methods, all implemented in the Scanpy library.

*Seurat v3.* The mean–variance relationship of gene expression is modelled with a local regression (LOESS) fit. For each gene $g$, the expected variance is estimated as a function of the mean expression $\mu_g$, and a standardised variance $z_g$ is computed as:

$$z_g = \frac{\sigma_g^2}{\hat{\sigma}_g^2(\mu_g)} \tag{S19}$$

where $\hat{\sigma}_g^2(\mu_g)$ is the LOESS-predicted variance at mean $\mu_g$. The fit therefore describes a heteroscedastic mean–variance relationship; in the Scanpy implementation the variance is taken on values standardised by the predicted standard deviation and clipped at $\sqrt{n}$. Genes are ranked by $z_g$ and the top $n_{\text{top\_genes}}$ are selected. The `span` parameter controls the smoothness of the LOESS fit.

*Pearson residuals.* This variance-stabilising alternative is particularly suited to raw count data. Under a negative binomial model of gene expression, the Pearson residual for gene $g$ in cell $i$ is:

$$r_{ig} = \frac{x_{ig} - \mu_{ig}}{\sqrt{\mu_{ig} + \mu_{ig}^2/\theta}} \tag{S20}$$

where $\mu_{ig}$ is the expected count under the fitted model and $\theta$ is the overdispersion parameter. Genes whose residuals vary most across cells are selected as highly variable. The formulation accounts explicitly for the mean–variance dependence of count data, which should make it more robust than variance-only criteria in datasets with wide dynamic ranges.

*Cell Ranger.* A normalised dispersion criterion, originally developed for the 10x Genomics Cell Ranger pipeline, bins genes by mean expression and selects those with dispersion above the within-bin median.

If the `pearson_residuals` method fails (for instance because of library version incompatibilities), the function falls back automatically to `seurat_v3`. The output is an `AnnData` object filtered to the selected HVGs only.

**Mutual Information (`mi_feature_selection`)** Mutual Information (MI) is an information-theoretic measure of the statistical dependency between a gene's expression profile and a categorical target variable (e.g. cell type or disease label). Unlike correlation-based measures, MI captures non-linear as well as linear relationships and makes no assumptions about the underlying distribution of the data, which suits it to the non-Gaussian structure of single-cell expression data.

For a gene $g$ with expression values $X_g$ and a target variable $Y$, the mutual information is defined as:

$$I(X_g; Y) = \sum_{y \in \mathcal{Y}} \int p(x_g, y) \log \frac{p(x_g, y)}{p(x_g)\, p(y)} \, \mathrm{d}x_g \tag{S21}$$

In the discrete-continuous mixed setting typical of supervised feature selection in scRNA-seq, MI is estimated with a $k$-nearest-neighbour entropy estimator, as implemented in `sklearn.feature_selection.mutual_info_classif`. Each gene is scored independently, and the top $n_{\text{features}}$ genes ranked by $I(X_g; Y)$ are selected.

The method is supervised, using label information to find genes that predict the target. Its performance depends on how distinct the target classes are at the transcriptional level. One limitation is that genes are scored independently, so redundancy between selected features is not penalised: two genes carrying identical information about $Y$ would both receive high MI scores and both be selected. The mRMR method described below deals with this.

**Boruta (`boruta_selection`)** Boruta is a supervised, ensemble-based feature selection algorithm that treats feature relevance as a statistical hypothesis test. Instead of selecting a fixed number of top-ranked features, it tries to identify all features that carry more information about the target than chance would explain, which suits settings where the number of relevant genes is not known *a priori*.

The algorithm augments the original feature matrix with *shadow features* — permuted copies of each original gene expression vector. Because each shadow feature is constructed by independently shuffling the values of its corresponding original feature, the within-feature marginal distribution is preserved while any association with the target is destroyed. A Random Forest classifier is then trained on the augmented dataset $[\mathbf{X} \mid \mathbf{X}_{\text{shadow}}]$, and the importance of each real feature is compared against the maximum importance observed among the shadow features:

$$\tau = \max_j \ \operatorname{imp}(\tilde{X}_j) \tag{S22}$$

A real gene $g$ is considered relevant if $\operatorname{imp}(X_g) > \tau$. The procedure is repeated over several rounds, and a binomial hypothesis test is used to classify each gene as confirmed, rejected or tentative.

`boruta_selection` first attempts to use the `BorutaPy` package, which provides the full iterative statistical testing procedure. If the package is not available, a single-round shadow feature fallback (`_shadow_feature_fallback`) is run instead: shadow features are constructed by permuting each column independently, a Random Forest is trained on the augmented matrix, and real features whose importance exceeds the maximum shadow importance are selected. The fallback omits the iterative refinement and statistical testing of the full Boruta algorithm. It keeps the shadow-feature comparison and is cheaper to compute, but it is only an approximation. The output is a boolean mask indicating selected features, together with a per-feature importance score derived from the Random Forest rankings.

**mRMR (`mrmr_selection`)** Minimum Redundancy Maximum Relevance (mRMR) is a supervised feature selection criterion that maximises the relevance of each selected gene to the target variable while minimising the redundancy among the selected genes. It addresses a limitation of univariate filter methods such as MI-based selection, in which high-scoring genes may be mutually redundant and carry largely overlapping information about the target.

The mRMR score for a candidate feature $X_g$ given an already-selected set $\mathcal{S}$ is defined as:

$$\phi(X_g) = I(X_g; Y) - \frac{1}{|\mathcal{S}|} \sum_{X_j \in \mathcal{S}} I(X_g; X_j) \tag{S23}$$

where $I(X_g; Y)$ is the mutual information between gene $g$ and the target, and $I(X_g; X_j)$ is the pairwise mutual information between gene $g$ and each already-selected gene $X_j$. At each greedy step, the gene maximising $\phi$ is added to $\mathcal{S}$, so that the selected set progressively accumulates features that are individually informative but collectively diverse.

`mrmr_selection` first attempts to use the `mrmr` package, which provides an optimised implementation of the mRMR criterion for classification tasks. If the package is not available, a greedy MI-based approximation (`_greedy_mi_-mrmr`) is run instead. In the fallback, all features are pre-binned into 8 quantile bins (a choice motivated by the bimodal expression distributions commonly observed in scRNA-seq data), and pairwise MI values are computed on

**Supplementary table 9:** Summary of non-linear dimensionality-reduction methods exposed by scINTILLA.

| Function | Key parameters (defaults) |
|---|---|
| `run_umap` | `n_neighbors`=15, `min_dist`=0.5, `n_components`=2 |
| `run_tsne` | `perplexity`=30, `n_components`=2 |
| `run_diffusion_map` | `n_comps`=10 |
| `run_force_directed` | layout = ForceAtlas2 |
| `benchmark_embeddings` | methods $\subseteq$ {umap, tsne, diffmap} |

the binned representations to reduce computational cost. For large datasets, cells are also subsampled to at most 5,000 observations before MI estimation, a threshold at which MI estimates are empirically stable. The output consists of the selected feature indices in the order of selection (which is the greedy ranking) and their associated mRMR scores.

**Benchmark (`benchmark_feature_selection`)** `benchmark_feature_selection` compares the feature selection methods described above by their downstream predictive performance. The motivation is that no single feature selection strategy is best for every dataset: the relative performance of PCA loadings, MI, Boruta, HVG and mRMR depends on the dataset's dimensionality, class structure and noise characteristics. Instead of committing to one method *a priori*, scINTILLA lets the user identify empirically the most informative feature subset for their own data.

For each method in the specified list, $n_{\text{features}}$ genes are selected from the training split, a Logistic Regression classifier is trained on the reduced training set, and performance is evaluated on the held-out test set (80/20 stratified split). Performance is quantified by two metrics: overall accuracy and macro-averaged F1 score, the latter being less sensitive to class imbalance. Formally, for a selected feature index set $\mathcal{I}_m$ produced by method $m$:

$$\text{Acc}_m = \frac{1}{|\mathcal{D}_{\text{test}}|} \sum_{i \in \mathcal{D}_{\text{test}}} \mathbf{1}[\hat{y}_i = y_i]\,, \qquad F1_m = \frac{1}{C} \sum_{c=1}^{C} F1_c \tag{S24}$$

where $C$ is the number of classes and $F1_c$ is the per-class F1 score. Results are returned as a `DataFrame` for visual and quantitative comparison across methods. The function accepts an optional `AnalysisConfig` object, through which the list of methods and the number of features can be set in a central configuration instead of per-call arguments; this makes runs easier to reproduce.

A complementary function, `hvg_sensitivity_analysis`, tests how sensitive HVG-based feature selection is to the choice of $n_{\text{top\_genes}}$. It sweeps a user-defined range of values (default: $\{1000, 2000, 3000, 5000\}$), runs a full PCA $\rightarrow$ neighbourhood graph $\rightarrow$ clustering pipeline for each, and computes the Adjusted Rand Index (ARI) between the resulting clusters and the ground-truth labels. The stability of the optimal HVG count is assessed by comparing the best ARI with the median ARI across all runs: if the relative improvement falls within 5%, the selection is flagged as stable, meaning that downstream clustering is not materially sensitive to the precise choice of $n_{\text{top\_genes}}$.

### S4 Dimensionality Reduction

Beyond the linear PCA backbone used by upstream modules, scINTILLA exposes four non-linear and graph-theoretic embedding strategies through the `dimensionality_reduction` sub-package. Each operates on an existing high-dimensional representation (by default `X_pca` stored in `adata.obsm`), so embeddings stay comparable across the pipeline. Their inputs and outputs are summarised in Supplementary Table 9.

**UMAP (`run_umap`)** Uniform Manifold Approximation and Projection [29] models the data as a fuzzy topological simplicial complex built on the $k$-nearest-neighbour graph in the input representation. For each cell $i$ it defines a local connectivity radius $\rho_i$ and a smooth scale $\sigma_i$ such that the probability that $j$ is connected to $i$ is

$$p_{j|i} \;=\; \exp\!\left(-\frac{\max(0,\, d(x_i, x_j) - \rho_i)}{\sigma_i}\right), \tag{S25}$$

which is symmetrised to $p_{ij} = p_{j|i} + p_{i|j} - p_{j|i}\, p_{i|j}$. A low-dimensional embedding $\{y_i\}_{i=1}^{n}$ is then sought whose weights $q_{ij}(y) = (1 + a\|y_i - y_j\|^{2b})^{-1}$ approximate $p_{ij}$ in the sense of the binary cross-entropy

$$\mathcal{L}_{\text{UMAP}}(y) \;=\; \sum_{i \neq j} \left[ p_{ij} \log \frac{p_{ij}}{q_{ij}} + (1 - p_{ij}) \log \frac{1 - p_{ij}}{1 - q_{ij}} \right], \tag{S26}$$

where $a$ and $b$ are determined from the `min_dist` parameter, which controls how tightly local clusters are packed in the embedding. scINTILLA delegates the optimisation to `scanpy.tl.umap`, which calls the `umap-learn` implementation.

**t-SNE (`run_tsne`)** $t$-Distributed Stochastic Neighbour Embedding [27] matches pairwise affinities by minimising the Kullback–Leibler divergence between Gaussian neighbourhoods in the input space and Student-$t$ neighbourhoods in the embedding. For input vectors $\{x_i\}$ one defines

$$\begin{aligned} p_{j|i} &= \frac{\exp(-\|x_i - x_j\|^2/2\sigma_i^2)}{\sum_{k\neq i}\exp(-\|x_i - x_k\|^2/2\sigma_i^2)}, \\ p_{ij} &= \frac{p_{j|i} + p_{i|j}}{2n}, \end{aligned} \tag{S27}$$

with $\sigma_i$ set so that the perplexity $2^{-\sum_{j\neq i} p_{j|i}\log_2 p_{j|i}}$ matches the user-supplied `perplexity`. In the embedding,

$$q_{ij} = \frac{(1 + \|y_i - y_j\|^2)^{-1}}{\sum_{k\neq \ell}(1 + \|y_k - y_\ell\|^2)^{-1}}, \tag{S28}$$

and the embedding minimises $\mathrm{KL}(P\|Q) = \sum_{i\neq j} p_{ij}\log(p_{ij}/q_{ij})$ by gradient descent. scINTILLA uses the scanpy backend and falls back to `sklearn.manifold.TSNE` when scanpy is not available; for very small datasets the perplexity is clipped to $n - 1$.

**Diffusion Map (`run_diffusion_map`)** Diffusion maps [9] embed cells through the spectral decomposition of a Markov transition matrix on the kNN graph. Starting from an affinity kernel $W$ and its degree matrix $D$, the row-stochastic transition operator is $P = D^{-1}W$. The diffusion-map embedding at time $t$ retains the top $k$ non-trivial eigenpairs $(\lambda_\ell, \psi_\ell)$:

$$\Psi_t(x_i) = \big(\lambda_1^t\,\psi_1(i),\, \lambda_2^t\,\psi_2(i),\, \ldots,\, \lambda_k^t\,\psi_k(i)\big), \tag{S29}$$

so that Euclidean distance in $\Psi_t$ approximates the diffusion distance on the graph. The embedding suits continuous developmental trajectories and the detection of branching structure. The computation itself is done by `scanpy.tl.diffmap`, with $k$ = `n_comps` (default 10).

**Force-directed layout (`run_force_directed`)** A force-directed layout treats the graph as a mechanical system, with attractive springs between connected nodes and repulsive forces between all node pairs, and runs it to equilibrium; the resting positions are the embedding. scINTILLA uses the ForceAtlas2 [21] layout exposed through `scanpy.tl.draw_graph`, which operates on the same kNN graph constructed by `sc.pp.neighbors`. For sparse, branched topologies, where UMAP and diffusion maps may compress local structure too far, this layout can be the more informative view.

**Embedding Benchmark (`benchmark_embeddings`)** The wrapper `benchmark_embeddings` produces each requested embedding in turn and scores it against the high-dimensional reference with the local trustworthiness metric [40],

$$T(K) = 1 - \frac{2}{nK(2n - 3K - 1)}\sum_{i=1}^{n}\sum_{j\in U_K(i)}\big(r(i,j) - K\big), \tag{S30}$$

where $U_K(i)$ is the set of points that appear among the $K$ nearest neighbours of $i$ in the embedding but not in the input space, and $r(i,j)$ is the rank of $j$ in the input-space neighbourhood of $i$. It returns a `DataFrame` keyed by method, to be read alongside the qualitative views that the embedding functions produce. UMAP, t-SNE and the diffusion map are benchmarked in this way (the first two by default), each on its first two dimensions; the force-directed layout is meant for visual inspection and is not scored.

### S5 Clustering

This section gives the full description of the clustering module. Six of its families were run in this study and are summarised in the main Methods (Section 2.2); spectral, consensus and patient-level clustering were not run. Supplementary Tables 10 and 11 list the inputs and outputs of all clustering functions.

The analysis starts by evaluating several clustering strategies on the data, so that the one best suited to it can be chosen. scINTILLA implements nine clustering algorithms, covering partitional, hierarchical, density-based, graph-based, spectral and ensemble approaches; their inputs and outputs are listed in Supplementary Table 10. All of them sit inside one benchmarking framework (`benchmark_clustering_methods`), which scores each algorithm against ground-truth cell-type labels with the Adjusted Rand Index (ARI), so that the strategy is chosen from the data. The `unsupervised_analysis` dispatcher runs the whole pipeline (PCA preprocessing, clustering, evaluation and storage of the best-performing labels) in a single call; its inputs and outputs are listed in Supplementary Table 11.

**K-Means and Variants (`kmeans_clustering`)** K-Means [18] divides a dataset of $n$ cells into $K$ non-overlapping clusters by iteratively minimising the total within-cluster variance. Given an assignment of cells to cluster centroids $\boldsymbol{\mu}_1, \ldots, \boldsymbol{\mu}_K$, the objective function is:

$$\mathcal{J} = \sum_{k=1}^{K} \sum_{i \in \mathcal{C}_k} \|\mathbf{x}_i - \boldsymbol{\mu}_k\|^2 \tag{S31}$$

where $\mathcal{C}_k$ is the set of cells assigned to cluster $k$ and $\mathbf{x}_i \in \mathbb{R}^p$ is the feature vector of cell $i$. The algorithm alternates between an assignment step, in which each cell goes to its nearest centroid, and an update step, in which each centroid is recomputed as the mean of its assigned cells, until convergence.

`kmeans_clustering` offers three variants. **Standard K-Means** uses `k-means++` initialisation by default, which picks initial centroids with probability proportional to the squared distance from the centres already chosen. During benchmarking, `k-means++` and random initialisation are evaluated independently.

**Spherical K-Means** [19] deals with a known weakness of standard K-Means on high-dimensional sparse data such as scRNA-seq: Euclidean distance is sensitive to differences in total expression level, and these can outweigh the compositional differences between gene expression profiles. Each cell vector $\mathbf{x}_i$ is therefore L2-normalised before clustering:

$$\tilde{\mathbf{x}}_i = \frac{\mathbf{x}_i}{\|\mathbf{x}_i\|_2} \tag{S32}$$

so that all cells lie on the unit hypersphere and distances reflect angle, not magnitude.

**Bisecting K-Means** [37] is a divisive hierarchical variant. It reaches the $K$-cluster partition through a sequence of binary splits instead of a single global optimisation: at each step the cluster with the highest inertia is selected and split in two by K-Means with $K = 2$. This top-down scheme tends to give more balanced partitions than standard K-Means and is less sensitive to initialisation, but it does not optimise the global objective directly.

All three variants return the silhouette score and inertia alongside the cluster labels.

**Agglomerative Hierarchical Clustering (`hierarchical_clustering`)** Agglomerative hierarchical clustering [30] builds a nested hierarchy of partitions, starting from $n$ singleton clusters and repeatedly merging the two most similar until a single cluster holds all cells. Which clusters count as most similar depends on the *linkage method*, which defines the distance between two clusters as a function of the pairwise distances between their cells.

`hierarchical_clustering` supports four linkage strategies: *complete linkage*, which takes the maximum pairwise distance; *average linkage* (UPGMA), which takes the mean; *single linkage*, which takes the minimum; and *Ward linkage*, which minimises the total within-cluster variance at each merge and is restricted to Euclidean distance. Seven distance metrics are also supported (Euclidean, Cosine, Manhattan, Correlation, Chebyshev, Canberra and Braycurtis), and the resulting grid of method–metric combinations is evaluated during benchmarking, excluding Ward with a non-Euclidean metric, which is mathematically invalid.

Two computational backends are available. The fast `sklearn` mode uses `AgglomerativeClustering` from scikit-learn and is the one used during benchmarking, because it scales better. The `scipy` mode also computes the full linkage matrix $\mathbf{Z}$, draws the dendrogram and evaluates the *Cophenetic Correlation Coefficient* (CPCC), which measures how well the dendrogram preserves the pairwise distances in the original data:

$$\text{CPCC} = \frac{\sum_{i<j}(d_{ij} - \bar{d})(c_{ij} - \bar{c})}{\sqrt{\sum_{i<j}(d_{ij} - \bar{d})^2 \cdot \sum_{i<j}(c_{ij} - \bar{c})^2}} \tag{S33}$$

where $d_{ij}$ is the original pairwise distance between cells $i$ and $j$, $c_{ij}$ is the cophenetic distance, and $\bar{d}$, $\bar{c}$ are their respective means. A CPCC close to 1 means that the dendrogram represents the underlying distance structure faithfully.

**DBSCAN (`dbscan_clustering`)** Density-Based Spatial Clustering of Applications with Noise (DBSCAN) [14] treats clusters as contiguous regions of high point density separated by regions of low density. Unlike the partitional methods, it needs no number of clusters in advance, and points that fall in no cluster are labelled as noise with $-1$.

A cell $\mathbf{x}_i$ is a *core point* if at least `min_samples` cells lie within its $\varepsilon$-neighbourhood. Clusters are formed by connecting core points that are mutually density-reachable; border points (within $\varepsilon$ of a core point but not core points themselves) join the cluster of their nearest core point.

The two hyperparameters, $\varepsilon$ (neighbourhood radius) and `min_samples` (minimum neighbourhood size), are swept over a predefined grid during benchmarking. When `auto_eps=True`, $\varepsilon$ is instead estimated from the data by the *k-distance graph elbow method*, which gives a data-adaptive $\varepsilon$ that need not be tuned by hand. Because the elbow is itself an uncertain estimate, the benchmark also evaluates $\varepsilon \times 0.8$ and $\varepsilon \times 1.2$.

**HDBSCAN (`hdbscan_clustering`)** Hierarchical DBSCAN (HDBSCAN) [28] extends DBSCAN by building a full hierarchy of density-based clusters and extracting the most stable partition from it. In place of a fixed $\varepsilon$, it works on *mutual reachability distances*, a transformation of the pairwise distances. A minimum spanning tree is built on the mutual reachability graph and condensed into a cluster hierarchy, from which clusters are extracted by selecting the subtrees that maximise a persistence (stability) criterion. HDBSCAN can therefore handle clusters of varying density, and $\varepsilon$ no longer has to be specified.

`hdbscan_clustering` runs a grid search over `min_cluster_size` (the minimum number of cells for a group to be considered a cluster) and `min_samples` (which sets how conservative the core-point definition is). Each combination is scored as the number of clusters minus the noise fraction, and the highest-scoring one is kept. Because the first term is an integer and the second lies below one, the rule favours the finest partition in the grid whatever its noise. Only this partition enters the clustering benchmark, which is why HDBSCAN returned no partition with at most 50% of cells as noise for the lung and HNOCA atlases. Both the dedicated `hdbscan` package and the `sklearn` implementation (available from scikit-learn $\geq$ 1.3) are supported; the former is used when it is available.

**Leiden Clustering (`leiden_clustering`)** [39] Leiden is a community-detection algorithm that works on a $k$-nearest-neighbour (kNN) graph of cells, in which each cell is joined to its $k$ most similar neighbours in the PCA embedding space. Communities are found by optimising the *modularity* objective, which measures the difference between the observed density of edges within communities and the density expected under a null model:

$$Q(\gamma) = \frac{1}{2m} \sum_{i,j} \left[ A_{ij} - \gamma \frac{k_i k_j}{2m} \right] \delta(c_i, c_j), \tag{S34}$$

where $A_{ij}$ is the adjacency matrix of the kNN graph, $k_i$ and $k_j$ are the degrees of nodes $i$ and $j$, $m$ is the total number of edges, $c_i$ is the community assignment of cell $i$, and $\delta$ is the Kronecker delta. Leiden improves on the earlier Louvain method by guaranteeing that communities are internally connected, so poorly connected or disconnected partitions cannot form. The resolution parameter $\gamma$ scales the null-model term and sets the granularity of the partition: higher values give finer communities and lower values coarser ones. scINTILLA sweeps a resolution grid from 0.1 to 3.0 during benchmarking. Two other ways of choosing the resolution are available: an *adaptive bisection search* (`adaptive_resolution=True`), which refines the resolution around the ARI-maximising value with a two-pass bisection, and an *NVI stability criterion* (`resolution_selection="nvi_stability"`), which picks the resolution plateau of maximum stability in the Normalised Variation of Information curve. The second needs no ground-truth labels and is fully unsupervised.

**Louvain Clustering (`louvain_clustering`)** [3] Louvain optimises the same modularity objective on the kNN graph as Leiden. It iterates two phases until convergence: in the first phase, each cell is moved to the neighbouring community that gives the largest increase in modularity; in the second, the graph is coarsened by collapsing each community into a single node, with edge weights preserved. `louvain_clustering` is implemented through Scanpy's `sc.tl.louvain` interface and sweeps the same resolution grid as Leiden (0.1–3.0) during benchmarking. It is there as a comparator for Leiden: running both shows whether Leiden's connectivity guarantee translates into a higher ARI on the datasets under study.

**Spectral Clustering (`spectral_clustering`)** [41] Spectral clustering embeds cells in a low-dimensional space defined by the eigenvectors of the graph Laplacian and applies K-Means in that space. Given an affinity matrix $\mathbf{W}$, the normalised graph Laplacian is:

$$\mathbf{L}_{\text{sym}} = \mathbf{D}^{-1/2}(\mathbf{D} - \mathbf{W})\mathbf{D}^{-1/2} = \mathbf{I} - \mathbf{D}^{-1/2}\mathbf{W}\mathbf{D}^{-1/2} \tag{S35}$$

where $\mathbf{D}$ is the diagonal degree matrix with $D_{ii} = \sum_j W_{ij}$. The $K$ eigenvectors belonging to the $K$ smallest eigenvalues of $\mathbf{L}_{\text{sym}}$ form the spectral embedding, and K-Means on this embedding gives the final partition.

The method is well suited to non-convex cluster shapes, which methods based on Euclidean distance cannot capture. `spectral_clustering` supports a single-run mode (fixed $K$) and a grid-search mode, in which a range of $K$ values is evaluated and the partition with the highest silhouette score is returned.

**Consensus Clustering (`consensus_clustering`)** Consensus clustering is an ensemble approach: it combines the outputs of many clustering runs into one partition. The algorithm constructs a *consensus matrix* $\mathbf{M} \in [0,1]^{n\times n}$, where entry $M_{ij}$ records the fraction of runs in which cells $i$ and $j$ were assigned to the same cluster:

$$M_{ij} = \frac{1}{R}\sum_{r=1}^{R} \mathbf{1}\left[\ell_i^{(r)} = \ell_j^{(r)}\right] \tag{S36}$$

where $R$ is the total number of runs and $\ell_i^{(r)}$ is the label of cell $i$ in run $r$. Values of $M_{ij}$ close to 1 mean that cells $i$ and $j$ are consistently co-clustered, and values close to 0 that they are consistently separated. The consensus matrix is converted to a distance matrix $\mathbf{D} = \mathbf{1} - \mathbf{M}$, and the final partition is obtained by average-linkage hierarchical clustering, with the dendrogram cut at the desired number of clusters.

`consensus_clustering` accepts K-Means, Leiden and Spectral as base clustering methods, with $n_{\text{runs\_per\_method}}$ parameter variations per method. For K-Means and Spectral the parameter varied across runs is the number of clusters $K$; for Leiden it is the resolution. Each base method's stability is the average ARI over all pairs of its runs, and this per-method reliability score is returned with the final consensus labels. The number of clusters for the hierarchical cut of the consensus matrix comes from a square-root heuristic: $K_{\text{consensus}} = \max\Big(2,\ \lfloor\sqrt{n/2}\rfloor\Big)$.

All of the methods above are evaluated in parallel through one benchmarking framework, which ranks every configuration by its agreement with the ground-truth cell-type annotations.

**Benchmark and Pipeline (`benchmark_clustering_methods`, `unsupervised_analysis`)** For each method and hyperparameter configuration, cluster labels are computed and compared with the ground-truth cell-type annotations using the Adjusted Rand Index (ARI) [20]:

$$\text{ARI} = \frac{\text{RI} - \mathbb{E}[\text{RI}]}{\max(\text{RI}) - \mathbb{E}[\text{RI}]} \tag{S37}$$

where RI is the Rand Index and the expectation is taken under the hypergeometric model of random partitions. ARI ranges from $-1$ to 1: 1 is perfect agreement, 0 the agreement expected by chance, and negative values agreement worse than chance. Cells labelled as noise are excluded from the ARI; a configuration that classifies more than 50% of cells as noise is given a `NaN` ARI, so that degenerate solutions are not rewarded. Density-based methods are thus scored on their core cells only, which favours them slightly; their partitions are otherwise ranked by the same ARI. Results come back as a `DataFrame` with columns `method`, `params`, `ari`, `n_clusters` and `noise_fraction`, together with a summary bar chart.

The `unsupervised_analysis` dispatcher is the single entry point to the whole unsupervised pipeline. PCA is optionally run first (default: 30 components) to compute the low-dimensional representation that every clustering method takes as input. After benchmarking, the method with the highest ARI is selected and, when `store_-labels=True`, its cluster labels are written to `adata.obs["scintilla_cluster"]` for downstream analyses. An optional `AnalysisConfig` object lets the list of clustering methods, the resolution grids and the HDBSCAN parameter ranges be set in one place, which makes an analysis easier to configure and to reproduce.

**Patient-Level Clustering (`build_per_celltype_clustering`, `patient_level_clustering`)** scINTILLA also clusters patients, stratifying them by their transcriptional profiles across cell types. In disease cohorts, patients differ at the single-cell level; they also differ in the relative composition and expression of their cell populations. The framework has three stages. First, `build_per_celltype_clustering` averages each patient's single-cell expression within each cell type, giving one pseudobulk profile per patient–cell-type combination. Patients are then clustered independently within each cell type, by agglomerative hierarchical clustering (default: Canberra distance with complete linkage) or, as a fallback, K-Means. A patient with no cells of a given type receives the sentinel label $-1$ for that type.

Next, `build_similarity_matrices` constructs a binary co-clustering similarity matrix for each cell type: entry $(i,j)$ is 1 if patients $i$ and $j$ fell in the same cluster within that cell type, and 0 otherwise. A pair in which either patient lacks the cell type is not counted as co-clustered. `weighted_assembly` then combines these matrices into one aggregate similarity matrix by taking a weighted average across cell types; user-supplied weights set how much each cell type contributes to the final patient similarity.

Finally, `patient_level_clustering` converts the aggregate similarity matrix into a distance matrix ($D_{ij} = 1 - S_{ij}$) and derives the final patient partition. In the default `mds_kmeans` mode, Multi-Dimensional Scaling (MDS) first embeds the patients in a low-dimensional Euclidean space that preserves their pairwise dissimilarities, and K-Means is run on that embedding. Applying K-Means directly to the non-Euclidean similarity matrix would be unstable. The two-step route avoids this, and is intended to give more interpretable cluster boundaries.

**Supplementary table 10:** Summary of clustering methods implemented in scINTILLA, including their required inputs, optional parameters (shown in parentheses), and outputs.

| Method | Inputs | Outputs |
|---|---|---|
| K-Means & variants<br>`kmeans_clustering` | `data`: expression matrix or AnnData<br>`n_clusters`: number of clusters $K$<br>(`init`: `k-means++` or `random`)<br>(`spherical`: L2-normalise rows)<br>(`bisecting`: use BisectingKMeans) | `labels`: cluster label array ($n$)<br>`model`: fitted sklearn model<br>`metrics`: dict with `inertia` and `silhouette` |
| Agglomerative Hierarchical<br>`hierarchical_clustering` | `data`: expression matrix or AnnData<br>`n_clusters`: number of clusters<br>(`metric`: `euclidean`, `cosine`, `manhattan`, `correlation`, `chebyshev`, `canberra`, `braycurtis`)<br>(`linkage`: `complete`, `average`, `single`, `ward`)<br>(`mode`: `sklearn` or `scipy`) | *sklearn*: `labels` array ($n$)<br>*scipy*: `labels` array ($n$)<br>linkage matrix $\mathbf{Z}$<br>CPCC score<br>dendrogram figure |
| DBSCAN<br>`dbscan_clustering` | `data`: expression matrix or AnnData<br>(`eps`: neighbourhood radius $\varepsilon$)<br>(`min_samples`: minimum neighbourhood size)<br>(`metric`: distance metric) | `labels`: array ($n$, $-1$ = noise)<br>`n_clusters`: number of clusters<br>`n_noise`: number of noise points<br>`silhouette`: score (non-noise) |
| HDBSCAN<br>`hdbscan_clustering` | `data`: expression matrix or AnnData<br>(`min_cluster_size_range`)<br>(`min_samples_range`)<br>(`metric`: distance metric) | `results_df`: DataFrame<br>`best_labels`: array ($n$) |
| Leiden<br>`leiden_clustering` | `data`: AnnData (PCA in `obsm`)<br>(`resolution`: $\gamma$)<br>(`use_rep`: embedding key) | `labels`: array ($n$) |
| Louvain<br>`louvain_clustering` | `data`: AnnData (PCA in `obsm`)<br>(`resolution`: $\gamma$)<br>(`use_rep`: embedding key) | `labels`: array ($n$) |
| Spectral<br>`spectral_clustering` | `data`: expression matrix or AnnData<br>(`n_clusters`)<br>(`affinity`: `rbf`)<br>(`n_clusters_range`) | single: `labels`, model, metrics<br>grid: `results_df`, `best_labels` |
| Consensus<br>`consensus_clustering` | `adata`: AnnData<br>(`methods`: default `kmeans`)<br>(`n_runs_per_method`)<br>(`resolution_range`) | `consensus_matrix`: $\mathbf{M}$<br>`consensus_labels`: array ($n$)<br>`stability_scores` |
| Patient-level clustering<br>`build_per_celltype_-clustering`<br>`build_similarity_matrices`<br>`weighted_assembly`<br>`patient_level_clustering` | `data`: AnnData<br>`patient_col`<br>`celltype_col`<br>(`n_clusters`)<br>(`method`)<br>(`metric`, `linkage`)<br>(`weights`) | `clustering_results`<br>`similarity_matrices`<br>`combined_matrix`<br>`patient_labels` |

**Supplementary table 11:** High level functions for unsupervised analysis implemented in scINTILLA.

| Method | Inputs | Outputs |
|---|---|---|
| Benchmark<br>`benchmark_clustering_-`<br>`methods` | `data`: AnnData<br>`cell_type_col`<br>(`n_clusters`)<br>(`use_rep`)<br>(`adaptive_resolution`)<br>(`auto_eps`)<br>(`config`) | `results_df`<br>`labels_dict`<br>summary figure |
| Pipeline<br>`unsupervised_analysis` | `data`: AnnData or DataFrame<br>`cell_type_col`<br>(`n_clusters`)<br>(`run_pca_first`)<br>(`n_pca_comps`)<br>(`store_labels`)<br>(`config`) | `adata` (updated)<br>`results_df`<br>`labels_dict`<br>`fig`<br>`best_method` |

**Supplementary table 12:** Summary of classification methods implemented in scINTILLA, including their required inputs, optional parameters (shown in parentheses), and outputs.

| Method | Inputs | Outputs |
|---|---|---|
| Linear Discriminant Analysis `lda_classification` | `X_train, X_test`: feature matrices `y_train, y_test`: label arrays | `model`: fitted LDA model `metrics`: dict (accuracy, F1, etc.) |
| Quadratic Discriminant Analysis `qda_classification` | `X_train, X_test, y_train, y_test` (`reg_param`: 0.01) | `model`: fitted QDA model `metrics`: dict with performance scores |
| Support Vector Machine `svm_classification` | `X_train, X_test, y_train, y_test` (`max_iter`: 2000) (`random_state`: `RANDOM_SEED`) | `model`: fitted `LinearSVC` model `metrics`: dict with performance scores |
| Random Forest `random_forest_classification` | `X_train, X_test, y_train, y_test` (`n_estimators`: 100) (`random_state`: `RANDOM_SEED`) | `model`: fitted RandomForest model `metrics`: dict with performance scores |
| Logistic Regression `logistic_regression_classification` | `X_train, X_test, y_train, y_test` (`max_iter`: 500) (`solver`: `lbfgs`) | `model`: fitted LogReg model `metrics`: dict with performance scores |
| Multi-Layer Perceptron `mlp_classification` | `X_train, X_test, y_train, y_test` (`hidden_layer_sizes`: (100, 50)) (`max_iter`: 500) | `model`: fitted MLP model `metrics`: dict with performance scores |
| k-Nearest Neighbours `knn_classification` | `X_train, X_test, y_train, y_test` (`n_neighbors`: 5) | `model`: fitted kNN model `metrics`: dict with performance scores |
| XGBoost / LightGBM `xgboost_classification` `lightgbm_classification` | `X_train, X_test, y_train, y_test` (`eval_metric`: `mlogloss`) (`verbosity`: 0 or -1) | `model`: fitted gradient boosting model `metrics`: dict (requires XGBoost/LightGBM) |
| Gradient Boosting (Hist) `gradient_boosting_classification` | `X_train, X_test, y_train, y_test` (`random_state`: `RANDOM_SEED`) | `model`: fitted HistGBM model `metrics`: dict with performance scores |
| Naive Bayes `naive_bayes_classification` | `X_train, X_test, y_train, y_test` | `model`: fitted GaussianNB model `metrics`: dict with performance scores |
| Stacking Ensemble `stacking_ensemble_classification` | `X_train, X_test, y_train, y_test` (`estimators`: LogReg, RF, `LinearSVC`) (`final_estimator`: LogReg) (`cv`: 5) | `model`: fitted Stacking model `metrics`: dict with performance scores |

### S6 Classification

This section gives the full description of the classification module. All twelve classifiers were run in this study and are summarised in the main Methods (Section 2.3); the feature-importance module at the end of the section was not used. Supplementary Tables 12 and 13 list the inputs and outputs of the classification functions.

The supervised analysis starts by comparing several classification strategies, judged on both predictive accuracy and interpretability. scINTILLA implements up to twelve classifiers: generative models, discriminative linear models, and ensemble or neural approaches. Their inputs and outputs are listed in Supplementary Table 12. All methods sit inside one benchmarking framework (`benchmark_models_comprehensive`), which scores each algorithm by the macro-averaged F1-score so that the best-discriminating model in each feature space can be chosen from the data. The chosen model has two uses: it assigns labels to future unlabelled observations, and, through SHAP-based importance ranking, it shows how much each feature contributes to the decision. All twelve classifiers were run in this study. The full pipeline is called through the `supervised_analysis` dispatcher, which handles data normalisation, stratified train-test splitting, model training, cross-validation and, optionally, SHAP importance scores in a single call. Its high-level entry points are listed in Supplementary Table 13.

**LDA Classification (`lda_classification`)** Linear Discriminant Analysis (LDA) [15] finds a linear combination of features that separates two or more classes. It projects the features from a high-dimensional space onto a lower-dimensional one while maximising the ratio of between-class to within-class variance.

The data are assumed to follow a multivariate normal distribution $X \sim N(\mu, \Sigma)$. The decision boundary is determined by the linear score function:

$$\delta_k(x) = x^T \Sigma^{-1} \mu_k - \frac{1}{2} \mu_k^T \Sigma^{-1} \mu_k + \log \pi_k \tag{S38}$$

where $\mu_k$ is the class mean, $\Sigma$ is the shared covariance matrix, and $\pi_k$ is the prior probability of class $k$. Maximising this function partitions the feature space into regions with linear boundaries, which keeps the model cheap to fit and its decisions easy to read off, for binary and multiclass problems alike. It is implemented through the function `LinearDiscriminantAnalysis` from `sklearn.discriminant_analysis`.

**QDA Classification (`qda_classification`)** Quadratic Discriminant Analysis [15] relaxes the LDA assumption of homoscedasticity by giving each class $k \in \{1, \ldots, K\}$ its own covariance matrix $\Sigma_k$. The model can then capture second-order interactions between features, and its decision boundaries become quadratic. Deriving the posterior probability from the multivariate Gaussian density gives the quadratic discriminant score:

$$\begin{aligned} Q_k(x) = & -\frac{1}{2} \log |\Sigma_k| - \frac{1}{2} (x - \mu_k)^T \Sigma_k^{-1} (x - \mu_k) \\ & + \log \pi_k + \text{const}, \end{aligned} \tag{S39}$$

where the additive constant collects terms that do not depend on the class index $k$ and hence do not affect the arg max.

When there are many features relative to observations, the per-class covariance estimates have high variance. A regularisation parameter such as `reg_param` shrinks them towards a common average, which places the model between the rigidity of LDA and the full flexibility of QDA. It is implemented through the function `QuadraticDiscriminantAnalysis` from `sklearn.discriminant_analysis`.

**SVM Classification (`svm_classification`)** Support Vector Machines [10] are maximum-margin classifiers that determine a separating hyperplane $f(x) = w^\top x + b$ between distinct classes. scINTILLA adopts the linear formulation implemented by `LinearSVC` (`sklearn.svm`), which scales to the tens of thousands of features and tens of thousands of cells that are typical of single-cell datasets. Multi-class problems are handled internally through a one-vs-rest decomposition.

The primal optimisation problem solved by `LinearSVC` is the soft-margin programme

$$\begin{aligned} \min_{w,\, b,\, \xi} \ & \frac{1}{2} \|w\|^2 + C \sum_{i=1}^{n} \xi_i^2, \\ \text{s.t. } & y_i (w^\top x_i + b) \geq 1 - \xi_i, \quad \xi_i \geq 0, \end{aligned} \tag{S40}$$

with squared-hinge loss as the default surrogate. The hyperparameter $C$ governs the trade-off between margin width and slack-variable mass, hence between bias and variance. `LinearSVC` returns uncalibrated decision-function scores, not calibrated posteriors, and its one-vs-rest margins are not comparable across classes, so AUROC is not reported for it; probability-calibrated metrics that rely on Platt scaling or isotonic regression [31] are left out on purpose, so that calibration uncertainty is not counted twice inside the benchmark. The wrapper sets `max_iter`=2000, to help convergence on the higher-dimensional gene-space inputs, and `random_state`=`RANDOM_SEED` for reproducibility.

**Random Forest Classification (`random_forest_classification`)** A Random Forest [6] builds many decision trees $\{T_1(x), \ldots, T_M(x)\}$ by Bootstrap Aggregating (bagging). Randomness in both data sampling and feature selection decorrelates the trees, which reduces model variance without a proportional increase in bias. The final classification $\hat{C}$ for a given input $x$ follows the soft-vote rule, in which class probabilities estimated by each tree are averaged across the ensemble:

$$\hat{C}(x) \ = \ \arg\max_k \ \frac{1}{M} \sum_{m=1}^{M} \hat{p}_m(y = k \mid x), \tag{S41}$$

where $\hat{p}_m(y = k \mid x)$ denotes the class-$k$ probability returned by tree $T_m$ and $M$ is the value of `n_estimators`. The scikit-learn implementation in `RandomForestClassifier` uses this soft-vote convention, which is generally preferred to hard-vote majority counting because it keeps the trees' uncertainty. The ensemble is comparatively insensitive to outliers and handles high-dimensional data with non-linear interactions between variables without requiring explicit feature scaling. It is implemented through the function `RandomForestClassifier` from `sklearn.ensemble`.

**Logistic Regression Classification** (`logistic_regression_classification`) Logistic Regression [12] estimates the probability of a categorical dependent variable from one or more independent features. Unlike linear regression, it uses the logistic sigmoid to map any real number to a probability between 0 and 1. The model assumes that the log-odds of the event are linear in the predictors:

$$\log\left(\frac{p}{1-p}\right) = w^T x + b \tag{S42}$$

where $p = P(y = 1|x)$. The optimal parameters $w$ and $b$ are usually found by Maximum Likelihood Estimation (MLE), that is, by minimising the cross-entropy loss. In the implementation, the `lbfgs` solver performs the optimisation iteratively, and the maximum number of iterations (`max_iter`) can be set as an input (by default = 500). Because the model returns probabilities that are generally well calibrated [31], and because its coefficients carry a direct interpretation, it remains a natural first choice wherever the influence of individual features has to be understood rather than merely exploited. It is implemented through the function `LogisticRegression` from `sklearn.linear_model`.

**Multi Layer Perceptron Classification** (`mlp_classification`) The Multi-Layer Perceptron [35] is a feedforward neural network that models a non-linear mapping from input features to target outputs. It has an input layer, one or more hidden layers, and an output layer, where each neuron applies a non-linear activation function $\sigma(\cdot)$ to a weighted sum of its inputs. Indexing layers by $l = 1, \ldots, L$, the transformation reads

$$h^{(l)} \;=\; \sigma\Big(W^{(l)} h^{(l-1)} + b^{(l)}\Big)\,, \qquad h^{(0)} \equiv x, \tag{S43}$$

where $W^{(l)}$ and $b^{(l)}$ are the layer-$l$ weights and biases, and the network output $\hat{y} = h^{(L)}$ is passed through a softmax for multi-class classification. Here the network has two hidden layers of 100 and 50 neurons, set by `hidden_layer_sizes`. It is trained by backpropagation, which computes the gradient of the loss with respect to each weight by the chain rule. With `max_iter=500`, the stochastic optimiser has up to 500 iterations to minimise the cross-entropy error. The architecture can capture structure in high-dimensional data that linear classifiers miss. It is implemented through the function `MLPClassifier` from `sklearn.neural_network`.

**K-Nearest Neighbours Classification** (`knn_classification`) The k-nearest neighbours algorithm [11] is non-parametric and instance-based: it classifies a point from the closest training examples in feature space. It estimates no parameters during training and relies instead on local density estimation. A query point $x$ takes the majority label of its neighbours, with distance usually measured by the Minkowski metric, a generalisation of the Euclidean distance:

$$D(x, x') = \left(\sum_{j=1}^{d} |x_j - x'_j|^q\right)^{1/q} \tag{S44}$$

In the function, the parameter `n_neighbors` (set to $k = 5$) governs the bias–variance trade-off: smaller values of $k$ capture fine-grained local structure but are sensitive to noise, whereas larger values give smoother decision boundaries. Class probabilities are the proportions of the $k$ neighbours in each class, and these allow ranking-based metrics to be computed within the `_fit_and_eval` pipeline. It is implemented through the function `KNeighborsClassifier` from `sklearn.neighbors`.

**Gradient Boosting Classification** (`gradient_boosting_classification`) The Histogram-based Gradient Boosting Classifier [24] is a faster variant of the gradient boosting decision tree (GBDT) algorithm [16]: it bins continuous input features into discrete integer-valued bins. Binning cuts the cost of finding the best split point from $O(n_{\text{samples}})$ to $O(n_{\text{bins}})$, so training stays fast on large datasets. The model minimises a loss function $L(y, F(x))$ by sequentially adding weak learners $h_t(x)$ that fit the functional gradient $g_i$ of the loss and are then subtracted, so that at iteration $t$

$$\begin{aligned} F_t(x) \;&=\; F_{t-1}(x) - \eta\, h_t(x), \\ h_t &\approx \arg\min_{h\in\mathcal{H}} \sum_{i=1}^{n} (g_i - h(x_i))^2\,, \end{aligned} \tag{S45}$$

where $\eta$ is the learning rate, $g_i = [\partial L(y_i, F)/\partial F]_{F=F_{t-1}(x_i)}$ is the pointwise gradient evaluated at the previous ensemble, and $\mathcal{H}$ is the family of regression trees fitted on the binned feature space. In the implementation, `HistGradientBoostingClassifier` runs in parallel on multiple cores and handles missing values (NaNs) natively, sending them to whichever branch minimises the loss during training. It trades little accuracy for a considerable gain in speed. That matters when it is one of a dozen models fitted inside an automated evaluation pipeline. It is implemented through the function `HistGradientBoostingClassifier` from `sklearn.ensemble`.

**Naive Bayes Classification (`naive_bayes_classification`)** Gaussian Naive Bayes [33] is a probabilistic classifier based on Bayes' Theorem under the *naive* assumption of conditional independence between every pair of features given the class label. For a given class $y$ and feature vector $x = (x_1, \ldots, x_n)$, the model computes the posterior probability:

$$P(y|x_1, \ldots, x_n) \propto P(y) \prod_{i=1}^{n} P(x_i|y) \tag{S46}$$

In the Gaussian variant, the conditional density $P(x_i|y)$ is modelled using a normal distribution characterised by the class-specific mean $\mu_{y,i}$ and variance $\sigma^2_{y,i}$. The classification rule selects the class that maximises this product, known as the Maximum A Posteriori (MAP) decision rule.

In scINTILLA, `GaussianNB` is a cheap baseline. It is simple to compute and needs little training data, because it estimates only the first two moments (mean and variance) of each feature's distribution. The independence assumption is rarely satisfied in practice, yet GNB holds up tolerably well on high-dimensional problems where linear decision boundaries suffice. It is implemented through the function `GaussianNB` from `sklearn.naive_bayes`.

**Stacking Ensemble Classification (`stacking_ensemble_classification`)** Stacking (stacked generalisation) trains a meta-model to combine the predictions of several heterogeneous base estimators [42, 5]. Where averaging or voting simply pools them, stacking treats the outputs of the individual classifiers as new features in a higher-level feature space. Given $K$ base learners $\{f_1, \ldots, f_K\}$, the ensemble output is defined by a meta-classifier $\Phi$:

$$\hat{y} = \Phi(f_1(x), f_2(x), \ldots, f_K(x)) \tag{S47}$$

The `StackingClassifier` uses Logistic Regression, a Random Forest with `n_estimators`=30 (the standalone wrapper uses 100), and a `LinearSVC` as base learners. The meta-model therefore receives the decision values of the `LinearSVC` beside the class probabilities of the other two. To limit overfitting, training uses 5-fold cross-validation (`cv=5`): the meta-model is fitted on out-of-fold predictions, so the final Logistic Regression estimator weights each base model by how well it generalises. The two-level arrangement is intended to capture relationships that a single model would miss and to give more stable predictions. It is implemented through the function `StackingClassifier` from `sklearn.ensemble`.

**XGBoost Classification (`xgboost_classification`)** XGBoost (Extreme Gradient Boosting) [8] is a gradient boosting library built for speed on large datasets. It adds decision trees to the ensemble one at a time, each trained to predict the residuals (errors) of the cumulative prediction from all previous trees. The model minimises a regularised objective function:

$$\text{Obj}^{(t)} = \sum_{i=1}^{n} l(y_i, \hat{y}_i^{(t-1)} + f_t(x_i)) + \Omega(f_t) \tag{S48}$$

where $l$ is a convex loss function and $\Omega(f_t)$ penalises model complexity to limit overfitting. With a second-order Taylor expansion of the loss function around the previous prediction $\hat{y}_i^{(t-1)}$,

$$\begin{aligned} \mathcal{L}^{(t)} &\approx \sum_{i=1}^{n} \left[ g_i f_t(x_i) + \tfrac{1}{2} h_i f_t(x_i)^2 \right] + \Omega(f_t), \\ g_i &= \partial_{\hat{y}} \ell, \quad h_i = \partial^2_{\hat{y}} \ell, \end{aligned} \tag{S49}$$

XGBoost obtains closed-form leaf-weight updates and a tractable split-gain criterion, and these account for its faster convergence and greater flexibility compared with first-order gradient boosting. In the wrapper, `LabelEncoder` converts the labels to the form expected by the underlying C++ implementation, and the `mlogloss` metric points the optimisation at the cross-entropy of the multi-class predictions. It is implemented through the function `XGBClassifier` from `xgboost`.

**LightGBM Classification (`lightgbm_classification`)** LightGBM [24] is another gradient boosting framework. It reduces training time and memory use with two techniques, Gradient-based One-Side Sampling (GOSS) and Exclusive Feature Bundling (EFB), and it grows trees leaf-wise (best-first) rather than level by level, always splitting the leaf with the largest loss reduction. The objective is to minimise a regularised loss of the same form as XGBoost,

$$\mathcal{L}^{(t)} = \sum_{i=1}^{n} \ell\Big(y_i, \hat{y}_i^{(t-1)} + f_t(x_i)\Big) + \Omega(f_t), \tag{S50}$$

where $\ell(\cdot,\cdot)$ is a convex per-sample loss (typically multi-class log-loss for classification) and $\Omega(f_t) = \gamma T + \frac{1}{2}\lambda\|w\|^2$ penalises the number of leaves $T$ and the magnitude of the leaf weights $w$ in tree $f_t$. GOSS keeps the instances with large gradients and randomly samples those with small ones, which keeps the estimate of information gain balanced. In the `lightgbm_classification` wrapper, the `verbosity=-1` parameter suppresses training output. Because LightGBM copes with data that are both large and high-dimensional, and returns usable probability estimates, it sits comfortably within an automated benchmarking pipeline. It is implemented through the function `LGBMClassifier` from `lightgbm`.

**Benchmark and Pipeline (`benchmark_models_comprehensive, supervised_analysis`)** The `benchmark_models_comprehensive` function evaluates all the classifiers above in one framework, with three validation strategies: stratified k-fold cross-validation, a single holdout split, and the .632+ bootstrap estimator. Each model's predictions are scored against ground-truth annotations in both the original high-dimensional gene space and a reduced PCA space (default: 30 components). The dimensionality reduction step is wrapped in a `Pipeline`, so PCA is fitted only on the training folds and no information leaks from the held-out data during bootstrap or CV. Performance is assessed mainly by Accuracy and the macro-averaged F1-score:

$$\mathrm{F1}_{\mathrm{macro}} = \frac{1}{K}\sum_{k=1}^{K} \frac{2 \cdot \mathrm{Precision}_k \cdot \mathrm{Recall}_k}{\mathrm{Precision}_k + \mathrm{Recall}_k} \tag{S51}$$

where $K$ is the number of classes; per-class terms with $\mathrm{Precision}_k + \mathrm{Recall}_k = 0$ contribute zero to the sum by convention. For cross-validation, the framework reports the mean metric across folds together with the standard error of the mean, $\mathrm{SE} = \sigma/\sqrt{k}$, where $\sigma$ is the sample standard deviation of the fold-wise metric and $k$ is the number of folds. When the `bootstrap_ci` flag is enabled, BCa (Bias-Corrected and accelerated) confidence intervals give non-parametric bounds for the estimates. Results come back as a `DataFrame` with columns for `space`, `model`, and the `mean`, `std` and `se` of each metric, together with a grouped bar chart comparing performance across feature spaces. A separate utility, `compare_classifiers`, tests any two models against each other with a paired bootstrap test and returns p-values for the difference in classification performance.

**Feature Importance (`shap_analysis`)** Model-agnostic interpretability comes from the `shap_analysis` module, which measures how much each feature contributes to a model's predictions. Its main method is SHAP (SHapley Additive exPlanations), which comes from cooperative game theory. The SHAP value for a feature $i$ is defined as the average marginal contribution across all possible feature subsets $S \subseteq N \setminus \{i\}$:

$$\phi_i = \sum_{S \subseteq N\setminus\{i\}} \frac{|S|!(|N|-|S|-1)!}{|N|!}[v(S \cup \{i\}) - v(S)] \tag{S52}$$

where $v(S)$ is the model's prediction using only the features in subset $S$. Global feature importance is the mean absolute SHAP value across all samples and, in multi-class problems, across all classes. If the `shap` library is not installed, the module falls back automatically to Permutation Feature Importance, which scores a feature by how much the model's performance drops when that feature's values are randomly shuffled. Either way, the output is a `pd.DataFrame` of features sorted by importance score. Reading down it gives the candidate markers behind the classification boundaries.

**Supplementary table 13:** High level functions for supervised benchmark and feature importance implemented in scINTILLA.

| **Method** | **Inputs** | **Outputs** |
|---|---|---|
| Comprehensive Benchmark<br>`benchmark_models_-comprehensive` | `data`: AnnData or DataFrame<br>`target_col`: class labels in `obs`<br>(`estimator`: `"cv"`, `"holdout"`, `"bootstrap_632plus"`)<br>(`cv_folds`: default 5)<br>(`use_pca`: default `True`)<br>(`n_pca_comps`: default 30)<br>(`bootstrap_ci`: BCa intervals) | `results_df`: mean metrics, SE, and optional CIs<br>`fig`: grouped bar chart (Accuracy) |
| Classifier Comparison<br>`compare_classifiers` | `y_true`: ground-truth labels<br>`predictions`: dict of `{model_name: y_-pred}`<br>(`metric_fn`: default accuracy)<br>(`B`: bootstrap replicates) | `results_df`: pairwise differences, CIs, and $p$-values |
| Feature Importance<br>`shap_analysis` | `model`: fitted estimator<br>`X_train`: background data<br>`X_test`: evaluation data<br>(`y_test`: required for fallback)<br>(`feature_names`) | `shap_values`: raw SHAP array (or `None`)<br>`feature_importance_df`: ranked feature importance |

### S7 Differential Expression

The `differential_expression` sub-package measures expression shifts between groups with several statistical models: classical rank- and mean-based tests, pseudobulk linear models and non-parametric permutation procedures. All entry points share a common interface that returns a tidy `DataFrame` of per-gene statistics together with multiple-testing-adjusted $p$-values.

**Wilcoxon rank-sum test (`wilcoxon_de`)** The Wilcoxon (Mann–Whitney $U$) test is the standard non-parametric test for a shift in the distribution of a single gene between two groups. Let $\mathrm{rk}_{g,i}$ denote the rank of cell $i$ for gene $g$ across the pooled samples and $\mathcal{C}_a, \mathcal{C}_b$ the two groups; the rank-sum statistic is

$$U_g = \sum_{i \in \mathcal{C}_a} \mathrm{rk}_{g,i} - \frac{|\mathcal{C}_a|(|\mathcal{C}_a| + 1)}{2}. \tag{S53}$$

Under the null hypothesis of identical distributions, $U_g$ is approximately Gaussian with mean $|\mathcal{C}_a||\mathcal{C}_b|/2$ and a variance that incorporates a tie-correction term. scINTILLA reports the standardised $z$-score, the two-sided $p$-value, and the log-fold-change of group means.

**Two-sample $t$-test (`ttest_de`)** For datasets in which a Gaussian-like model on transformed expression is appropriate, scINTILLA also exposes Welch's two-sample $t$-test,

$$t_g = \frac{\bar{x}_{g,a} - \bar{x}_{g,b}}{\sqrt{s_{g,a}^2/n_a + s_{g,b}^2/n_b}}, \tag{S54}$$

with Satterthwaite degrees of freedom. Its inclusion lets users sanity-check Wilcoxon hits in a parametric setting and quantify directional effect sizes.

**Pseudobulk DE (`pseudobulk_de`)** Per-cell tests treat each cell as an independent replicate, which inflates type-I error when biological replication occurs at the sample level. The pseudobulk strategy collapses cells of the same group within each donor or replicate into a single summed count vector and then applies a sample-level test (Welch's $t$ on log-CPM by default). For donor $d$ in group $a$ and gene $g$,

$$y_{g,d,a} = \log\left( \frac{\sum_{i \in \mathcal{C}_a \cap \mathcal{D}_d} x_{g,i}}{\sum_j \sum_{i \in \mathcal{C}_a \cap \mathcal{D}_d} x_{j,i}} \cdot 10^6 + 1 \right), \tag{S55}$$

and the test is performed on $\{y_{g,d,a}\}_d$ versus $\{y_{g,d,b}\}_d$. This honours the experimental unit of replication and yields properly calibrated $p$-values [36].

**Permutation test (`permutation_de`)** For genes whose marginal distribution departs strongly from Gaussianity, scINTILLA computes an exact non-parametric $p$-value by permuting the group labels $B$ times and recomputing the chosen statistic $T$. The two-sided estimator

$$\hat{p}_g = \frac{1 + \#\{b : |T_g^{(b)}| \geq |T_g^{\mathrm{obs}}|\}}{B + 1} \tag{S56}$$

includes the observed statistic in both numerator and denominator to guarantee a valid (non-zero) $p$-value under any $B$ [32].

**Rank-genes-groups wrapper (`rank_genes_groups`)** `rank_genes_groups` is a thin convenience layer around `scanpy.tl.rank_genes_groups` that runs a one-vs-rest test for every level of a categorical covariate and returns the top-$n$ genes per group as a tidy table. The default test is Wilcoxon with Benjamini–Hochberg correction; the user may switch to the $t$-test or to a logistic-regression score.

**Multiple-testing correction and benchmark** All raw $p$-values are passed through `correct_pvalues`, which applies Benjamini–Hochberg control of the false discovery rate [2]; Bonferroni and Holm corrections are also exposed for users who require strict family-wise error rate control. The companion routine `benchmark_de_methods` runs the Wilcoxon, $t$-test and pseudobulk methods on the same two groups and reports, for each, the number of genes tested and the number significant at an adjusted $p < 0.05$, so that the three can be compared on the same data.

### S8 Cell-Type Annotation

The `annotation` sub-package assigns cell-type labels under two complementary regimes: marker-based scoring, which uses curated gene panels, and label transfer, which propagates labels from a reference dataset. Both regimes are supported by a marker-discovery utility and an over-representation test for downstream functional interpretation.

**Marker-based annotation (`annotate_by_markers`)** Given a dictionary $\mathcal{M} = \{c \mapsto \mathcal{G}_c\}$ that associates each candidate cell type $c$ with a marker-gene set $\mathcal{G}_c$, each cell $i$ is scored using the Scanpy convention,

$$s_{c,i} \;=\; \frac{1}{|\mathcal{G}_c|} \sum_{g \in \mathcal{G}_c} x_{g,i} \;-\; \frac{1}{|\mathcal{B}_c|} \sum_{g \in \mathcal{B}_c} x_{g,i}, \tag{S57}$$

where $\mathcal{B}_c$ is a control set of similarly expressed genes sampled from the same expression bins; it centres the score, so that random gene sets score around zero. Cells are then assigned the label of the highest-scoring panel, with a confidence margin reported as the gap between the top two scores.

**Marker discovery (`find_marker_genes`)** Marker discovery is implemented as a one-vs-rest Wilcoxon ranking with Benjamini–Hochberg-adjusted $p$-values, restricted to genes that exceed user-defined fold-change and minimum-expression thresholds. The output is a per-cluster ranked list suitable both for downstream over-representation testing and for hand-curation of new marker panels.

**Over-representation analysis (`ora_test`)** For a candidate gene set $\mathcal{S}$, a reference gene-set library $\mathcal{R}$ (e.g. a pathway database), and a chosen background of $N$ genes, the over-representation $p$-value for each library term $T \in \mathcal{R}$ follows the hypergeometric distribution,

$$P(X \geq k) \;=\; \sum_{x=k}^{\min(|T|,|\mathcal{S}|)} \frac{\binom{|T|}{x}\binom{N-|T|}{|\mathcal{S}|-x}}{\binom{N}{|\mathcal{S}|}}, \tag{S58}$$

where $k$ is the observed overlap between $\mathcal{S}$ and $T$. Raw $p$-values are corrected with Benjamini–Hochberg.

**Label transfer (`transfer_labels`)** Labels are propagated from a reference dataset to a query dataset by majority vote among the $k$ nearest reference cells in a shared embedding (PCA by default). For a query cell $q$ with reference neighbours $\mathcal{N}_k(q)$ carrying labels $\{c(r)\}_{r \in \mathcal{N}_k(q)}$,

$$\hat{c}(q) \;=\; \arg\max_{c} \sum_{r \in \mathcal{N}_k(q)} \mathbf{1}[c(r) = c], \tag{S59}$$

with an associated certainty score equal to the modal vote share. The routine returns both the predicted labels and the per-cell certainty, so low-confidence assignments can be filtered out by a threshold.

### S9 Batch Correction

In multi-batch single-cell studies, technical batch effects have to be removed before cells from different batches can be compared. scINTILLA exposes the empirical-Bayes ComBat correction together with a set of batch-mixing metrics to evaluate it.

**ComBat (`combat_correct`)** ComBat [23] models the log-transformed expression of gene $g$ in cell $i$ from batch $b$ as

$$Y_{b,g,i} \;=\; \alpha_g + X_i^\top \beta_g + \gamma_{b,g} + \delta_{b,g}\, \varepsilon_{b,g,i}, \tag{S60}$$

with $\alpha_g$ the gene mean, $X_i$ the vector of biological covariates of cell $i$ that are to be preserved, $\gamma_{b,g}$ the additive batch shift, $\delta_{b,g}$ the multiplicative batch variance, and $\varepsilon_{b,g,i} \sim \mathcal{N}(0, \sigma_g^2)$. An empirical-Bayes step shrinks the batch parameters towards a common prior, which stabilises the estimates when individual batches contain few cells. The adjusted expression is

$$Y^*_{b,g,i} \;=\; \frac{Y_{b,g,i} - \hat{\alpha}_g - X_i^\top \hat{\beta}_g - \hat{\gamma}^*_{b,g}}{\hat{\delta}^*_{b,g}} + \hat{\alpha}_g + X_i^\top \hat{\beta}_g, \tag{S61}$$

where starred parameters are the empirical-Bayes posterior estimates. The implementation delegates to `scanpy.pp.combat` and preserves any user-supplied biological covariates.

**Evaluation metrics** Batch correction is evaluated jointly on *batch mixing* (the corrected embedding should not retain batch identity) and *biology preservation* (it should still separate biological cell types). scINTILLA implements the following metrics.

*Local Inverse Simpson's Index (LISI [25]).* For each cell $i$, with neighbourhood weights $w_{ij}$ from a Gaussian kernel on the kNN graph and discrete labels $b(j)$ (batch for iLISI, cell type for cLISI),

$$\begin{aligned} \mathrm{LISI}_i &= \left(\sum_c \hat{p}_c(i)^2\right)^{-1}, \\ \hat{p}_c(i) &= \frac{\sum_j w_{ij}\,\mathbf{1}[b(j)=c]}{\sum_j w_{ij}}. \end{aligned} \tag{S62}$$

$\mathrm{LISI}_i$ approaches the number of categories represented in $i$'s neighbourhood; iLISI close to the number of batches indicates good mixing, whereas cLISI close to unity indicates preserved cell-type purity.

*kBET [7].* kBET tests whether the local batch composition of each cell's neighbourhood matches the global composition via a $\chi^2$ test on counts in a $K$-nearest-neighbour window, summarised across cells as the rejection rate at significance level $\alpha$.

*Batch-ASW.* The batch-aware silhouette width recasts the silhouette coefficient with batch as the grouping variable, so that values near zero indicate well-mixed batches.

**Benchmark (`benchmark_batch_correction`)** The wrapper applies each correction method to a common input and computes the metrics above on the corrected and uncorrected embeddings. It returns the comparison as a table, from which a method can be chosen.

### S10 Robust Statistics

A separate `statistical_tests` sub-package consolidates the inferential routines that underpin the benchmark modules. The accent throughout is on calibrated, assumption-light estimators with explicit uncertainty quantification.

**Per-gene ANOVA and Kruskal–Wallis** One-way ANOVA and its rank-based analogue, the Kruskal–Wallis test, screen for genes whose expression differs across more than two groups. Their statistics, applied per gene, are

$$F = \frac{\mathrm{MS}_{\text{between}}}{\mathrm{MS}_{\text{within}}}, \qquad H = \frac{12}{n(n+1)}\sum_{g=1}^{G}\frac{R_g^2}{n_g} - 3(n+1), \tag{S63}$$

where $R_g$ is the sum of ranks in group $g$ and $n_g$ its size.

**Box's $M$ test** Box's $M$ tests whether covariance matrices are equal across groups. LDA assumes they are; a rejection points towards QDA. With pooled covariance $S_p$ and per-group estimates $S_g$,

$$M = (n-G)\log|S_p| - \sum_{g=1}^{G}(n_g-1)\log|S_g|, \tag{S64}$$

distributed asymptotically as a $\chi^2$ on $\frac{1}{2}(G-1)p(p+1)$ degrees of freedom. The determinants exist only when every group has more cells than features ($n_g > p$), so the test is computed in a low-dimensional space such as the PCA and is skipped for smaller groups.

**Multiple-testing correction (`correct_pvalues`)** scINTILLA exposes Benjamini–Hochberg (FDR), Bonferroni, and Holm corrections through a single dispatcher. The Benjamini–Hochberg threshold is

$$\hat{k} = \max\left\{k : p_{(k)} \le \tfrac{k}{m}\,\alpha\right\}, \tag{S65}$$

with $p_{(1)} \le \cdots \le p_{(m)}$ the sorted raw $p$-values and $\alpha$ the target FDR level.

**.632+ bootstrap (`dot632plus_bootstrap`)** For nearly unbiased generalisation error on benchmark datasets, scINTILLA implements the Efron–Tibshirani .632+ bootstrap [13],

$$\begin{aligned} \widehat{\mathrm{Err}}^{(.632+)} &= (1-\hat{w})\,\widehat{\mathrm{err}}_{\mathrm{train}} + \hat{w}\,\widehat{\mathrm{Err}}^{(1)}, \\ \hat{w} &= \frac{0.632}{1-0.368\,\hat{R}}, \end{aligned} \tag{S66}$$

where $\widehat{\mathrm{err}}_{\mathrm{train}}$ is the apparent training error, $\widehat{\mathrm{Err}}^{(1)}$ is the leave-one-out bootstrap error, and $\hat{R} = (\widehat{\mathrm{Err}}^{(1)} - \widehat{\mathrm{err}}_{\mathrm{train}})/(\hat{\gamma} - \widehat{\mathrm{err}}_{\mathrm{train}})$ is the relative overfitting rate with $\hat{\gamma}$ the no-information error rate. The estimator interpolates between optimistic and pessimistic bootstrap variants and corrects for the .632 estimator's residual bias under heavy overfitting.

**Effect sizes** Beyond $p$-values, scINTILLA reports calibrated effect sizes: Cohen's $d$, Hedges' $g$ with small-sample correction, and Cliff's $\delta$. For two independent samples of sizes $n_1, n_2$,

$$\begin{aligned} d &= \frac{\bar{x}_1 - \bar{x}_2}{s_p}, \quad g = J(\nu)\,d, \\ \delta &= \frac{1}{n_1 n_2}\sum_{i,j}\left[\mathbf{1}(x_{1i} > x_{2j}) - \mathbf{1}(x_{1i} < x_{2j})\right], \end{aligned} \tag{S67}$$

where $s_p$ is the pooled standard deviation, $\nu = n_1 + n_2 - 2$, and $J(\nu) = 1 - 3/(4\nu - 1)$ is Hedges' bias correction.

**Data-adaptive thresholds** For automated PCA truncation, scINTILLA implements the Gavish–Donoho optimal hard threshold [17],

$$\lambda^* = \omega(\beta)\,\sigma\,\sqrt{n}, \qquad \beta = m/n \leq 1, \tag{S68}$$

where $m \leq n$ are the two dimensions of the data matrix, $\sigma$ is the estimated noise level and $\omega(\beta)$ is the asymptotically optimal scaling factor; together with the Marchenko–Pastur edge

$$\lambda_+ = \sigma\,\sqrt{n}\left(1 + \sqrt{\beta}\right), \tag{S69}$$

written here on the same singular-value scale, above which singular values are statistically distinguishable from noise. These thresholds drive automatic component-count selection inside the preprocessing module.
