## Supplementary Figures and Tables for "scINTILLA: Single-Cell Integrated Inference, Labelling, and Landscape Analysis for Assessing the Quality and Hidden Diversity of Cell Annotations"

Kanannejad *et al.*

**Supplementary table 1:** Sensitivity of the benchmark to batch correction (exploratory). Balanced AUROC on the label-free PCA (main analysis) and on the same PCA corrected with Harmony (donor as batch), for the eleven methods that score every label; same cells and perturbations. SCCAF does not use the embedding.

| Method | PCA | Rank | Harmony | Rank |
|---|---|---|---|---|
| scINTILLA-CS + Silhouette | 0.897 | 4 | 0.886 | 1 |
| scINTILLA-F + Silhouette | 0.908 | 1 | 0.882 | 2 |
| scINTILLA learned fusion (additive) | 0.904 | 2 | 0.874 | 3 |
| scINTILLA-F | 0.890 | 6 | 0.872 | 4 |
| scINTILLA learned fusion (interaction) | 0.903 | 3 | 0.868 | 5 |
| SCCAF + Silhouette | 0.869 | 8 | 0.858 | 6 |
| Silhouette | 0.896 | 5 | 0.857 | 7 |
| scINTILLA-CS | 0.851 | 9 | 0.846 | 8 |
| kNN purity + Silhouette | 0.882 | 7 | 0.824 | 9 |
| SCCAF | 0.770 | 11 | 0.770 | 10 |
| kNN purity | 0.815 | 10 | 0.747 | 11 |

**Supplementary table 2:** Effect of Harmony on donor structure in the benchmark embeddings. Share of each cell's 50 nearest neighbours that come from the same donor, before (PCA) and after Harmony correction (donor as batch), in the cell pools used for the benchmark.

| Atlas | Cells | Donors | PCA | Harmony |
|---|---|---|---|---|
| Brain | 23,241 | 4 | 0.518 | 0.393 |
| Eye | 101,921 | 104 | 0.107 | 0.056 |
| HEOCA | 36,702 | 200 | 0.608 | 0.222 |
| HNOCA | 51,203 | 54 | 0.704 | 0.318 |
| Lung | 58,583 | 482 | 0.227 | 0.077 |

**Supplementary table 3:** Sensitivity of the real-label scores to the embedding (exploratory). The original annotations were rescored with the same cells, labels and classifiers, replacing the label-free PCA with the Harmony-corrected PCA (all atlases) or with the atlas-provided scVI space (eye). Same-donor share: fraction of each cell's 50 nearest neighbours from the same donor. Spearman: rank correlation of the per-label scores with the PCA-based scores. Review-list overlap: shared labels among the lowest 10% of labels under scINTILLA-F + Silhouette. Case-study labels: rank under scINTILLA-F + Silhouette with the alternative embedding versus PCA.

| Atlas | Embedding | Same-donor share | Spearman with PCA | | | Review-list | Case-study labels |
|---|---|---|---|---|---|---|---|
| | | (PCA → alt.) | F + Silhouette | scINTILLA-F | Silhouette | overlap | (alt. vs PCA) |
| Brain | Harmony | 0.49 → 0.37 | 0.96 | 0.95 | 0.99 | 2/3 | neuron 1 vs 1 |
| Eye | Harmony | 0.09 → 0.05 | 0.92 | 0.93 | 0.92 | 10/13 | – |
| Eye | scVI | 0.09 → 0.13 | 0.83 | 0.88 | 0.74 | 8/13 | – |
| HEOCA | Harmony | 0.56 → 0.19 | 0.69 | 0.69 | 0.68 | 3/4 | stem cell 2 vs 1 |
| HNOCA | Harmony | 0.69 → 0.28 | 0.85 | 0.86 | 0.83 | 3/6 | unknown 1 vs 1; neuron 5 vs 3 |
| Lung | Harmony | 0.20 → 0.06 | 0.67 | 0.71 | 0.64 | 4/7 | Unknown 1 vs 1; Monocyte-derived Mph 3 vs 3 |

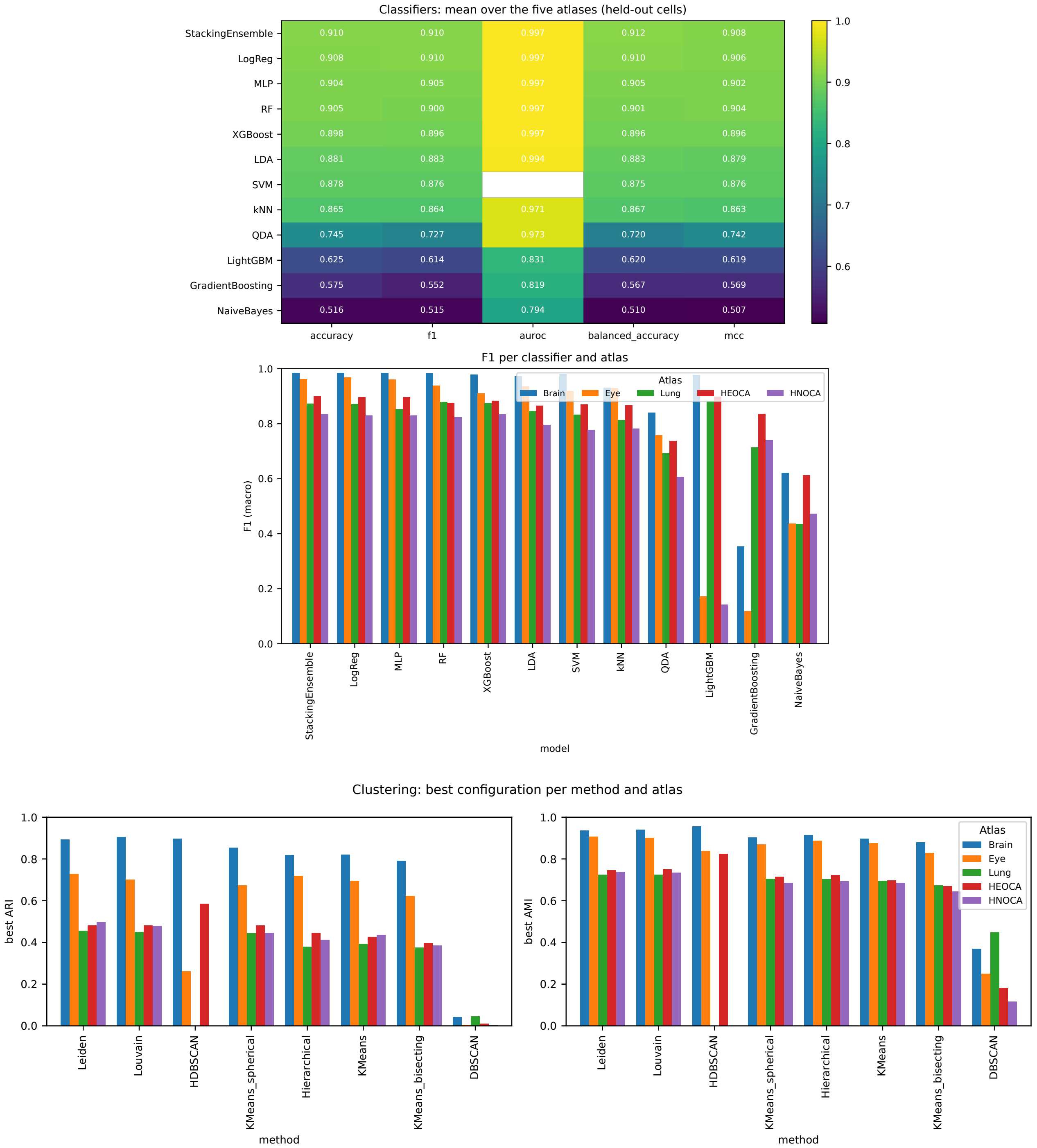


**Supplementary figure 1:** Classifier and clustering benchmarks across the five atlases (configuration used in this study: twelve classifiers trained on 2,000 highly variable genes; clustering on a 30-component PCA). Top: held-out performance of each classifier averaged over the atlases (accuracy, F1, AUROC, balanced accuracy and MCC; AUROC is not available for the SVM) and F1 per classifier and atlas. Bottom: best ARI (left) and AMI (right) for each clustering method and atlas over its parameter configurations; HDBSCAN returned no usable partition for the lung and HNOCA atlases.

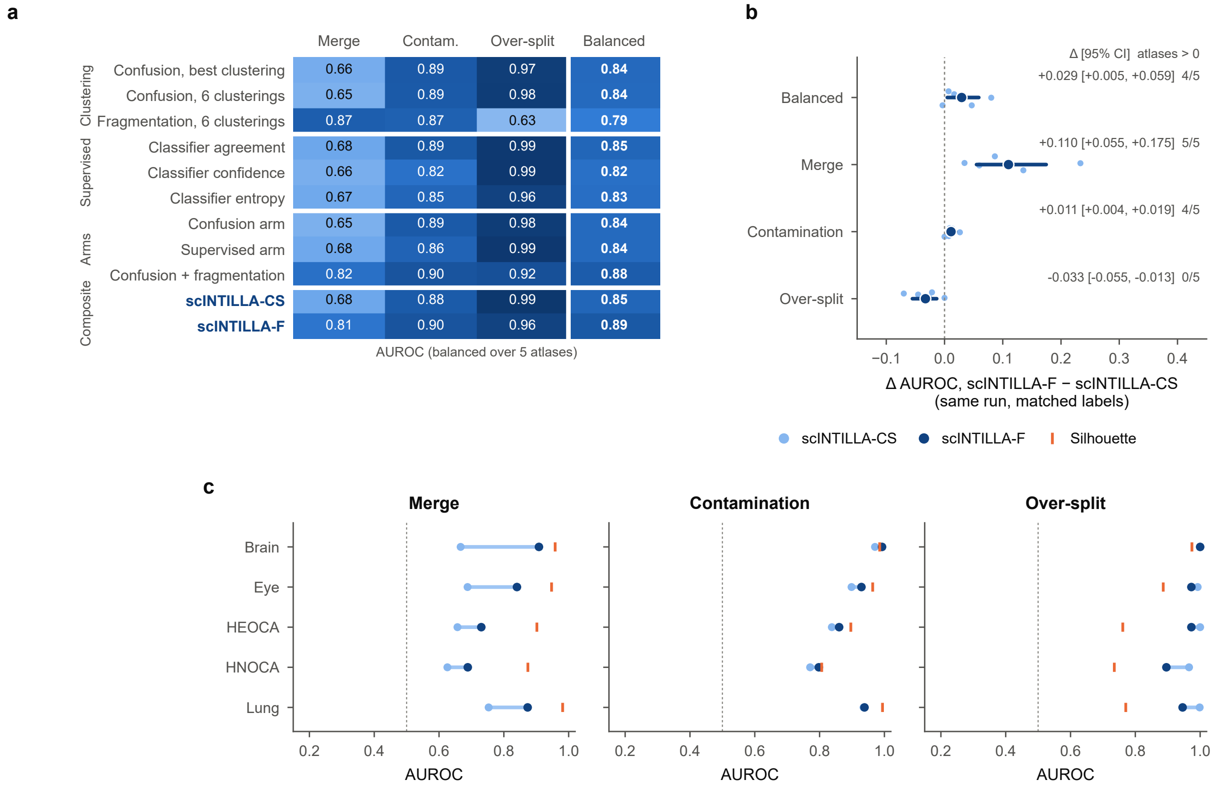


**Supplementary figure 2:** Contribution of each signal to scINTILLA. **(a)** AUROC by error type and balanced over the five atlases for the individual components (clustering confusion, fragmentation, classifier agreement, confidence and entropy), the two arms and the composites. **(b)** Paired difference between scINTILLA-F and scINTILLA-CS (without fragmentation) on matched labels of the same run, with 95% atlas-bootstrap intervals and the number of atlases with a positive difference. **(c)** AUROC per atlas and error type for scINTILLA-CS, scINTILLA-F and the Silhouette baseline.

**Supplementary table 4:** Re-scoring of the re-annotated compartments. The original and the refined labels were scored on identical cells (stratified by refined label, at most 1,000 cells per label; 9,000 lung and 13,000 HEOCA cells) with the benchmarked configuration, on the label-free PCA and on the same PCA corrected with Harmony (donor as batch). The HEOCA stem-cell group holds six original labels: stem cell, epithelial cell, goblet cell, enterocyte, colon epithelial cell and M cell of gut. Values are unscaled, cell-weighted means, original → refined; confusion and fragmentation refer to the best clustering. Classifier disagreement is the fraction of classifiers that predict a class other than the assigned label, and consensus mismatch is the fraction of cells whose majority prediction differs from it; neither uses the embedding. Composite scores are scaled within each labelling and are therefore not compared.

| | Lung myeloid (7 → 9 labels) | | HEOCA stem-cell group (6 → 13 labels) | |
|---|---|---|---|---|
| Metric | PCA | Harmony | PCA | Harmony |
| Classifier disagreement ↓ | 0.157 → 0.107 | 0.157 → 0.107 | 0.076 → 0.073 | 0.076 → 0.073 |
| Consensus mismatch ↓ | 0.045 → 0.038 | 0.045 → 0.038 | 0.010 → 0.025 | 0.010 → 0.025 |
| Clustering confusion ↓ | 0.295 → 0.347 | 0.396 → 0.572 | 0.095 → 0.204 | 0.270 → 0.382 |
| Label fragmentation ↓ | 0.437 → 0.751 | 0.558 → 0.735 | 0.645 → 0.527 | 0.604 → 0.556 |
| Silhouette ↑ | −0.042 → 0.000 | −0.056 → −0.029 | 0.021 → 0.157 | −0.044 → 0.042 |

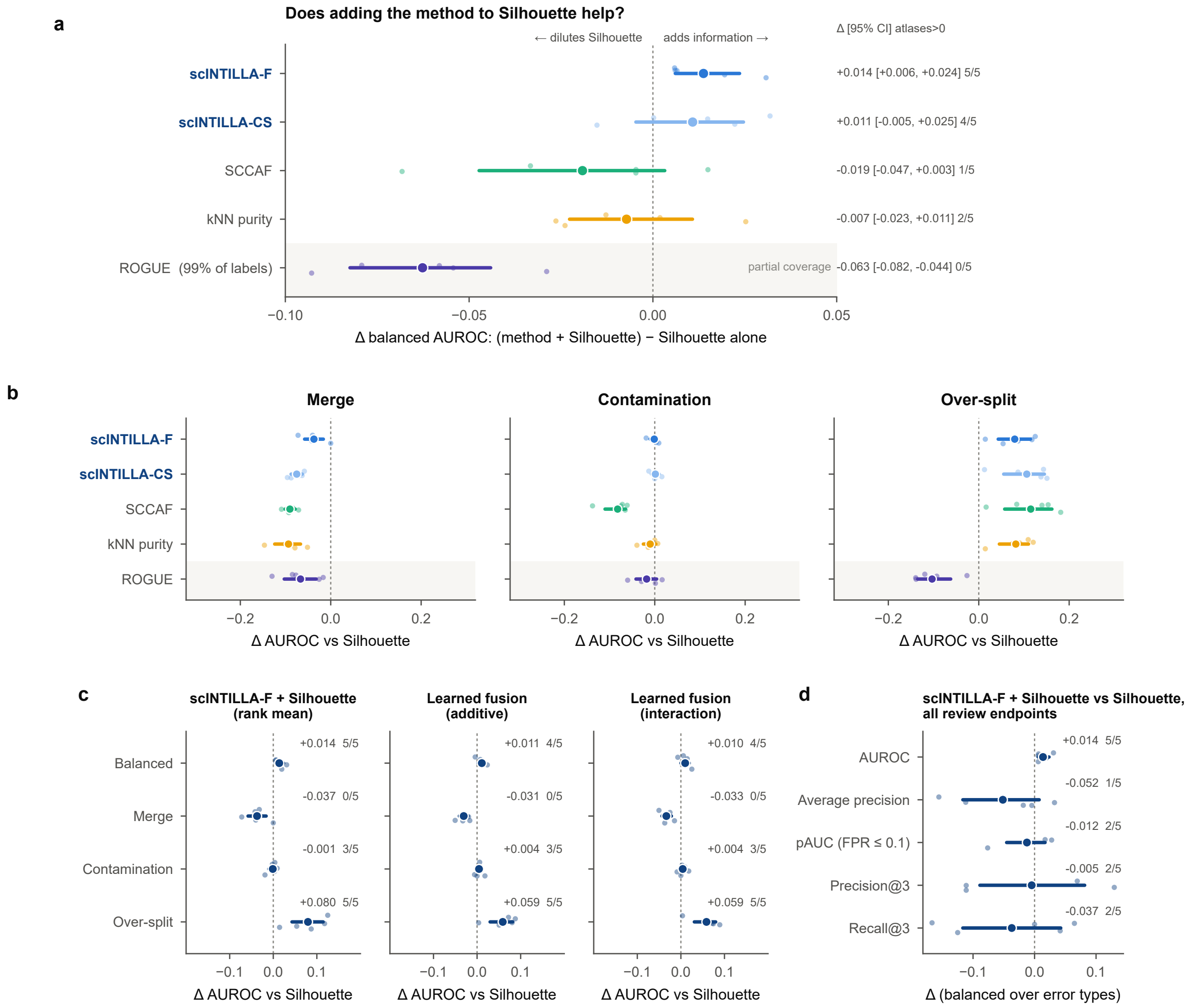


**Supplementary figure 3:** Does a method add information to the Silhouette baseline? **(a)** Change in balanced AUROC when each method is rank-fused with Silhouette, relative to the Silhouette baseline alone, with 95% atlas-bootstrap intervals and the number of atlases with a positive change; ROGUE scores 99% of labels. **(b)** The same change by error type; points are atlases. **(c)** Change in AUROC relative to the Silhouette baseline for the scINTILLA-F + Silhouette rank fusion and the two learned fusions, by error type. **(d)** scINTILLA-F + Silhouette against the Silhouette baseline on all review endpoints (AUROC, average precision, partial AUROC at FPR $\leq 0.1$, precision@3 and recall@3).

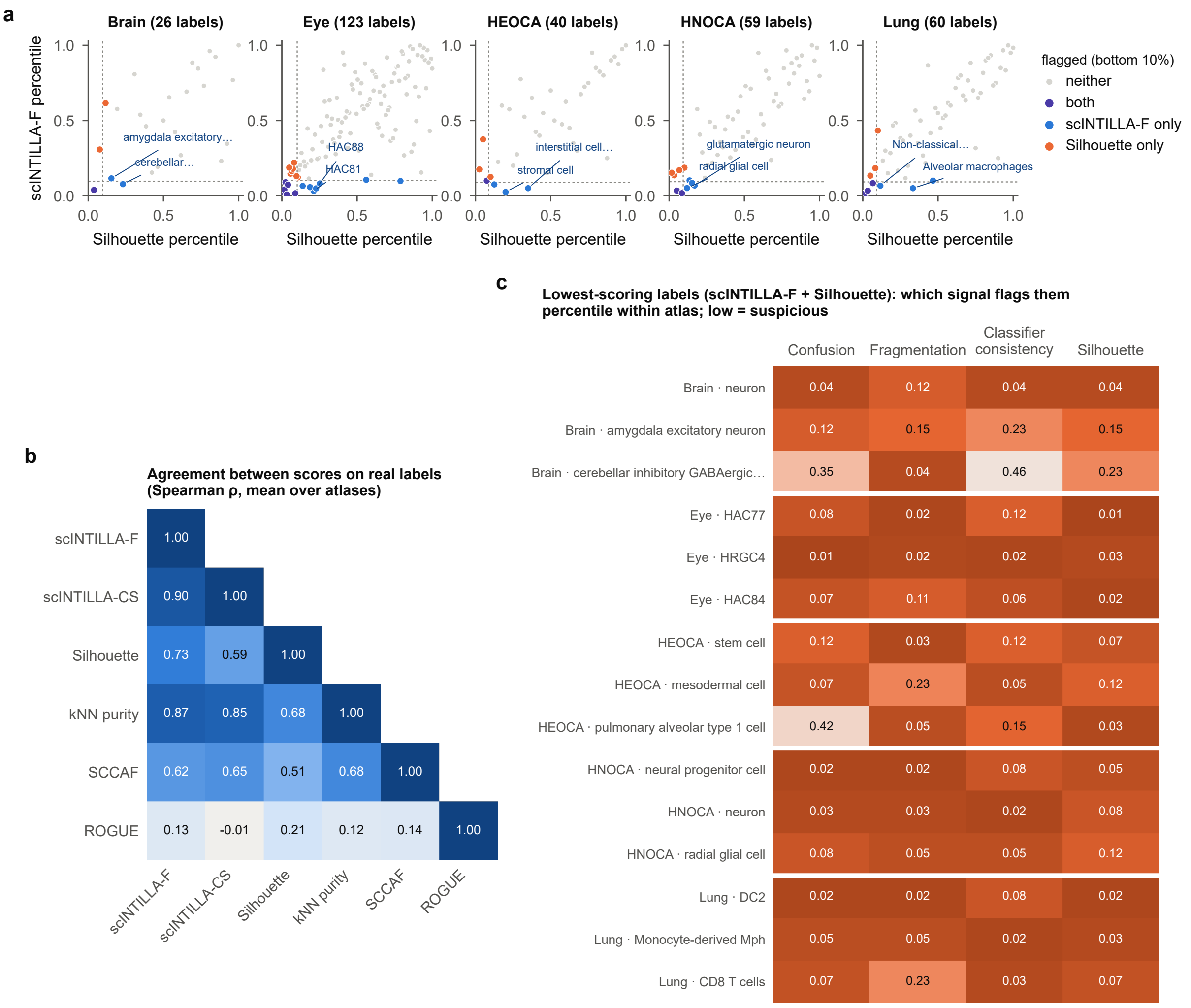


**Supplementary figure 4:** scINTILLA on the original (unperturbed) labels of the five atlases, scored with the benchmarked configuration (at most 500 cells per label); "unknown" labels are not shown, so HNOCA has 59 and the lung 60 labels here. **(a)** Percentile of each label under scINTILLA-F against its Silhouette percentile; labels in the bottom 10% of either score are coloured by which score flags them. **(b)** Spearman correlation between scores on the real labels, averaged over the atlases. **(c)** The three lowest-ranked labels per atlas under scINTILLA-F + Silhouette, with the percentile of each signal (low = suspicious) to show which signal flags each label.

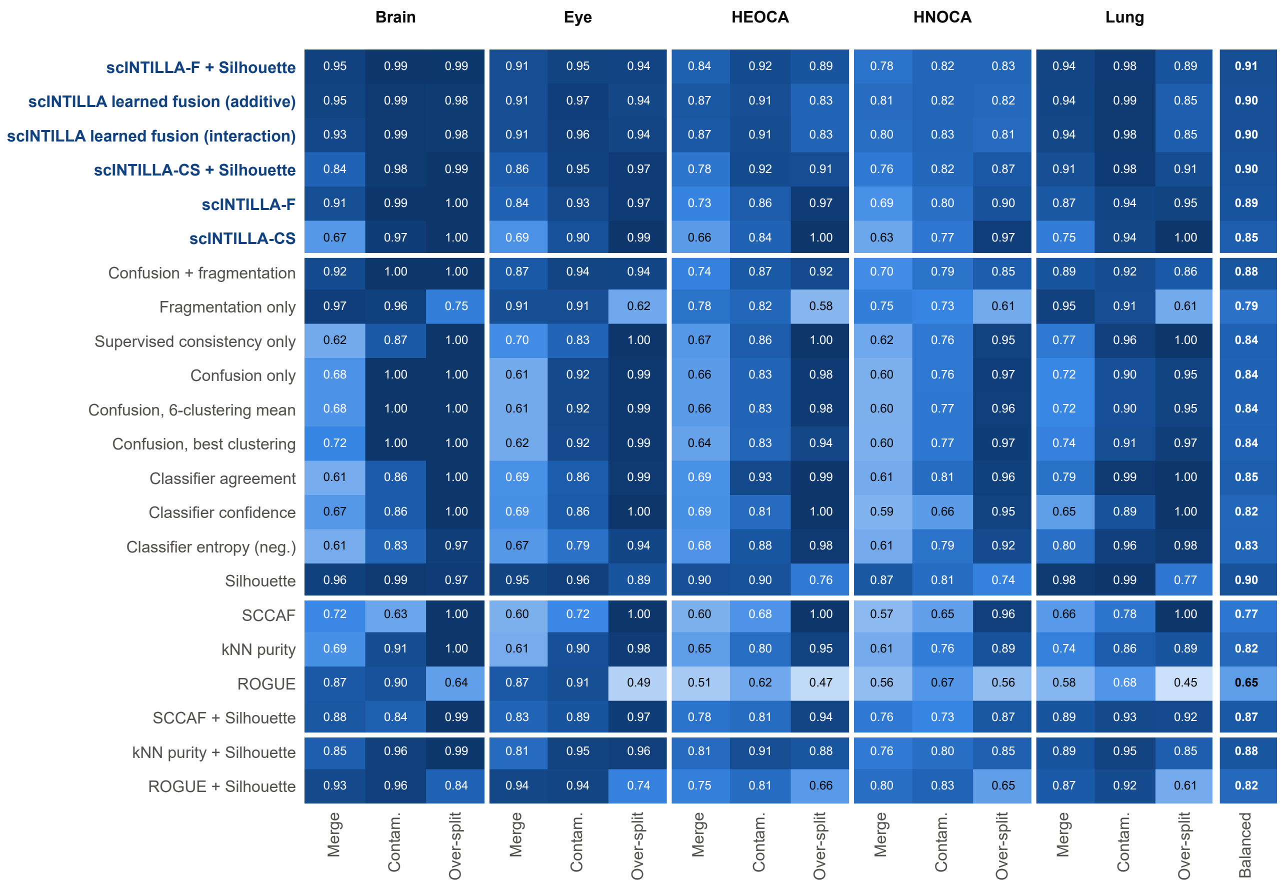


**Supplementary figure 5:** AUROC for every method and component per atlas and error type (merge, contamination and over-split; median of ten replicates), with the balanced value in the last column.

**Supplementary table 5:** Donor-level tests of the enrichments named in the text. A unit is one donor within one condition (lung) or tissue (HEOCA). Donors: units in the group, with the number that carry the state in brackets. Share: median share of a unit's cells in the state, inside against outside the group. MW: Benjamini–Hochberg FDR of the two-sided Mann–Whitney U test. QB: odds ratio of the quasi-binomial model, with its FDR in brackets. A dash marks a pair with fewer than three donors carrying the state, which was not tested.

| State | Group | Donors | Share (in vs out) | MW FDR; QB OR (FDR) | Verdict |
|---|---|---|---|---|---|
| *Lung* | | | | | |
| Cycling Mph | IPF | 52 (50) | 0.035 vs 0.003 | $3 \times 10^{-13}$; 3.7 ($1 \times 10^{-25}$) | supported |
| TREM2-high MDM | IPF | 52 (52) | 0.33 vs 0.12 | $1 \times 10^{-11}$; 2.0 ($8 \times 10^{-12}$) | supported |
| TREM2-high MDM | Pneumonia, non-COVID | 12 (12) | 0.38 vs 0.14 | 0.007; 2.0 (0.016) | supported |
| Classical monocytes (STR) | COVID-19 (merged) | 58 (43) | 0.002 vs 0 | $2 \times 10^{-10}$; 2.8 ($3 \times 10^{-4}$) | supported |
| Classical monocytes (STR) | COVID-19 severe | 6 (6) | 0.007 vs 0 | $7 \times 10^{-4}$; 9.5 ($4 \times 10^{-16}$) | one donor holds 89% |
| Cycling Mph | Hypersensitivity pneumonitis | 3 (3) | 0.046 vs 0.004 | 0.07; 3.4 (0.014) | one test only |
| TREM2-high MDM | Squamous cell carcinoma | 3 (3) | 0.37 vs 0.15 | 0.09; 1.9 (0.14) | not supported |
| Cycling Mph | NSIP | 3 (2) | – | – | too few donors |
| TREM2-high MDM; Cycling Mph | Myositis-associated ILD | 1 (1) | – | – | too few donors |
| *HEOCA* | | | | | |
| SOX9/PROM1 progenitors | Biliary system | 5 (5) | 0.41 vs 0.13 | 0.027; 4.0 (0.055) | one test only |
| AFP hepatoblast-like | Liver | 19 (13) | 0.026 vs 0.0002 | 0.026; 3.1 (0.84) | one test only |
| AFP hepatoblast-like | Intestine | 19 (17) | 0.025 vs 0 | 0.003; 3.6 (0.47) | one test only |
| Neural-like (AUTS2) | Colon | 12 (5) | 0 vs 0 | 0.99; 297 ($8 \times 10^{-7}$) | one test only |
| NKX2-1+ (single sample) | Alveolar sac | 6 (1) | – | – | one sample |
| KRT6B basal epithelial | Prostate gland | 1 (1) | – | – | one donor |
| SOX11 mesenchymal progenitors | Pancreas | 2 (2) | – | – | too few donors |

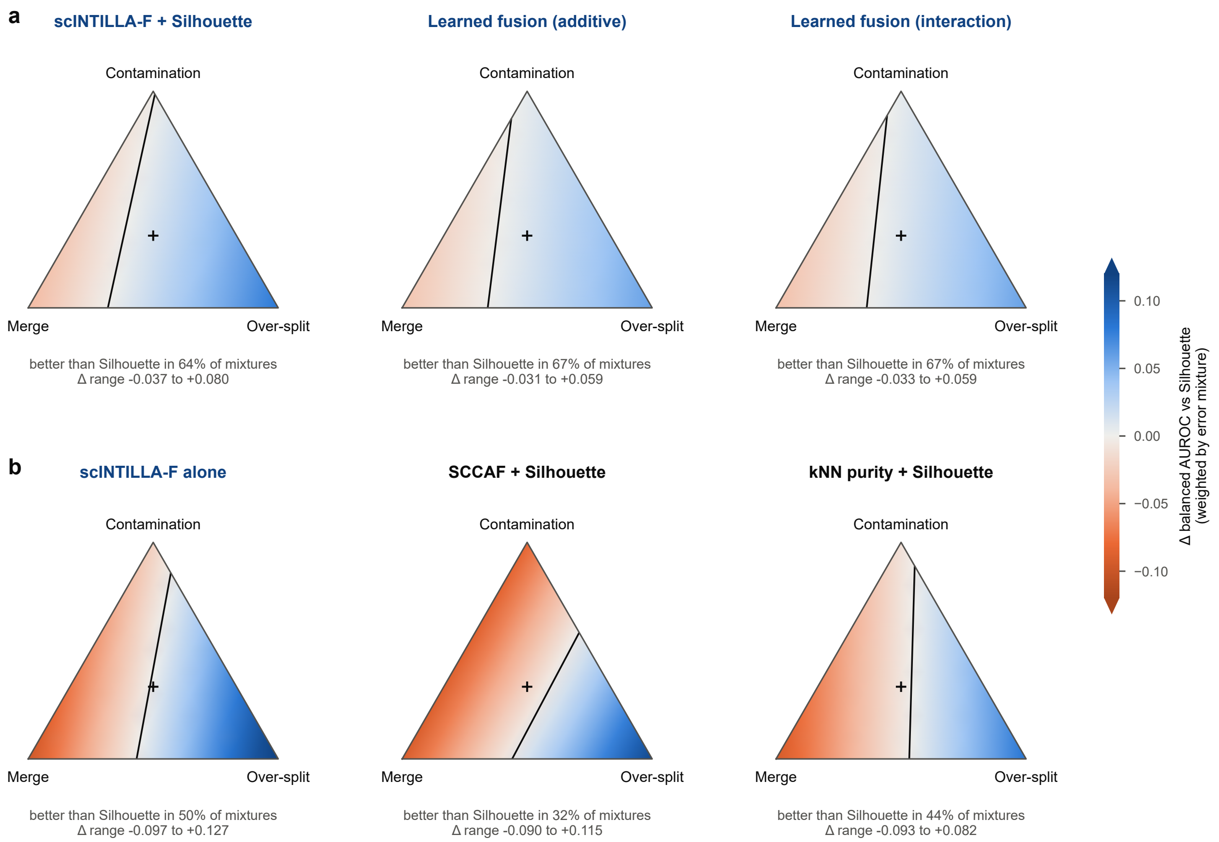


**Supplementary figure 6:** Advantage over the Silhouette baseline across mixtures of error types. Change in balanced AUROC relative to the Silhouette baseline for every mixture of merge, contamination and over-split weights on a 0.1 grid (66 mixtures); + marks equal weights and the black line marks no change. **(a)** scINTILLA-F + Silhouette (better in 64% of mixtures) and the two learned fusions (67%). **(b)** scINTILLA-F alone (50%), SCCAF + Silhouette (32%) and kNN purity + Silhouette (44%). The mixtures are not estimates of real error frequencies.

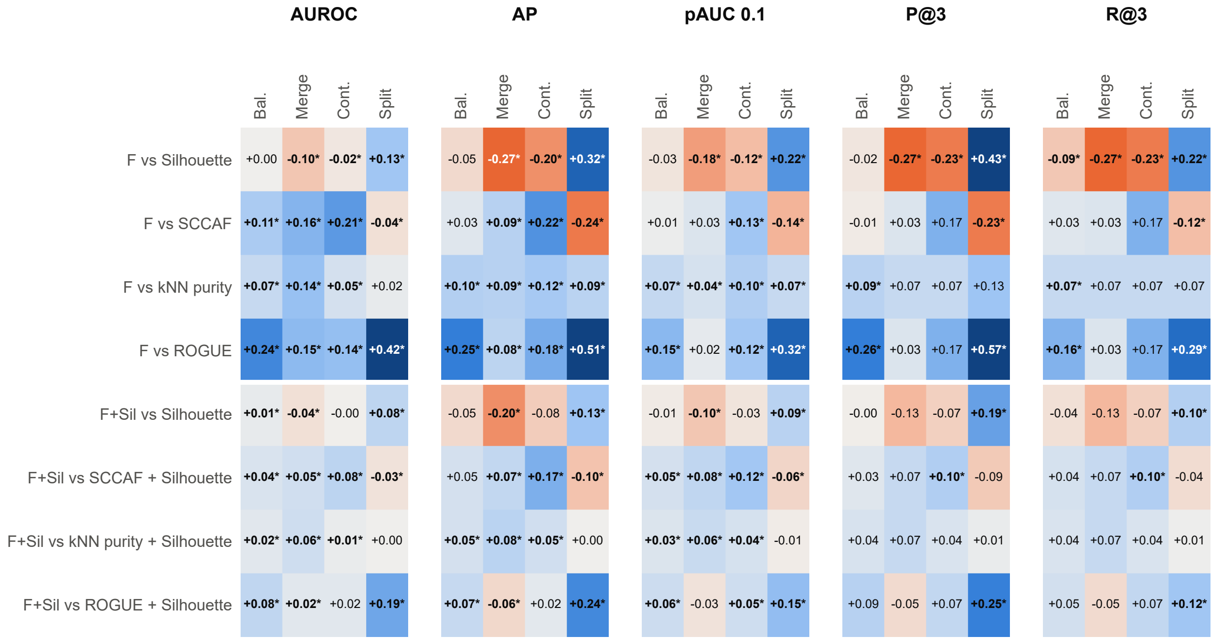


F = scINTILLA-F. Paired difference on matched labels (positive favours scINTILLA). * 95% atlas-bootstrap interval excludes 0 (unadjusted, 5 atlases; exploratory).

**Supplementary figure 7:** Paired effects of scINTILLA-F against every baseline, and of scINTILLA-F + Silhouette against every fusion built with the same rule, by error type and balanced; * marks a 95% atlas-bootstrap interval that excludes zero (unadjusted).

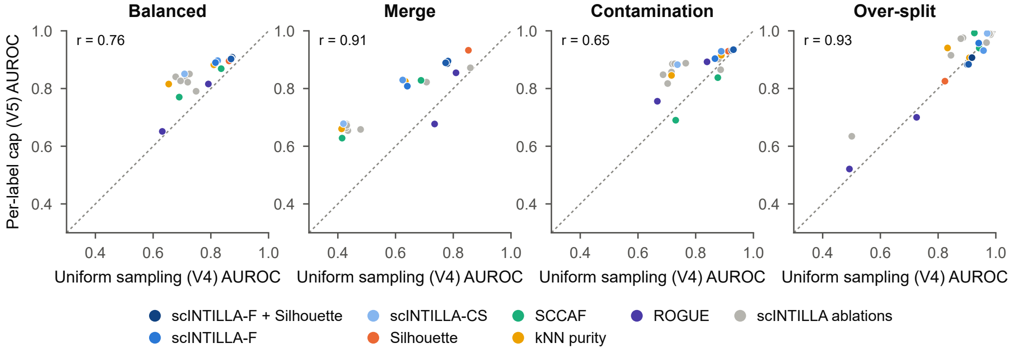


Each dot is one score; baseline + Silhouette fusions share their baseline colour. Different perturbation draws and cells, so the comparison is unpaired.

**Supplementary figure 8:** Sensitivity to the sampling design. Balanced AUROC of each method when 20,000 cells were sampled uniformly from each atlas before perturbation, against the main design in which every label is capped at 500 cells after perturbation. The two benchmarks use different cells and perturbation draws, so the comparison is unpaired.

**Supplementary table 6:** Leukaemia benchmark: every scINTILLA quantity against the annotated blast heterogeneity. All quantities are oriented so that higher is better (fragmentation, confusion and entropy are negated). $\rho$ label: Spearman correlation between the value of the patient label and the heterogeneity index. $\rho$ change: Spearman correlation between the change on splitting (blast-cluster labels minus patient label) and the index. Annotated and Shuffled: median change on splitting in patients with several blast clusters (68 AML, 19 B-ALL), with annotated and with shuffled blast-cluster labels. 99 AML and 84 B-ALL patients; fast preset, MrVI $z$ embedding, one seed. Composite scores (lower block) are min–max scaled or ranked within each run, and their changes are not comparable between runs.

| | AML | | | | B-ALL | | | |
|---|---|---|---|---|---|---|---|---|
| Quantity | $\rho$ label | $\rho$ change | Annotated | Shuffled | $\rho$ label | $\rho$ change | Annotated | Shuffled |
| Label fragmentation (mean) | −0.60 | 0.85 | 0.136 | 0.000 | −0.04 | 0.29 | 0.021 | 0.002 |
| Silhouette width | −0.65 | 0.75 | 0.180 | −0.021 | −0.15 | 0.18 | −0.031 | −0.037 |
| Clustering confusion (top clustering) | −0.44 | 0.02 | 0.033 | −0.161 | −0.10 | −0.33 | −0.019 | −0.048 |
| Clustering confusion (mean) | −0.51 | −0.24 | 0.005 | −0.195 | −0.08 | −0.52 | −0.053 | −0.082 |
| Prediction agreement | −0.50 | −0.58 | −0.018 | −0.148 | −0.02 | −0.71 | −0.037 | −0.054 |
| Prediction entropy | −0.50 | −0.57 | −0.048 | −0.394 | −0.02 | −0.71 | −0.096 | −0.142 |
| Prediction confidence | −0.50 | −0.67 | −0.040 | −0.222 | −0.03 | −0.73 | −0.062 | −0.105 |
| Confusion + fragmentation | −0.59 | 0.70 | 0.143 | −0.021 | −0.06 | −0.20 | 0.050 | 0.050 |
| Confusion only | −0.52 | 0.23 | 0.150 | −0.077 | −0.09 | −0.29 | 0.077 | 0.089 |
| Supervised only | −0.52 | 0.01 | 0.078 | −0.153 | −0.02 | −0.52 | −0.030 | −0.030 |
| scINTILLA-CS | −0.53 | 0.16 | 0.106 | −0.101 | −0.08 | −0.38 | −0.013 | −0.010 |
| scINTILLA-F | −0.59 | 0.56 | 0.117 | −0.064 | −0.06 | −0.34 | −0.005 | 0.000 |
| scINTILLA-CS + Silhouette | −0.61 | 0.64 | 0.198 | 0.089 | −0.10 | −0.17 | −0.049 | 0.110 |
| scINTILLA-F + Silhouette | −0.63 | 0.69 | 0.230 | 0.069 | −0.08 | −0.14 | −0.111 | 0.105 |
| Learned fusion (additive) | −0.65 | 0.68 | 1.712 | 0.564 | −0.13 | −0.03 | −0.776 | 0.722 |
| Learned fusion (interaction) | −0.65 | 0.70 | 1.507 | 0.545 | −0.14 | −0.04 | −0.760 | 0.665 |

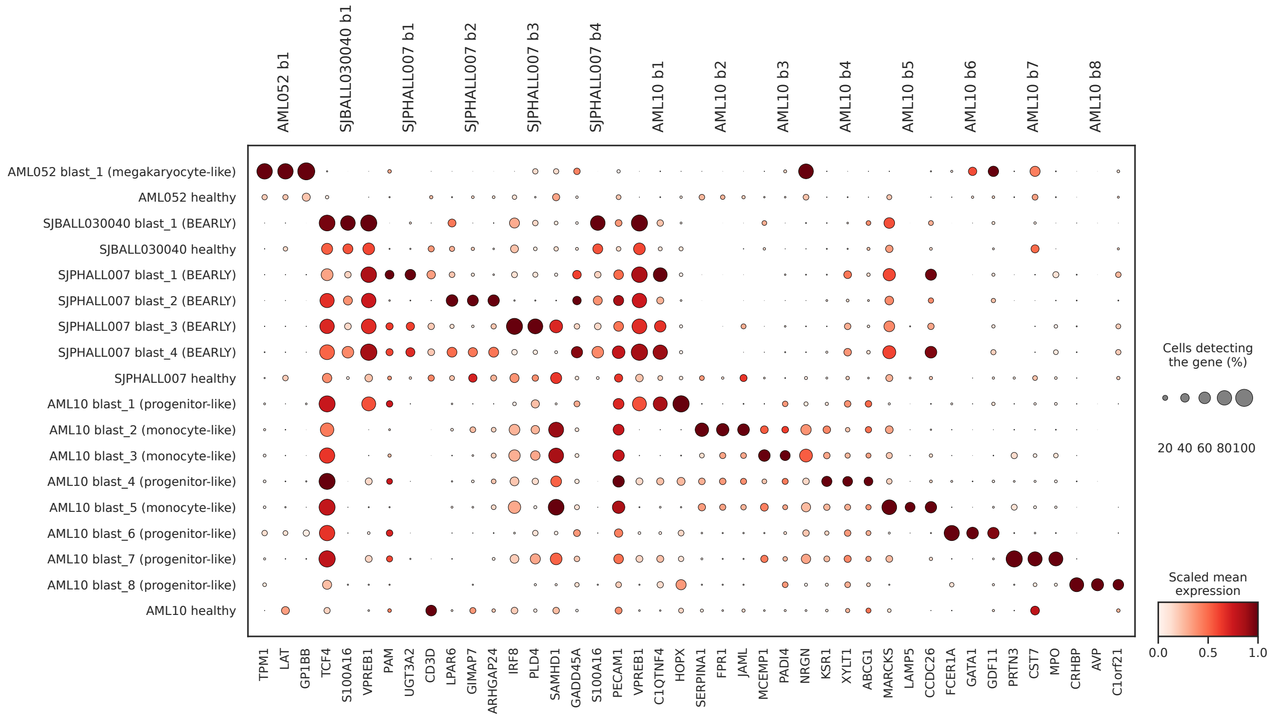


**Supplementary figure 9:** Marker genes of the annotated blast clusters in the four example patients of Fig. 5a. Rows are the blast clusters (b1, b2, . . . ) and the healthy cells of each patient, with the lineage assigned to each blast cluster in brackets (BEARLY, early B). Columns are the genes that distinguish each blast cluster, grouped by the cluster they mark. Dot size shows the percentage of cells in which the gene is detected and colour the scaled mean expression.

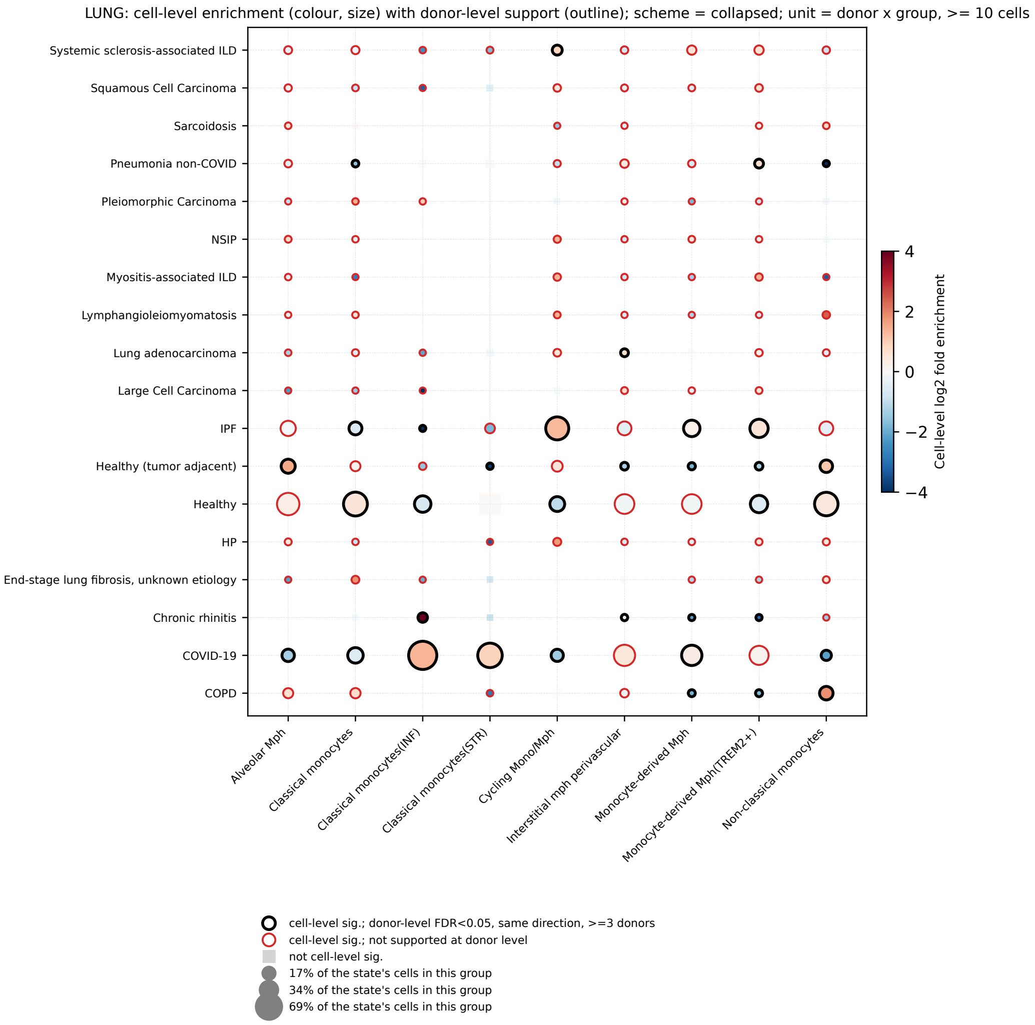


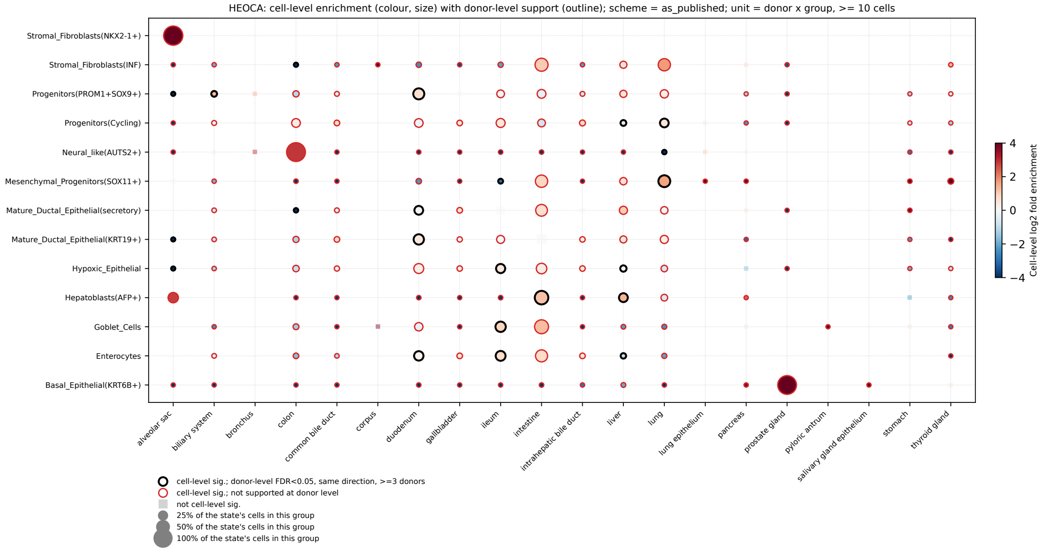


**Supplementary figure 10:** Enrichment of the refined states with donors as units, for the lung myeloid compartment (top) and the HEOCA stem-cell group (bottom). Layout as in Figs. 3e and 4e: colour shows the cell-level fold enrichment ($\log_2$) and dot size the share of the state's cells that come from the group. A black outline marks an enrichment supported at donor level, a red outline one that is significant at cell level only, and a grey square a pair that is not significant at cell level.

**Brain: medians over 10 replicates per error type**

| | AUROC | | | | Review utility | | | |
|---|---|---|---|---|---|---|---|---|
| Method | Merge | Contam. | Over-split | **Balanced** | AP | P@3 | R@3 | Coverage |
| **scINTILLA-F + Silhouette** | 0.954 | 0.993 | 0.991 | **0.979** | 0.840 | 0.833 | 0.667 | 100% |
| **scINTILLA learned fusion (additive)** | 0.950 | 0.986 | 0.978 | **0.971** | 0.837 | 0.778 | 0.611 | 100% |
| **scINTILLA learned fusion (interaction)** | 0.933 | 0.986 | 0.978 | **0.966** | 0.837 | 0.778 | 0.611 | 100% |
| **scINTILLA-CS + Silhouette** | 0.837 | 0.978 | 0.993 | **0.936** | 0.772 | 0.722 | 0.556 | 100% |
| **scINTILLA-F** | 0.908 | 0.993 | 1.000 | **0.967** | 0.852 | 0.833 | 0.667 | 100% |
| **scINTILLA-CS** | 0.667 | 0.971 | 1.000 | **0.879** | 0.698 | 0.667 | 0.500 | 100% |
| Confusion + fragmentation | 0.917 | 1.000 | 0.996 | **0.971** | 0.859 | 0.889 | 0.722 | 100% |
| Fragmentation only | 0.967 | 0.957 | 0.746 | **0.890** | 0.670 | 0.556 | 0.500 | 100% |
| Supervised consistency only | 0.617 | 0.870 | 1.000 | **0.829** | 0.560 | 0.444 | 0.278 | 100% |
| Confusion only | 0.683 | 1.000 | 1.000 | **0.894** | 0.757 | 0.667 | 0.500 | 100% |
| Confusion, 6-clustering mean | 0.683 | 1.000 | 1.000 | **0.894** | 0.757 | 0.667 | 0.500 | 100% |
| Confusion, best clustering | 0.725 | 1.000 | 1.000 | **0.908** | 0.761 | 0.778 | 0.611 | 100% |
| Classifier agreement | 0.608 | 0.862 | 0.996 | **0.822** | 0.552 | 0.444 | 0.278 | 100% |
| Classifier confidence | 0.667 | 0.862 | 1.000 | **0.843** | 0.557 | 0.444 | 0.278 | 100% |
| Classifier entropy (neg.) | 0.608 | 0.826 | 0.967 | **0.801** | 0.498 | 0.389 | 0.222 | 100% |
| Silhouette | 0.958 | 0.986 | 0.975 | **0.973** | 0.870 | 0.722 | 0.583 | 100% |
| SCCAF | 0.717 | 0.630 | 1.000 | **0.782** | 0.521 | 0.556 | 0.389 | 100% |
| kNN purity | 0.692 | 0.913 | 1.000 | **0.868** | 0.574 | 0.444 | 0.278 | 100% |
| ROGUE | 0.867 | 0.899 | 0.645 | **0.803** | 0.462 | 0.278 | 0.250 | 100% |
| SCCAF + Silhouette | 0.875 | 0.844 | 0.993 | **0.904** | 0.652 | 0.556 | 0.389 | 100% |
| kNN purity + Silhouette | 0.850 | 0.960 | 0.989 | **0.933** | 0.708 | 0.667 | 0.500 | 100% |
| ROGUE + Silhouette | 0.925 | 0.964 | 0.839 | **0.909** | 0.630 | 0.667 | 0.556 | 100% |

**Eye: medians over 10 replicates per error type**

| | AUROC | | | | Review utility | | | |
|---|---|---|---|---|---|---|---|---|
| Method | Merge | Contam. | Over-split | **Balanced** | AP | P@3 | R@3 | Coverage |
| **scINTILLA-F + Silhouette** | 0.912 | 0.947 | 0.944 | **0.934** | 0.299 | 0.139 | 0.125 | 100% |
| **scINTILLA learned fusion (additive)** | 0.910 | 0.965 | 0.935 | **0.937** | 0.341 | 0.111 | 0.111 | 100% |
| **scINTILLA learned fusion (interaction)** | 0.907 | 0.964 | 0.940 | **0.937** | 0.324 | 0.111 | 0.111 | 100% |
| **scINTILLA-CS + Silhouette** | 0.863 | 0.954 | 0.973 | **0.930** | 0.319 | 0.222 | 0.167 | 100% |
| **scINTILLA-F** | 0.840 | 0.929 | 0.973 | **0.914** | 0.285 | 0.333 | 0.222 | 100% |
| **scINTILLA-CS** | 0.688 | 0.899 | 0.992 | **0.860** | 0.338 | 0.222 | 0.111 | 100% |
| Confusion + fragmentation | 0.869 | 0.940 | 0.938 | **0.916** | 0.230 | 0.167 | 0.083 | 100% |
| Fragmentation only | 0.906 | 0.911 | 0.620 | **0.812** | 0.147 | 0.111 | 0.111 | 100% |
| Supervised consistency only | 0.704 | 0.832 | 0.997 | **0.844** | 0.376 | 0.333 | 0.167 | 100% |
| Confusion only | 0.613 | 0.924 | 0.989 | **0.842** | 0.307 | 0.222 | 0.111 | 100% |
| Confusion, 6-clustering mean | 0.614 | 0.924 | 0.989 | **0.842** | 0.307 | 0.222 | 0.111 | 100% |
| Confusion, best clustering | 0.617 | 0.922 | 0.987 | **0.842** | 0.299 | 0.222 | 0.111 | 100% |
| Classifier agreement | 0.690 | 0.857 | 0.986 | **0.844** | 0.363 | 0.389 | 0.222 | 100% |
| Classifier confidence | 0.694 | 0.861 | 1.000 | **0.852** | 0.392 | 0.333 | 0.167 | 100% |
| Classifier entropy (neg.) | 0.667 | 0.790 | 0.940 | **0.799** | 0.254 | 0.222 | 0.111 | 100% |
| Silhouette | 0.947 | 0.964 | 0.886 | **0.932** | 0.440 | 0.222 | 0.222 | 100% |
| SCCAF | 0.596 | 0.717 | 1.000 | **0.771** | 0.379 | 0.333 | 0.167 | 100% |
| kNN purity | 0.607 | 0.900 | 0.975 | **0.827** | 0.242 | 0.222 | 0.111 | 100% |
| ROGUE | 0.867 | 0.909 | 0.490 | **0.755** | 0.122 | 0.000 | 0.000 | 94% |
| SCCAF + Silhouette | 0.833 | 0.887 | 0.971 | **0.897** | 0.337 | 0.333 | 0.222 | 100% |
| kNN purity + Silhouette | 0.811 | 0.953 | 0.964 | **0.909** | 0.293 | 0.250 | 0.181 | 100% |
| ROGUE + Silhouette | 0.937 | 0.941 | 0.740 | **0.872** | 0.303 | 0.222 | 0.222 | 94% |

**HEOCA: medians over 10 replicates per error type**

| | AUROC | | | | Review utility | | | |
|---|---|---|---|---|---|---|---|---|
| Method | Merge | Contam. | Over-split | **Balanced** | AP | P@3 | R@3 | Coverage |
| **scINTILLA-F + Silhouette** | 0.843 | 0.919 | 0.886 | **0.883** | 0.564 | 0.528 | 0.431 | 100% |
| **scINTILLA learned fusion (additive)** | 0.873 | 0.910 | 0.833 | **0.872** | 0.603 | 0.556 | 0.500 | 100% |
| **scINTILLA learned fusion (interaction)** | 0.868 | 0.914 | 0.833 | **0.872** | 0.595 | 0.500 | 0.444 | 100% |
| **scINTILLA-CS + Silhouette** | 0.777 | 0.919 | 0.912 | **0.869** | 0.579 | 0.537 | 0.426 | 100% |
| **scINTILLA-F** | 0.730 | 0.860 | 0.973 | **0.855** | 0.500 | 0.389 | 0.222 | 100% |
| **scINTILLA-CS** | 0.657 | 0.838 | 1.000 | **0.832** | 0.480 | 0.333 | 0.167 | 100% |
| Confusion + fragmentation | 0.735 | 0.869 | 0.923 | **0.843** | 0.401 | 0.278 | 0.167 | 100% |
| Fragmentation only | 0.784 | 0.820 | 0.579 | **0.728** | 0.304 | 0.222 | 0.222 | 100% |
| Supervised consistency only | 0.667 | 0.860 | 1.000 | **0.842** | 0.498 | 0.389 | 0.222 | 100% |
| Confusion only | 0.657 | 0.833 | 0.977 | **0.823** | 0.446 | 0.222 | 0.111 | 100% |
| Confusion, 6-clustering mean | 0.657 | 0.833 | 0.977 | **0.823** | 0.446 | 0.222 | 0.111 | 100% |
| Confusion, best clustering | 0.637 | 0.829 | 0.937 | **0.801** | 0.402 | 0.222 | 0.111 | 100% |
| Classifier agreement | 0.689 | 0.928 | 0.993 | **0.870** | 0.639 | 0.611 | 0.444 | 100% |
| Classifier confidence | 0.691 | 0.811 | 1.000 | **0.834** | 0.467 | 0.333 | 0.167 | 100% |
| Classifier entropy (neg.) | 0.681 | 0.878 | 0.984 | **0.848** | 0.504 | 0.500 | 0.333 | 100% |
| Silhouette | 0.902 | 0.896 | 0.761 | **0.853** | 0.511 | 0.444 | 0.444 | 100% |
| SCCAF | 0.598 | 0.680 | 1.000 | **0.759** | 0.437 | 0.333 | 0.167 | 100% |
| kNN purity | 0.652 | 0.802 | 0.948 | **0.801** | 0.377 | 0.222 | 0.111 | 100% |
| ROGUE | 0.515 | 0.622 | 0.466 | **0.534** | 0.143 | 0.000 | 0.000 | 100% |
| SCCAF + Silhouette | 0.782 | 0.806 | 0.943 | **0.844** | 0.502 | 0.468 | 0.356 | 100% |
| kNN purity + Silhouette | 0.814 | 0.912 | 0.882 | **0.869** | 0.486 | 0.500 | 0.389 | 100% |
| ROGUE + Silhouette | 0.745 | 0.806 | 0.663 | **0.738** | 0.379 | 0.306 | 0.264 | 100% |

**HNOCA: medians over 10 replicates per error type**

| | AUROC | | | | Review utility | | | |
|---|---|---|---|---|---|---|---|---|
| Method | Merge | Contam. | Over-split | **Balanced** | AP | P@3 | R@3 | Coverage |
| **scINTILLA-F + Silhouette** | 0.781 | 0.823 | 0.827 | **0.810** | 0.210 | 0.111 | 0.056 | 100% |
| **scINTILLA learned fusion (additive)** | 0.805 | 0.824 | 0.815 | **0.815** | 0.212 | 0.000 | 0.000 | 100% |
| **scINTILLA learned fusion (interaction)** | 0.802 | 0.827 | 0.814 | **0.814** | 0.210 | 0.056 | 0.028 | 100% |
| **scINTILLA-CS + Silhouette** | 0.764 | 0.817 | 0.868 | **0.816** | 0.245 | 0.222 | 0.167 | 100% |
| **scINTILLA-F** | 0.689 | 0.798 | 0.896 | **0.794** | 0.239 | 0.111 | 0.056 | 100% |
| **scINTILLA-CS** | 0.626 | 0.771 | 0.966 | **0.787** | 0.349 | 0.278 | 0.167 | 100% |
| Confusion + fragmentation | 0.704 | 0.786 | 0.854 | **0.781** | 0.202 | 0.056 | 0.028 | 100% |
| Fragmentation only | 0.748 | 0.726 | 0.615 | **0.696** | 0.152 | 0.000 | 0.000 | 100% |
| Supervised consistency only | 0.623 | 0.765 | 0.951 | **0.779** | 0.301 | 0.278 | 0.167 | 100% |
| Confusion only | 0.597 | 0.765 | 0.966 | **0.776** | 0.346 | 0.278 | 0.167 | 100% |
| Confusion, 6-clustering mean | 0.597 | 0.768 | 0.964 | **0.777** | 0.351 | 0.278 | 0.167 | 100% |
| Confusion, best clustering | 0.601 | 0.768 | 0.972 | **0.780** | 0.377 | 0.333 | 0.222 | 100% |
| Classifier agreement | 0.610 | 0.807 | 0.958 | **0.792** | 0.334 | 0.278 | 0.167 | 100% |
| Classifier confidence | 0.588 | 0.658 | 0.952 | **0.733** | 0.299 | 0.222 | 0.111 | 100% |
| Classifier entropy (neg.) | 0.610 | 0.786 | 0.921 | **0.772** | 0.304 | 0.278 | 0.167 | 100% |
| Silhouette | 0.874 | 0.807 | 0.735 | **0.805** | 0.249 | 0.000 | 0.000 | 100% |
| SCCAF | 0.566 | 0.646 | 0.963 | **0.725** | 0.292 | 0.222 | 0.111 | 100% |
| kNN purity | 0.610 | 0.756 | 0.894 | **0.753** | 0.198 | 0.000 | 0.000 | 100% |
| ROGUE | 0.557 | 0.673 | 0.557 | **0.595** | 0.110 | 0.000 | 0.000 | 100% |
| SCCAF + Silhouette | 0.764 | 0.726 | 0.874 | **0.788** | 0.209 | 0.111 | 0.056 | 100% |
| kNN purity + Silhouette | 0.759 | 0.799 | 0.847 | **0.802** | 0.219 | 0.139 | 0.083 | 100% |
| ROGUE + Silhouette | 0.800 | 0.835 | 0.649 | **0.761** | 0.181 | 0.000 | 0.000 | 100% |

**Lung: medians over 10 replicates per error type**

| | AUROC | | | | Review utility | | | |
|---|---|---|---|---|---|---|---|---|
| Method | Merge | Contam. | Over-split | **Balanced** | AP | P@3 | R@3 | Coverage |
| **scINTILLA-F + Silhouette** | 0.941 | 0.978 | 0.888 | **0.936** | 0.478 | 0.417 | 0.361 | 100% |
| **scINTILLA learned fusion (additive)** | 0.938 | 0.985 | 0.852 | **0.925** | 0.496 | 0.333 | 0.333 | 100% |
| **scINTILLA learned fusion (interaction)** | 0.935 | 0.982 | 0.854 | **0.924** | 0.482 | 0.333 | 0.333 | 100% |
| **scINTILLA-CS + Silhouette** | 0.906 | 0.977 | 0.913 | **0.932** | 0.469 | 0.426 | 0.370 | 100% |
| **scINTILLA-F** | 0.873 | 0.939 | 0.946 | **0.919** | 0.433 | 0.278 | 0.194 | 100% |
| **scINTILLA-CS** | 0.753 | 0.936 | 0.999 | **0.896** | 0.530 | 0.444 | 0.278 | 100% |
| Confusion + fragmentation | 0.886 | 0.915 | 0.864 | **0.888** | 0.326 | 0.167 | 0.139 | 100% |
| Fragmentation only | 0.954 | 0.912 | 0.613 | **0.826** | 0.354 | 0.278 | 0.278 | 100% |
| Supervised consistency only | 0.765 | 0.959 | 1.000 | **0.908** | 0.603 | 0.500 | 0.333 | 100% |
| Confusion only | 0.719 | 0.904 | 0.949 | **0.857** | 0.336 | 0.222 | 0.167 | 100% |
| Confusion, 6-clustering mean | 0.719 | 0.901 | 0.949 | **0.856** | 0.330 | 0.222 | 0.167 | 100% |
| Confusion, best clustering | 0.738 | 0.909 | 0.968 | **0.872** | 0.411 | 0.333 | 0.222 | 100% |
| Classifier agreement | 0.787 | 0.985 | 1.000 | **0.924** | 0.668 | 0.556 | 0.389 | 100% |
| Classifier confidence | 0.651 | 0.895 | 1.000 | **0.849** | 0.468 | 0.444 | 0.278 | 100% |
| Classifier entropy (neg.) | 0.796 | 0.959 | 0.984 | **0.913** | 0.556 | 0.444 | 0.278 | 100% |
| Silhouette | 0.981 | 0.994 | 0.770 | **0.915** | 0.637 | 0.444 | 0.444 | 100% |
| SCCAF | 0.664 | 0.778 | 1.000 | **0.814** | 0.450 | 0.389 | 0.222 | 100% |
| kNN purity | 0.741 | 0.857 | 0.886 | **0.828** | 0.226 | 0.056 | 0.056 | 100% |
| ROGUE | 0.580 | 0.678 | 0.447 | **0.569** | 0.125 | 0.000 | 0.000 | 100% |
| SCCAF + Silhouette | 0.887 | 0.925 | 0.921 | **0.911** | 0.396 | 0.435 | 0.356 | 100% |
| kNN purity + Silhouette | 0.890 | 0.953 | 0.851 | **0.898** | 0.343 | 0.222 | 0.222 | 100% |
| ROGUE + Silhouette | 0.866 | 0.917 | 0.610 | **0.798** | 0.420 | 0.333 | 0.306 | 100% |

**Supplementary figure 11:** Benchmark results per atlas (brain, eye, HEOCA, HNOCA and lung) for every method, component and fusion: AUROC by error type and balanced, average precision (AP), precision and recall among the three lowest-scoring labels (P@3, R@3) and coverage; medians over the ten replicates of each atlas and error type.

Balanced over error types and atlases. Colour scaled within each column (darker = better); values printed. Partial-coverage rows: ranking metrics on scored labels only.

| | AUROC | AP | Adj. AP | pAUC 0.1 | pAUC 0.2 | P@1 | R@1 | P@3 | R@3 | E@3 | P@10% | R@10% | Coverage |
|---|---|---|---|---|---|---|---|---|---|---|---|---|---|
| **scINTILLA-F + Silhouette** | 0.91 | 0.48 | 0.44 | 0.71 | 0.78 | 0.42 | 0.12 | 0.41 | 0.33 | 4.98 | 0.40 | 0.55 | 1.00 |
| **scINTILLA learned fusion (additive)** | 0.90 | 0.50 | 0.46 | 0.70 | 0.77 | 0.57 | 0.17 | 0.36 | 0.31 | 4.57 | 0.39 | 0.55 | 1.00 |
| **scINTILLA learned fusion (interaction)** | 0.90 | 0.49 | 0.45 | 0.70 | 0.77 | 0.50 | 0.14 | 0.36 | 0.31 | 4.55 | 0.39 | 0.53 | 1.00 |
| **scINTILLA-CS + Silhouette** | 0.90 | 0.48 | 0.44 | 0.70 | 0.77 | 0.38 | 0.10 | 0.43 | 0.34 | 5.61 | 0.37 | 0.51 | 1.00 |
| **scINTILLA-F** | 0.89 | 0.46 | 0.43 | 0.67 | 0.76 | 0.30 | 0.07 | 0.39 | 0.27 | 4.63 | 0.38 | 0.48 | 1.00 |
| **scINTILLA-CS** | 0.85 | 0.48 | 0.44 | 0.68 | 0.74 | 0.43 | 0.09 | 0.39 | 0.24 | 4.11 | 0.38 | 0.44 | 1.00 |
| Silhouette | 0.90 | 0.54 | 0.50 | 0.72 | 0.77 | 0.60 | 0.19 | 0.37 | 0.34 | 5.67 | 0.35 | 0.52 | 1.00 |
| SCCAF | 0.77 | 0.42 | 0.37 | 0.65 | 0.67 | 0.33 | 0.06 | 0.37 | 0.21 | 3.94 | 0.32 | 0.36 | 1.00 |
| kNN purity | 0.82 | 0.32 | 0.27 | 0.58 | 0.66 | 0.13 | 0.02 | 0.19 | 0.11 | 1.99 | 0.20 | 0.24 | 1.00 |
| ROGUE | 0.65 | 0.19 | 0.12 | 0.51 | 0.55 | 0.10 | 0.03 | 0.06 | 0.05 | 0.42 | 0.09 | 0.14 | 0.99 |
| SCCAF + Silhouette | 0.87 | 0.42 | 0.38 | 0.65 | 0.72 | 0.31 | 0.07 | 0.38 | 0.28 | 5.14 | 0.33 | 0.43 | 1.00 |
| kNN purity + Silhouette | 0.88 | 0.41 | 0.37 | 0.65 | 0.73 | 0.22 | 0.05 | 0.36 | 0.27 | 4.61 | 0.34 | 0.45 | 1.00 |
| ROGUE + Silhouette | 0.82 | 0.38 | 0.33 | 0.63 | 0.68 | 0.40 | 0.13 | 0.31 | 0.27 | 4.64 | 0.27 | 0.38 | 0.99 |
| Confusion + fragmentation | 0.88 | 0.40 | 0.36 | 0.64 | 0.73 | 0.20 | 0.04 | 0.31 | 0.23 | 3.07 | 0.36 | 0.47 | 1.00 |
| Fragmentation only | 0.79 | 0.33 | 0.27 | 0.59 | 0.66 | 0.17 | 0.06 | 0.23 | 0.22 | 3.39 | 0.23 | 0.33 | 1.00 |
| Supervised consistency only | 0.84 | 0.47 | 0.43 | 0.67 | 0.71 | 0.40 | 0.08 | 0.39 | 0.23 | 4.57 | 0.32 | 0.37 | 1.00 |

**Supplementary figure 12:** All review endpoints, balanced over error types and atlases: AUROC, AP, adjusted AP, standardised partial AUROC at false-positive rates 0.1 and 0.2, precision (P), recall (R) and enrichment (E) among the lowest-scoring 1 and 3 labels and the lowest 10% of labels, and coverage. Colour is scaled within each column (darker = better). Rows that do not score every label (ROGUE) are computed on the labels they score.

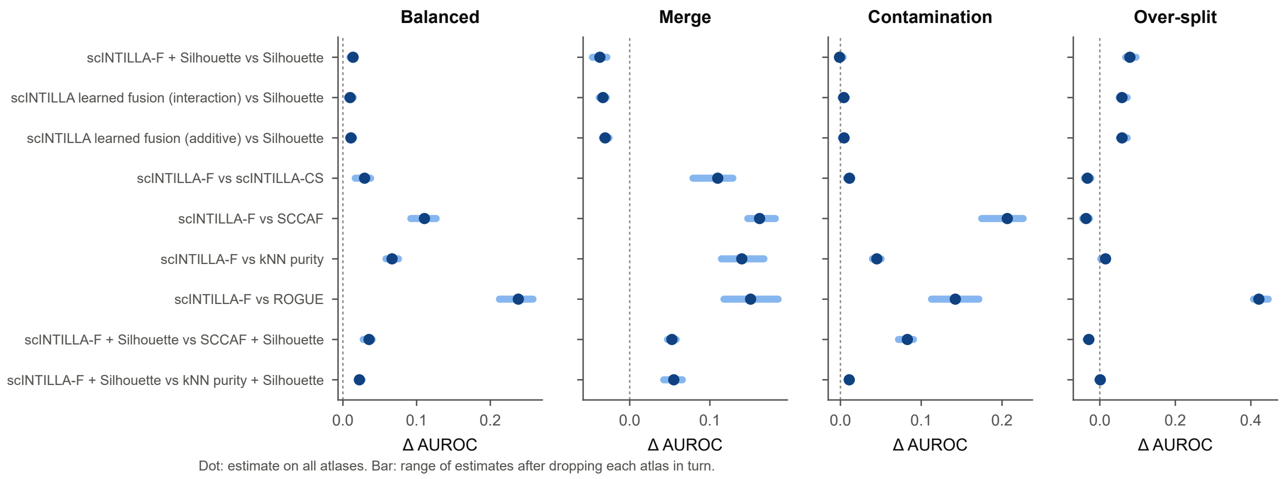


**Supplementary figure 13:** Leave-one-atlas-out stability of the key paired effects (difference in AUROC, balanced and by error type). Dots show the estimate on all five atlases and bars the range of estimates obtained when each atlas is dropped in turn.

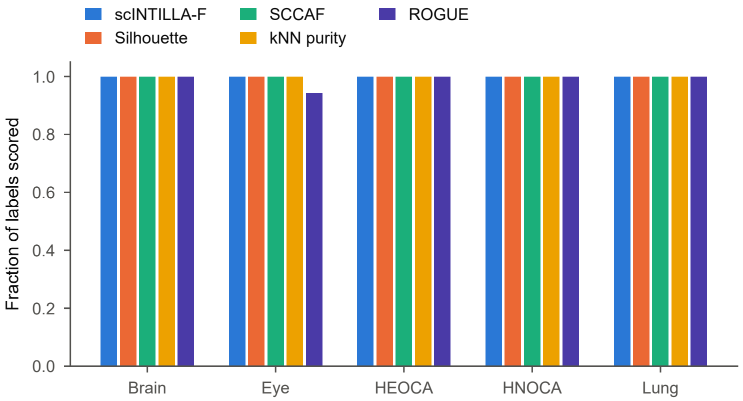


**Supplementary figure 14:** Label coverage per atlas: the fraction of labels that receive a score. Every scINTILLA variant, Silhouette, SCCAF and kNN purity score every label; ROGUE abstains on labels with too few cells per donor (99% coverage overall).

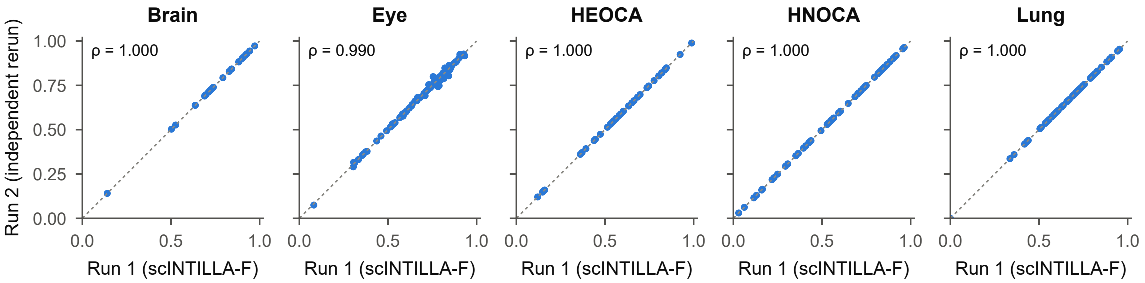


**Supplementary figure 15:** Run-to-run reproducibility of scINTILLA-F on the original annotations: per-label scores from two independent runs with the benchmarked configuration (Spearman $\rho = 1.000$ for the brain, HEOCA, HNOCA and lung atlases and 0.990 for the eye atlas).

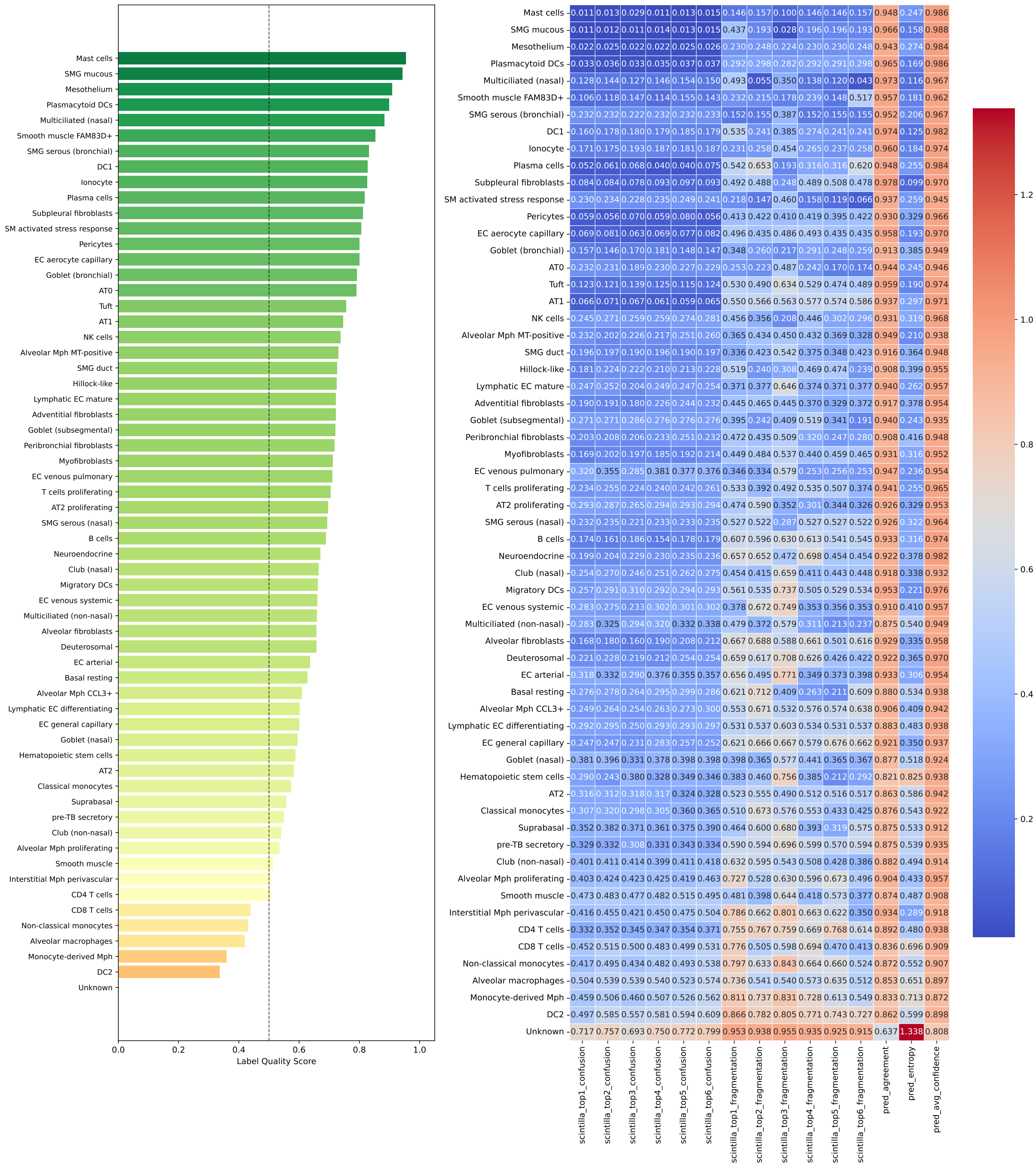


**Supplementary figure 16:** Per-cell-type scINTILLA-F scores for the human lung atlas (61 labels). Scored with the benchmarked configuration (at most 500 cells per label, label-free PCA). Left: label quality scores ranked from highest to lowest. Right: heatmap of the per-label means of the underlying metrics (clustering confusion and label fragmentation for the top six clusterings, prediction agreement, prediction entropy and average confidence).

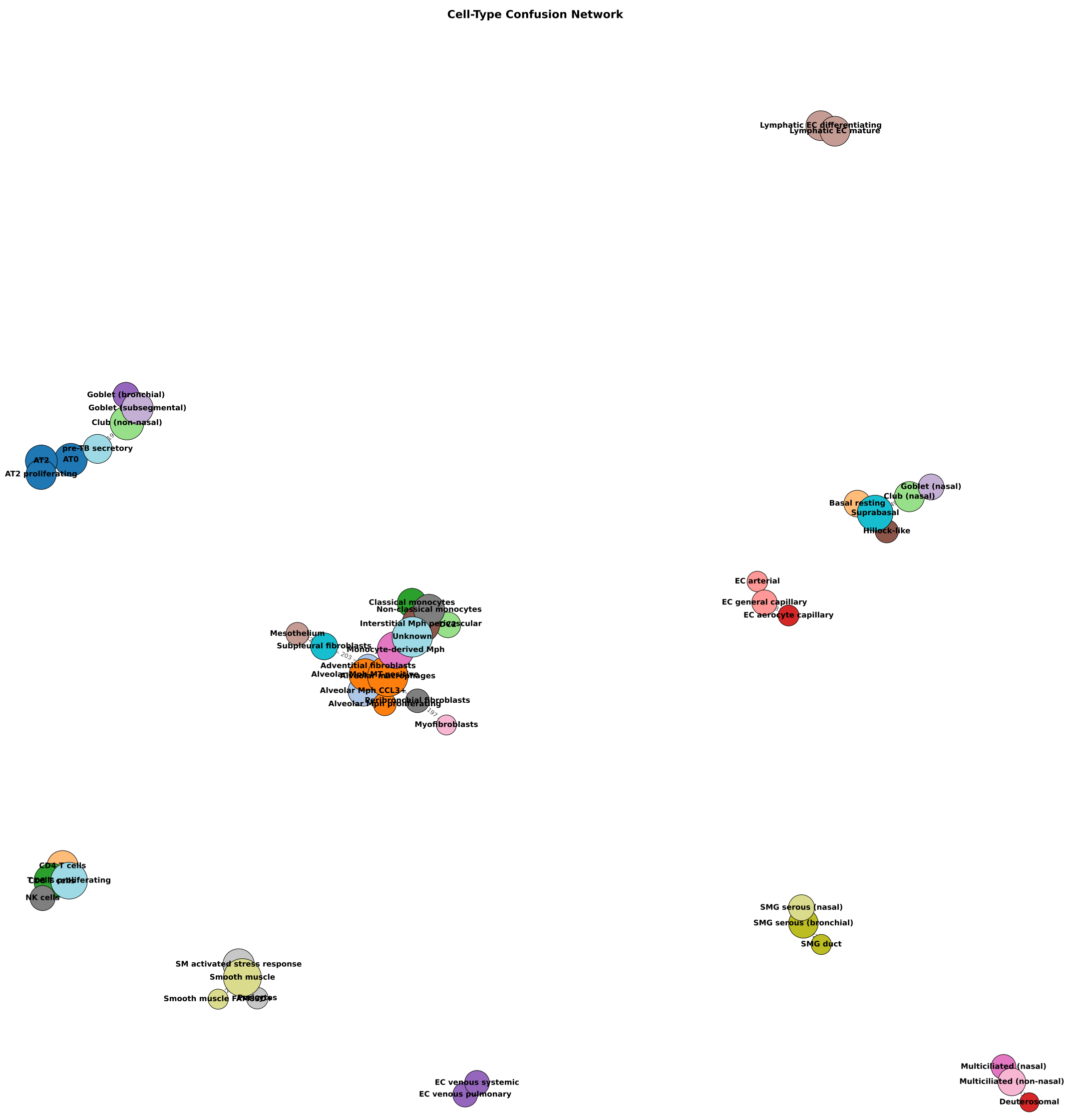


**Supplementary figure 17:** Confusion network for the human lung atlas. Nodes are annotated cell types; two labels are joined when the number of classifier predictions exchanged between them exceeds a threshold, and edge width reflects the amount of confusion. The monocyte, macrophage and “Unknown” labels form one connected module.

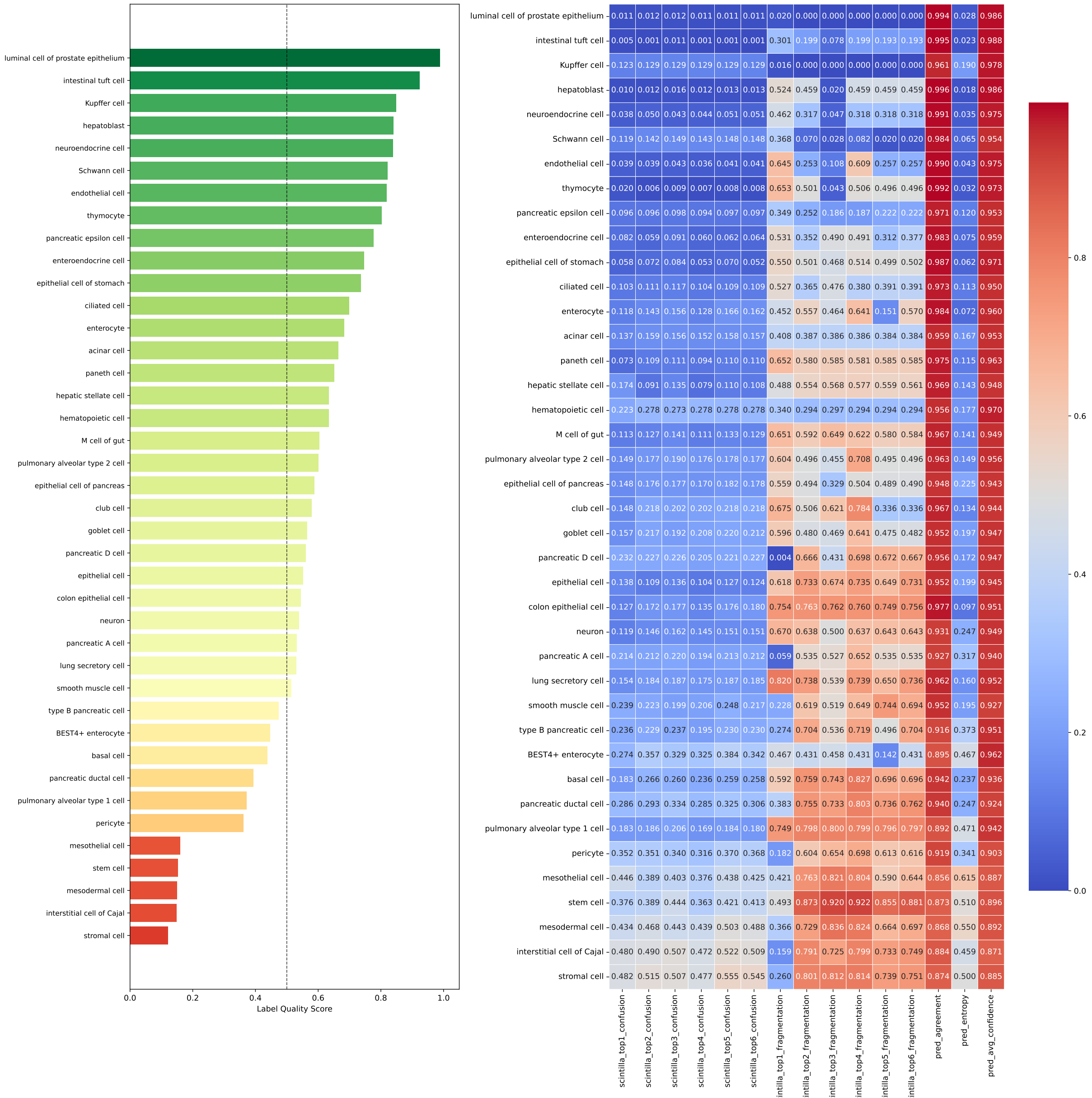


**Supplementary figure 18:** Per-cell-type scINTILLA-F scores for the HEOCA atlas (40 labels). Scored with the benchmarked configuration (at most 500 cells per label, label-free PCA). Left: label quality scores ranked from highest to lowest. Right: heatmap of the per-label means of the underlying metrics (clustering confusion and label fragmentation for the top six clusterings, prediction agreement, prediction entropy and average confidence). Stromal cells, interstitial cells of Cajal, mesodermal cells and stem cells score lowest.

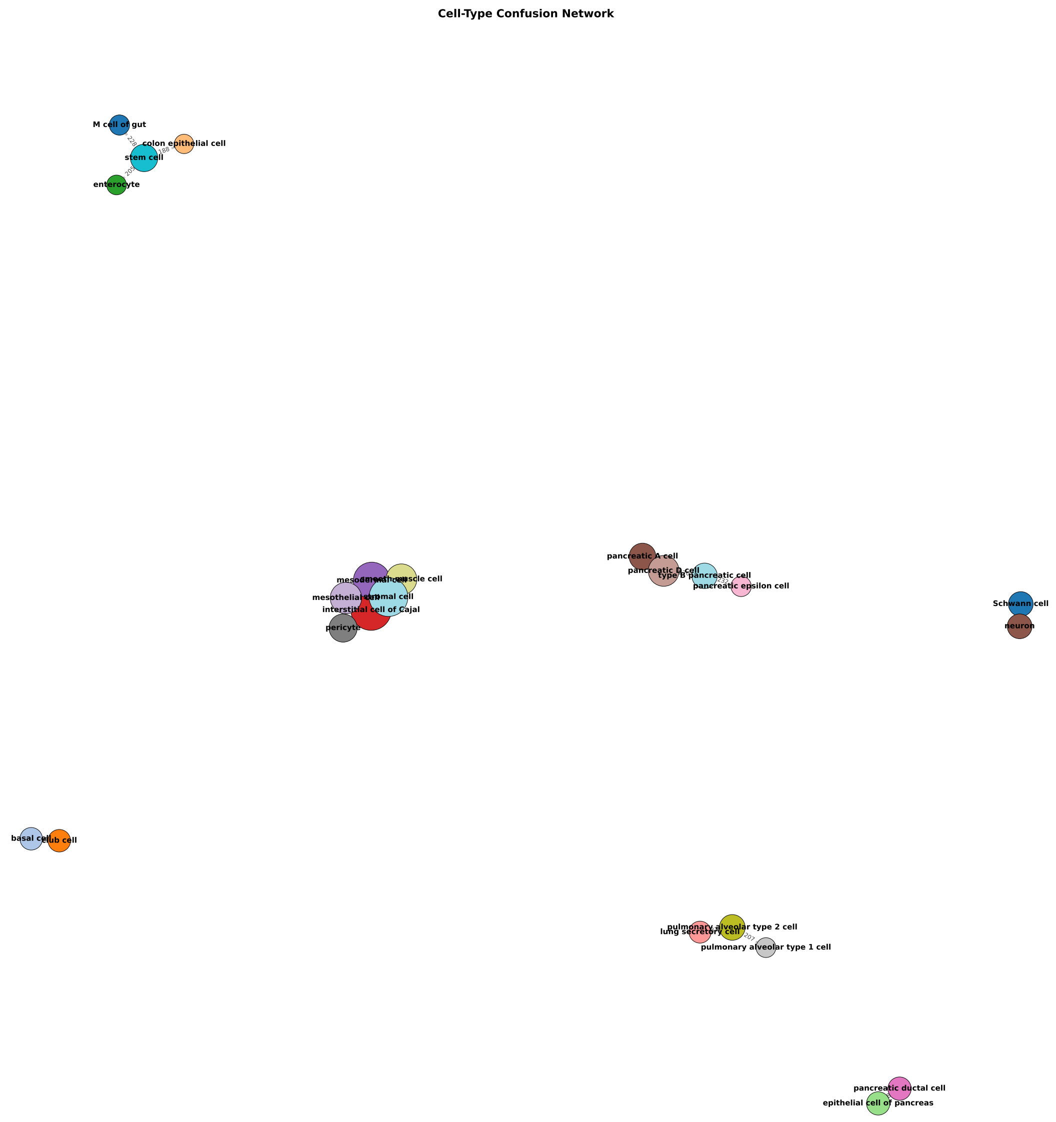


**Supplementary figure 19:** Confusion network for the HEOCA atlas. Nodes are annotated cell types; two labels are joined when the number of classifier predictions exchanged between them exceeds a threshold, and edge width reflects the amount of confusion. A mesenchymal and stromal module (mesodermal, mesothelial, stromal and smooth muscle cells, pericytes and interstitial cells of Cajal) is separate from a gut epithelial module in which the stem cell label is connected to enterocytes, M cells and colon epithelial cells.

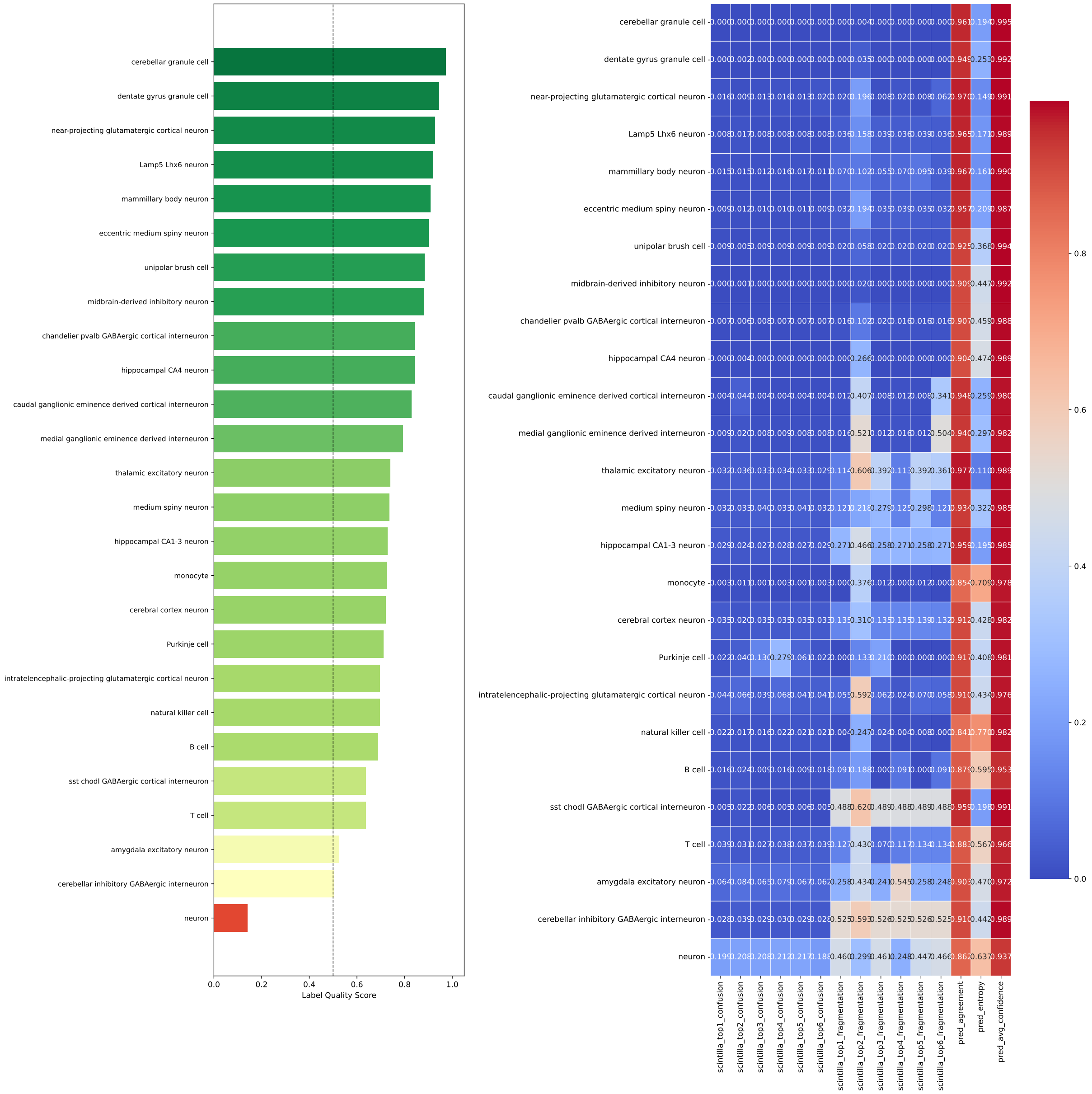


**Supplementary figure 20:** Per-cell-type scINTILLA-F scores for the adult human brain atlas (26 labels). Scored with the benchmarked configuration (at most 500 cells per label, label-free PCA). Left: label quality scores ranked from highest to lowest. Right: heatmap of the per-label means of the underlying metrics (clustering confusion and label fragmentation for the top six clusterings, prediction agreement, prediction entropy and average confidence). The broad “neuron” label ranks lowest, followed by cerebellar inhibitory GABAergic interneurons and amygdala excitatory neurons.

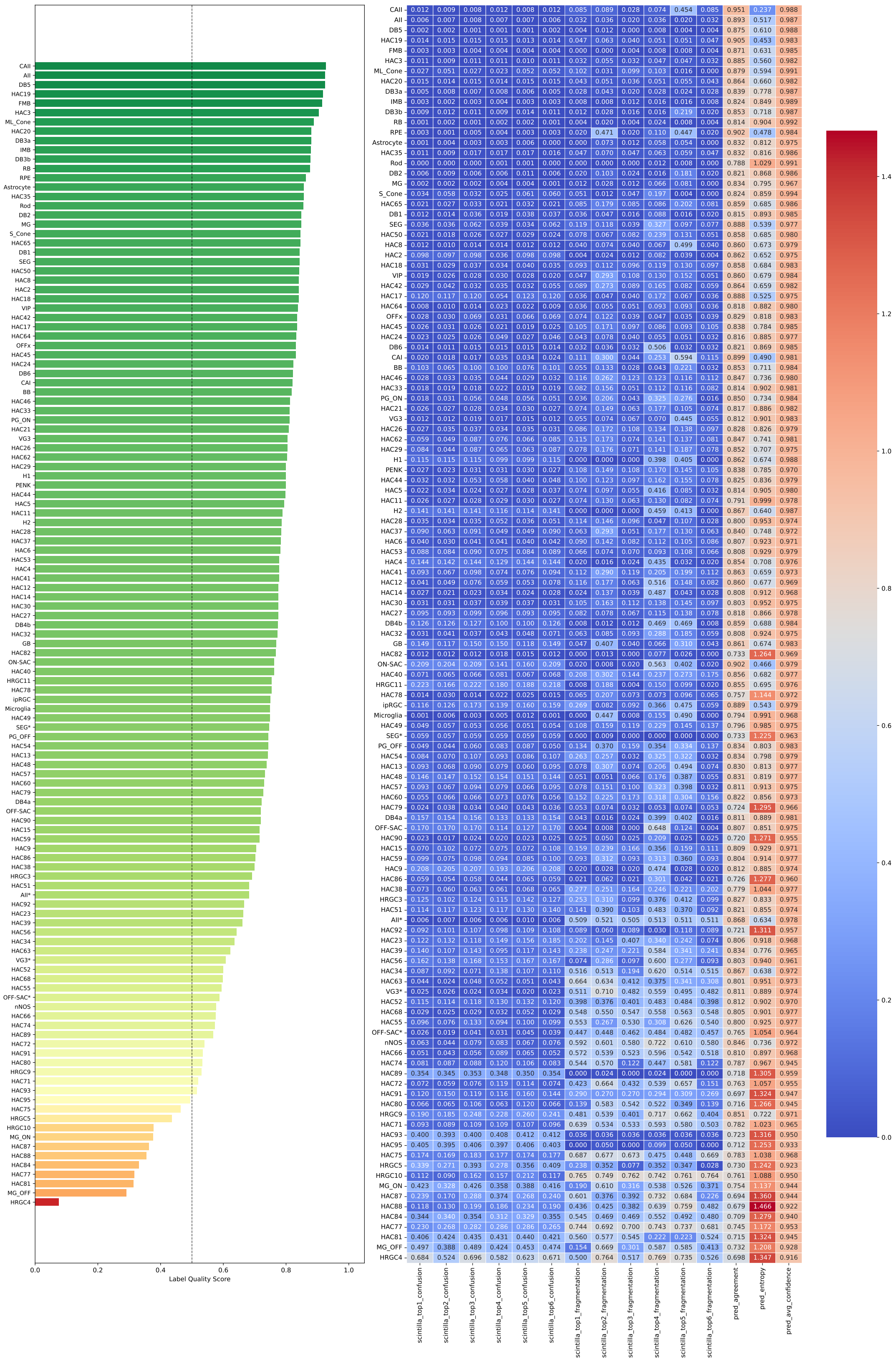


**Supplementary figure 21:** Per-cell-type scINTILLA-F scores for the human eye atlas (123 labels). Scored with the benchmarked configuration (at most 500 cells per label, label-free PCA). Label quality scores ranked from highest to lowest, with the corresponding heatmap of the underlying metrics.

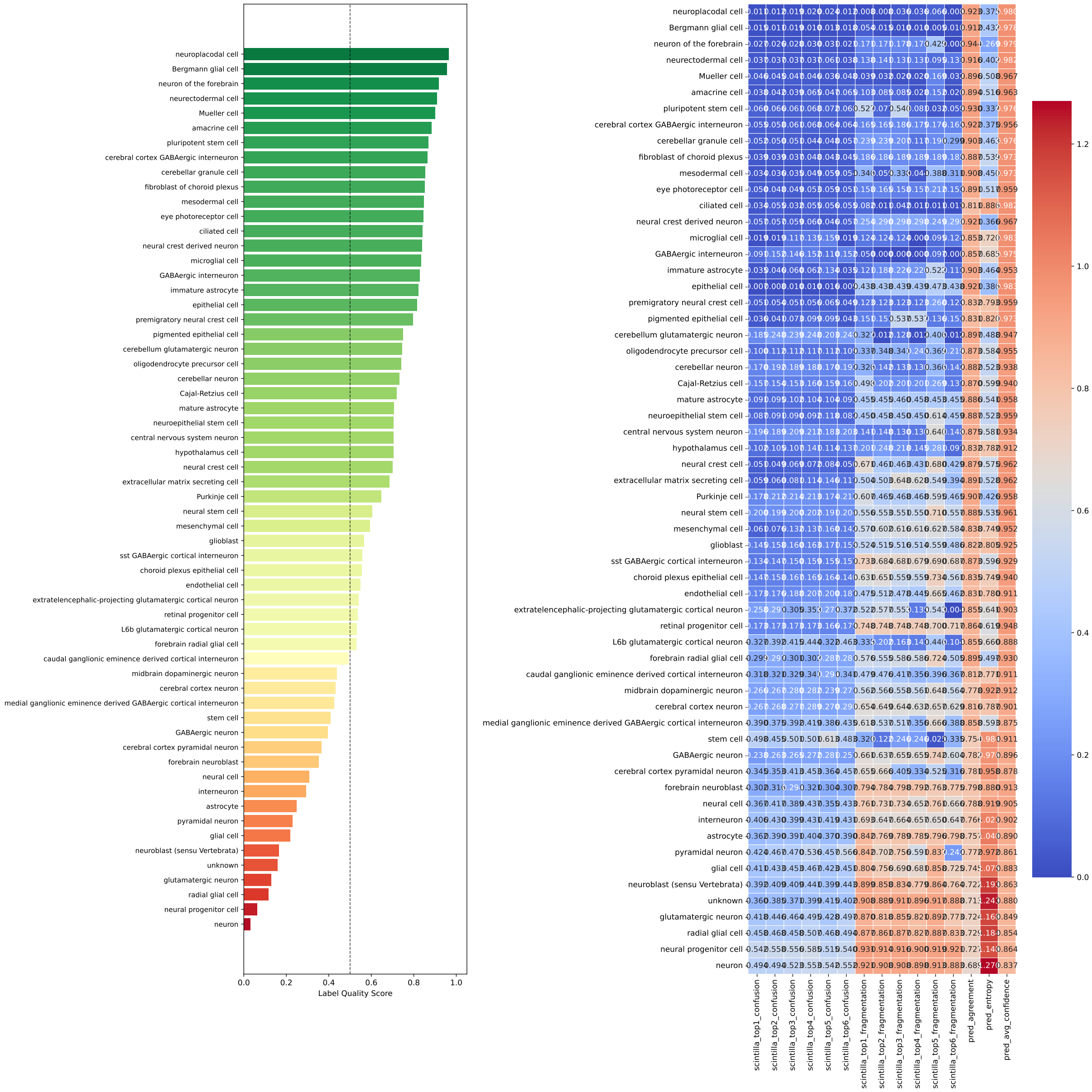


**Supplementary figure 22:** Per-cell-type scINTILLA-F scores for the Human Neural Organoid Cell Atlas (HNOCA; 60 labels). Scored with the benchmarked configuration (at most 500 cells per label, label-free PCA). Label quality scores ranked from highest to lowest, with the corresponding heatmap of the underlying metrics.
